# Dynamic anatomy of the replisome under stress

**DOI:** 10.64898/2026.06.01.729259

**Authors:** Laura Schulte, Maruthi K. Pabba, Maliwan Kamkaew, Paulina Prorok, Maria Arroyo, M. Cristina Cardoso

**Author notes:** These authors contributed equally to this work (Laura Schulte, Maruthi K. Pabba). Corresponding authors, Further information and requests for resources and reagents should be directed to and will be fulfilled by the lead contact, M. Cristina Cardoso.

## Abstract

When exposed to replication stress, the factors involved in DNA replication display differing responses that in many cases remain unexplained. In this study, we generated a series of genomically engineered embryonic stem cell lines with fluorescently tagged replication factors to track their spatiotemporal dynamics in live-cell microscopy during and after treatment with the replication inhibitors aphidicolin and hydroxyurea. All 3 replicative polymerases were found to accumulate upon stress, with the primase subunits of polymerase alpha retaining catalytic activity. This accumulation was dependent on new origin firing and accompanied by 9-1-1 clamp loading in the vicinity of stalled forks. Pulse-stress-pulse experiments and spatial correlation analyses of replicons indicated that, after stress is removed, replication preferentially restarts at new sites surrounding previously stalled forks. These findings shed light on the replisome’s immediate reaction to stalled replication forks and provide comprehensive kinetic maps of the replisome under stress.

**Highlights:**

- Comprehensible atlas of the replisome under stress
- DNA polymerases accumulate during stress, with clamp switch near stalled forks
- In the absence of DNA synthesis, new adjacent origins fire with active priming
- During stress recovery, replication restart predominantly at new fired origins

## INTRODUCTION

DNA replication is a fundamental biological process ensuring the accurate transmission of genetic information during cell division. It occurs during the S-phase of the cell cycle and relies on the concerted action of the replisome, an assortment of protein complex responsible for unwinding the DNA, synthesizing new strands, and restoring epigenetic information on nascent DNA (Bellelli and Boulton, 2021; Yao and O’Donnell, 2010). The coordination of all these activities is essential, as replication errors or stalling can lead to chromosomal aberrations, genome instability, and ultimately oncogenesis. To prevent this, the initiation and progression of replication are tightly regulated by cell cycle checkpoints that ensure replication occurs only under favorable conditions (Blow and Hodgson, 2002; Diffley, 2011; Técher et al., 2017).

Replisome assembly begins with the licensing of replication origins in the late G1 phase through the loading of the MCM2-7 complex onto chromatin (Blow and Dutta, 2005; Costa and Diffley, 2022; Gonzalez et al., 2005; Stillman et al., 2025). At the onset of S-phase, recruitment of CDC45 and GINS converts this complex into the active CMG helicase, which unwinds the DNA to create replication forks (Moyer et al., 2006; O’Donnell and Li, 2016). The exposed single-stranded DNA (ssDNA) is coated by replication protein A (RPA), preventing degradation and unwanted base modifications (Bellelli and Boulton, 2021). DNA synthesis is then carried out by 3 specialized polymerases (Garcia-Diaz and Bebenek, 2007): polymerase α (PolA)-primase initiates replication by synthesizing short RNA-DNA primers (Jones et al., 2023), while polymerases δ (PolD) and ε (PolE) elongate the nascent strands with the aid of the sliding clamp PCNA, which is loaded by replication factor C (RFC) (De March et al., 2017; Görisch et al., 2008; Johansson and Macneill, 2010). Polymerase ε synthesizes the leading strand continuously, whereas polymerase δ extends primers on the lagging strand into Okazaki fragments that are later processed by the Fen1 endonuclease and joined by DNA ligase 1 (Pursell et al., 2007). Following synthesis, DNA methyltransferase 1 (DNMT1) restores cytosine methylation marks on hemimethylated nascent DNA, thereby preserving epigenetic information (Gaudet et al., 1998; Schermelleh et al., 2007; Spada et al., 2007). These factors are reviewed in (Bellelli and Boulton, 2021; O’Donnell and Li, 2016; Yao and O’Donnell, 2010) and summarized in Figure 1A. Spatially, the replication machinery is not freely diffusing: microscopic evidence proposes that active forks cluster into discrete nuclear focal structures (replication foci) each containing several replicons that are activated and processed together (Berezney et al., 2000; Hozák et al., 1994).

**Figure 1.**
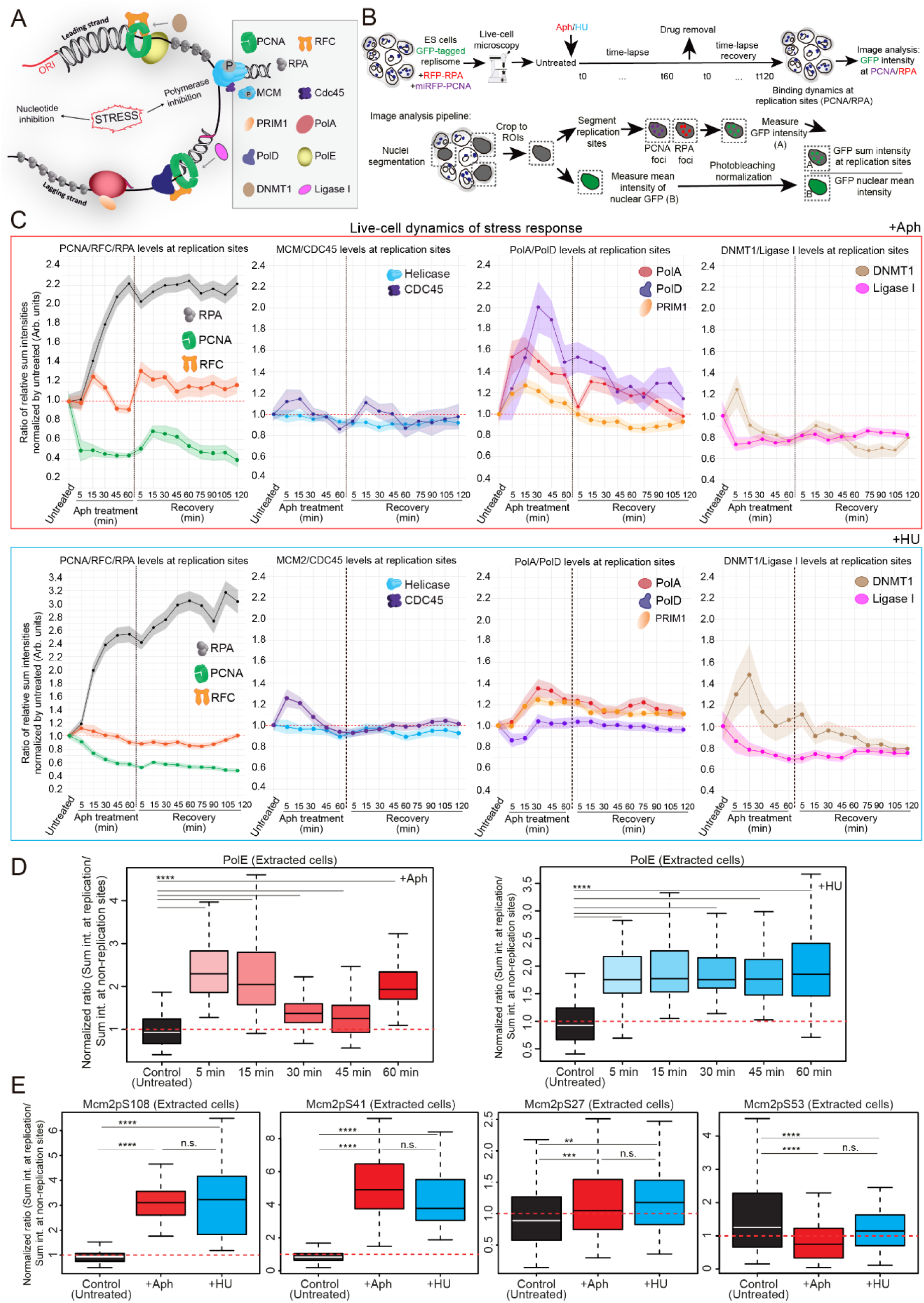
Time-lapse and fixed-cell analysis of replisome factor dynamics upon aphidicolin- and hydroxyurea-induced replication stress. (A) Diagram outlining the location and function of relevant replisome factors at an active replication bubble, along with the inhibitory targets of both aphidicolin and hydroxyurea and a legend assigning simplified shapes to each protein. GFP-tagged cell lines were generated for all factors shown except PolE. (B) Pipeline of the experimental setup and image analysis for tracking replisome factor presence at replication sites over time in live mESCs. Triple-tagged mESCs were imaged by superresolution fluorescence microscopy before, during, and after stress induction with aphidicolin or hydroxyurea. Images were processed using Zeiss ZEN and Volocity. The mean intensity of the GFP-tagged replisome factor of interest was measured within each replication focus and normalized to the mean GFP intensity in the entire nucleus. (C) Live cell quantification of the fluorescence intensity of replisome factors at replication sites before, during, and after replication stress induction with 50 µM aphidicolin (top, red) or 10 mM hydroxyurea (bottom, blue). Values were normalized to the mean nuclear intensity at each time point to account for photobleaching and focus drift. Foci smaller than 0.05 µm² were excluded. Each curve represents 1 replicate and 5 to 10 cells. (D) Fixed-cell immunostaining of polymerase epsilon (PolE) at replication sites during the first hour of replication stress induced by aphidicolin (red) or hydroxyurea (blue) was performed because genome engineering of the PolE locus was not successful even after multiple attempts and targeting strategies. (E) Fixed-cell immunostaining comparing Mcm2 phosphorylation levels at 4 phosphosites (Mcm2pS27, -pS41, -pS53, -pS108) before and after 1 hour of stress induction with aphidicolin (red) or hydroxyurea (blue). In (D) and (E), cells underwent pre-extraction prior to fixation to quantify the chromatin-bound signal of the factor/modification indicated. 2-4 independent replicates were performed with a total of 65-110 cells. Statistical significance was tested with a paired two-sample Wilcoxon test (n.s. is given for p-values ≥ 0.05; 2 stars (**) are given for values < 0.005 and ≥ 0.0005; 3 stars (***) are given for values < 0.0005 and ≥ 0.00005; 4 stars (****) are given for values < 0.00005).

Given the complexity and interdependence of replisome components, it is unsurprising that replication can be disrupted by a variety of stressors. Replication stress, defined as any perturbation that slows or stalls replication progression, can arise from endogenous sources such as nucleotide pool imbalance or DNA damage, as well as exogenous factors like radiation or chemotherapeutic agents (Técher et al., 2017; Zeman and Cimprich, 2014). Persistent or misregulated replication stress is a hallmark of many cancers, but controlled induction of replication stress can also be therapeutically beneficial. In fact, replication inhibitors have been used to target tumor-specific vulnerabilities or to treat disorders such as sickle cell anemia (McGann and Ware, 2015; Ubhi and Brown, 2019). Despite this, little is known about how the DNA replication program responds to replication inhibition, particularly on the level of the individual dynamics of central catalytic replisome components.

In experimental settings, replication stress is often induced using chemical inhibitors that perturb DNA synthesis. 2 of the most commonly used agents are aphidicolin (Aph) and hydroxyurea (HU). Aphidicolin acts as a competitive inhibitor of B-family DNA polymerases, including polymerases α, δ, and ε, thereby directly blocking DNA synthesis while leaving other replisome components unaffected (Baranovskiy et al., 2014; Hanaoka et al., 1979). Notably, the CMG helicase continues to unwind DNA under aphidicolin treatment, resulting in helicase-polymerase uncoupling and the accumulation of RPA-coated ssDNA (Görisch et al., 2008). In contrast, hydroxyurea inhibits ribonucleotide reductase, reducing the cellular pool of deoxynucleoside triphosphates (dNTPs) and indirectly stalling DNA polymerase activity, equally leaving the helicase complex unaffected (Koç et al., 2004; Rosenkranz and Levy, 1965). Despite decades of use, the molecular consequences of these agents on the dynamics and interactions of individual replisome components remain largely uncharacterized. At the level of the replication program, slowing down fork progression is known to elicit the firing of “dormant” origins as a first line of defense in order to complete genome replication (Blow et al., 2011; Woodward et al., 2006). Our recent study in human cancer cell lines revealed that aphidicolin treatment alters the abundance and phosphorylation state of several factors at replication sites, including the helicase subunit MCM2, polymerases α and δ, and PCNA (Pabba et al., 2023). Interestingly, while polymerases α and δ accumulated at replication sites under stress, PCNA dissociated, suggesting a reorganization of the replisome. Since polymerase δ is typically recruited to the replisome via PCNA (De March et al., 2017; Strzalka and Ziemienowicz, 2011), it can be hypothesized that alternative DNA clamps may stabilize polymerase binding under stress conditions. One such candidate is the 9-1-1 complex, a heterotrimeric clamp involved in DNA damage signaling and checkpoint activation (Day et al., 2022; Lee and Park, 2020). However, the mechanisms underlying polymerase retention and replisome remodeling during stress remain elusive.

In metazoa, replication origins are not defined by a strict consensus sequence but are instead specified by a combination of chromatin and epigenetic features. Their use is plastic and developmentally regulated (Ganier et al., 2019; Méchali, 2001; Pradhan et al., 2025, 2024). Mouse embryonic stem cells (mESCs) provide a unique context for studying DNA replication. In contrast to somatic cells, which typically spend most of their cell cycle in G1, mouse embryonic stem cells have an exceptionally short G1 phase of about 1 to 3 hours, while S-phase accounts for roughly 75% of the total cell cycle duration (Rausch et al., 2020; Savatier et al., 2002). This means that, in an asynchronously proliferating mESC culture, the majority of cells are actively replicating their DNA, making them an ideal system to study replication dynamics. Furthermore, mESCs have a stable karyotype and are diploid. In the present work, we used the murine ESC line ESJ1 (Li et al., 1992) as well as human HeLa cervix carcinoma cells as model systems to dissect the dynamic and immediate response of the replisome to stress. Furthermore, we characterized the dynamics of recovery and replication restart after stress removal through three-dimensional mapping of replication sites in fixed and live cells using (superresolution) fluorescence microscopy and AlphaFold modeling. Our data provide new insights into the mechanisms that govern replisome stability and adaptation and a kinetic map of the replisome anatomy and activity under stress.

## RESULTS AND DISCUSSION

### An experimental system to monitor replisome dynamics before intra-S-phase checkpoint activation

To dissect the response of the different replisome components to replicative stress by live-cell microscopy, we used CRISPR-Cas9 genome engineering to generate clonal mouse embryonic stem cell lines with endogenous GFP-tagged replication factors. We selected and characterized cell lines for GFP-tagged RPA, PCNA, RFC2, Mcm2, CDC45, PRIM1, PolA, PolD, DNMT1, and Ligase I (Figure 1A) (Table 1 and Extended data 1 for cell line generation and characterization). To accurately assess the effects of replicative stress in these cell lines, we also introduced fluorescently tagged RPA and PCNA using the Sleeping Beauty transposon system. This experimental approach allowed us to monitor replication and chromatin changes in living cells in real time, using PCNA accumulation as a marker for active replication sites and RPA accumulation at ssDNA as a marker for stalled replication (Görisch et al., 2008; Pabba et al., 2023; Prorok et al., 2024). As a source of replicative stress, we employed treatment with either 50 μM aphidicolin (Aph) or 10 mM hydroxyurea (HU). These concentrations cause DNA synthesis to stop, as determined by the lack of fluorescent nucleotide incorporation in treated samples. To examine the timeline of intra-S checkpoint activation under these experimental conditions, we monitored the levels of Chk1-pSer345 from 0.5 to 24 hours of stress induction. As shown in Figure S1A, between 0.5 and 2 hours, the fraction of Chk1-pSer345 positive cells is below 5% for both stressors, in contrast with the fraction of Chk1 positive cells (Figure S1B). While the Chk1-pSer345 positive fraction starts increasing from 3 hours of treatment, reaching nearly 100% after 12 hours of treatment in both Aph and HU (Figure S1A). We selected 1-hour-long stress treatments to study the response of the replisome to stress in the absence of checkpoint activation, followed by stress recovery after drug removal (2 hours). In these conditions, we performed live-cell time-lapse imaging followed by image analysis (Figure 1B) to quantify the dynamics of the different replisome components and their relative accumulation at replication sites, which were defined as spots of either PCNA or RPA accumulation. This quantification (Figure 1C) already shows notable RPA accumulation and PCNA dissociation 5 to 15 minutes after drug addition. Furthermore, we measured ATR levels after 1 hour of treatment, finding a significant increase for both total and chromatin-bound ATR fractions in both Aph and HU (Figure S1C). This observation has been reported to correspond with a loose checkpoint activation response, characterized by loading of ATR on chromatin but no, or weak, Chk1 activation, as we reported (Figure S1A) (Koundrioukoff et al., 2013). ATR monitors the level of RPA at the forks under low stress conditions, where severely increased RPA accumulation would cause ATR crowding, amplifying checkpoint signals, and shifting from loose to full checkpoint activation (Yin et al., 2021). Overall, these results are in agreement with previously published studies in other cell systems (Görisch et al., 2008; Sporbert et al., 2005) and validate our methodology.

### DNA Ligase I and DNMT1 dissociate from stalled forks following PCNA loss

Since we observed PCNA dissociation from replication sites upon stress induction, we next focused on the dynamics of PCNA-interacting factors that act downstream of DNA synthesis, Ligase I and DNMT1. The levels of the maturation factor DNA Ligase I showed a decrease in response to fork stalling (Figure 1C), as expected in the absence of DNA synthesis and in accordance with our previous data on other cell systems (Görisch et al., 2008; Sporbert et al., 2005). Thus, Ligase 1 behavior under stress can easily be explained by a lack of substrate, which would occur as soon as polymerase delta stops generating Okazaki fragments that require ligation. DNMT1, on the other hand, exhibited a unique dynamic characterized by a brief accumulation within the first 15 minutes of treatment followed by gradual dissociation below the starting level (Figure 1C). The initial spike in DNMT1 accumulation may be explained by the fact that DNMT1 primarily colocalized with large clusters of PCNA before treatment, while accumulation was less apparent at smaller foci. These clusters likely correspond to highly methylated heterochromatic regions, where the increased DNMT1 function is required and independent of PCNA (Qin et al., 2015). As PCNA begins to dissociate upon treatment, the smaller foci disappear first, leaving only the PCNA foci with the highest abundance of DNMT1, leading to a relative increase in its intensity. Beyond 15 minutes of stress, RPA clusters begin to replace dissociated PCNA, once again decreasing the relative DNMT1 intensity within the average replication focus. Similar to Ligase 1, the subsequent dissociation below the initial level may be explained by a lack of substrate as replication stalls. Dissociation of PCNA, previously reported and one of the major observations during stress response, remains to be mechanistically explained.

### Replicative polymerases and primase accumulate during replication stress concomitantly with DNA helicase activation

Upon recovery, we detected de novo loading of PCNA during recovery in live-cell experiments, verified by immunostaining in extracted cells (Figure S2). For the PCNA loader RFC, specifically the subunit RFC2 in live-cells and the subunit RFC1 in fixed cells (Figure 1C and Figure S3), loading levels fluctuated slightly but remained around the threshold of untreated (time 0) cells. Interestingly, we observed that RPA accumulated upon stress and remained highly accumulated during recovery instead of dropping back to the levels observed prior to stress induction (Figure 1C). The accumulation of the DNA helicase (Mcm2 subunit) and its binding partner CDC45 was unaffected both during stress and the recovery period. Considering that the inhibitory effects of Aph and HU occur downstream of the CMG helicase, this was the expected outcome: similar treatments with stress-inducing drugs have previously been shown to lead to an uncoupling of the helicase from the rest of the replisome as it continuously unwinds DNA while polymerase activity is stalled (Görisch et al., 2008), generating large stretches of single-stranded DNA, which explains the observed accumulation of RPA (Figure 1C). RPA accumulation remained high during recovery instead of dropping back to the levels observed prior to stress induction, indicating that the gap between the 2 halves of the replisome is not closed swiftly. Surprisingly, and despite being the direct targets of aphidicolin and hydroxyurea, the catalytic subunits of Primase (PRIM1) and DNA polymerases (PolA1 and PolD1) all showed increased accumulation during stress treatment and decreased during recovery (though they never fell below untreated levels). Although establishment of a cell line with GFP-tagged PolE was not successful, immunostaining of PolE at different time points after stress showed similar accumulation dynamics (Figure 1D). Representative time-lapse movies for all conditions are provided (Supplementary movies 1-18). To confirm that these dynamics reflected the behaviour of the endogenous, untagged proteins, live-cell results were validated in wild-type mESCs by immunostaining against the untagged replisome components (Figure S3). In these experiments, an EdU pulse was performed for 10 minutes before drug addition, and EdU foci were used as a marker for replication forks instead of RPA and PCNA foci (Figure S3A). We quantified the accumulation ratio of each factor in extracted (protein loaded/bound to DNA) cells, finding similar loading dynamics to the live cell experiments (Figure S3).

Because live cell microscopy reports on the bulk abundance of CMG helicase components at replication sites but does not provide information on post-translational modifications such as phosphorylation, known to play an important role in regulating the loading and activity of the helicase complex (Arroyo et al., 2024; Frisbie and Bleichert, 2026; Li et al., 2023), we asked whether the loaded CMG remained active under stress. For this reason, we performed immunostaining against specific phosphorylated residues in the Mcm2 subunit of the DNA helicase (Figure 1E). After 1 hour of stress induction, we found an increase in the levels of chromatin-loaded Mcm2pS108, Mcm2pS41, and Mcm2pS27, while the levels of Mcm2pS53 decreased compared to untreated cells (Figure 1E-right). S108 is a well-characterized ATR/ATM target during replication stress (Cortez et al., 2004; Yoo et al., 2004); S53 and S41 are DDK (Cdc7-Dbf4) target sites required for replication initiation (Montagnoli et al., 2006; Sheu and Stillman, 2010), while S27 is so far less characterized. Increased S108, together with decreased S53, would be consistent with ATR activation and suppression of S53-driven origin firing, previously reported (Zegerman and Diffley, 2010). This is also consistent with the increased level of chromatin-bound ATR we measured (Figure S1C). The latter could occur in favour of S41-mediated origin activation. In this context, continued helicase activity is supported by the increase in loaded Mcm2pS108, a modification linked to stress-response signalling, helicase activation, and chromatin loading. In summary, the overall increase in the levels of phosphorylated helicase demonstrates the maintenance and possible increase of its activity under stress.

The most striking finding was that, despite DNA synthesis being inhibited, all 3 of the replicative polymerases accumulated at replication sites within the first 30 minutes of stress induction. This counterintuitive result raised the question of whether the increase in polymerase presence occurred at preexisting replication forks or whether it can be attributed to the firing of additional replication origins in the proximity of the stalled ones in the absence of checkpoint activation and DNA synthesis. The latter hypothesis is supported by the increase in MCM phosphorylations indicating helicase activity, specifically Mcm2pS108 (stress response signaling and origin firing) and Mcm2pS41 (also responsible for helicase activation during origin firing), and would be consistent with the accumulation of PRIM1, which suggests RNA priming is occurring in preparation for DNA synthesis. Altogether, these data indicated that different replisome factors do not exhibit a single unified stress response, but that their dynamics shift individually. However, the overall trend was the same for both avenues of stress induction across all observed targets.

### In response to replicative stress, PCNA clamp dissociates while the 9-1-1 clamp loads and interacts with the PolA-PRIM1 complex

The retention of DNA polymerases (particularly polymerase delta and epsilon) during replication stress raises another interesting question, as they are commonly considered to rely on PCNA for stable association with DNA: what allows these factors to remain bound at stalled replication forks despite the rapid dissociation of the PCNA clamp? One possible contender for this role is the alternative DNA clamp 9-1-1 (Rad9-Hus1-Rad1), which is commonly associated with states of stress and DNA damage and has been shown to localize to sites of abundant RPA-bound ssDNA that also occur under our treatment conditions. Therefore, we investigated possible interactions between the DNA polymerases and the 9-1-1 complex using AlphaFold 3 modeling. The predictions performed include interactions of different combinations of polymerase subunits and PRIM1 with the 9-1-1 complex and its subunits, and compare them to previously described interactions for reference (e.g., polymerases/PCNA, 9-1-1/Rad17, RFC2-3-4/Rad17, PolE/Rad17) (Figure 2A and Figure S4). Interestingly, we found a comparatively high ipTM confidence score for the full PolA-PRIM1 complex interaction with the full 9-1-1 clamp in the presence of ssDNA, even though higher amino acid count and chain number typically affect ipTM scores negatively. This ipTM score was only marginally lower than the one predicted for PolA-PRIM1’s interaction with the PCNA clamp (0.59 versus 0.6) (Figure 2B). A similar ipTM score (0.58) was calculated for the PolD/9-1-1 complex (Figure 2A). The fact that the whole complex interactions showed much higher scores than partial or one-to-one predictions for subunits explains why the modeling of PolA-Rad9/Hus1/Rad1 complexes did not predict an effective interaction in previous studies (Bryant et al., 2022). We next investigated the structural features of these models, finding that the catalytic subunit of PolA was predicted to interact with Rad1, showing several amino acid contacts between the 2 subunits (involving SER149, ILE264, and GLY262 in Rad1 and GLN555, ASP595, and CYS600 in PolA) (Figure 2B). This is consistent with previous work reporting PolA requirement for loading of 9-1-1 at RPA-covered ssDNA (Yan and Michael, 2009). Interestingly, the PolD catalytic subunit was also predicted to interact with Rad1 at a similar relative position, though different amino acids were predicted to be involved in the interface (LYS57, GLN137, ASP144, and CYS272 in Rad1). Predictions for PolE showed lower ipTMs and predicted no contacts between PolE catalytic subunit and Rad1, though this could be attributed to the high complexity and size of PolE catalytic subunits, which may interfere with interface predictions. Recent in vitro biochemical analysis also suggested that the PolE catalytic subunit interacts with the 9-1-1 complex at the 3’ recessed end (Acharya et al., 2023).

**Figure 2.**
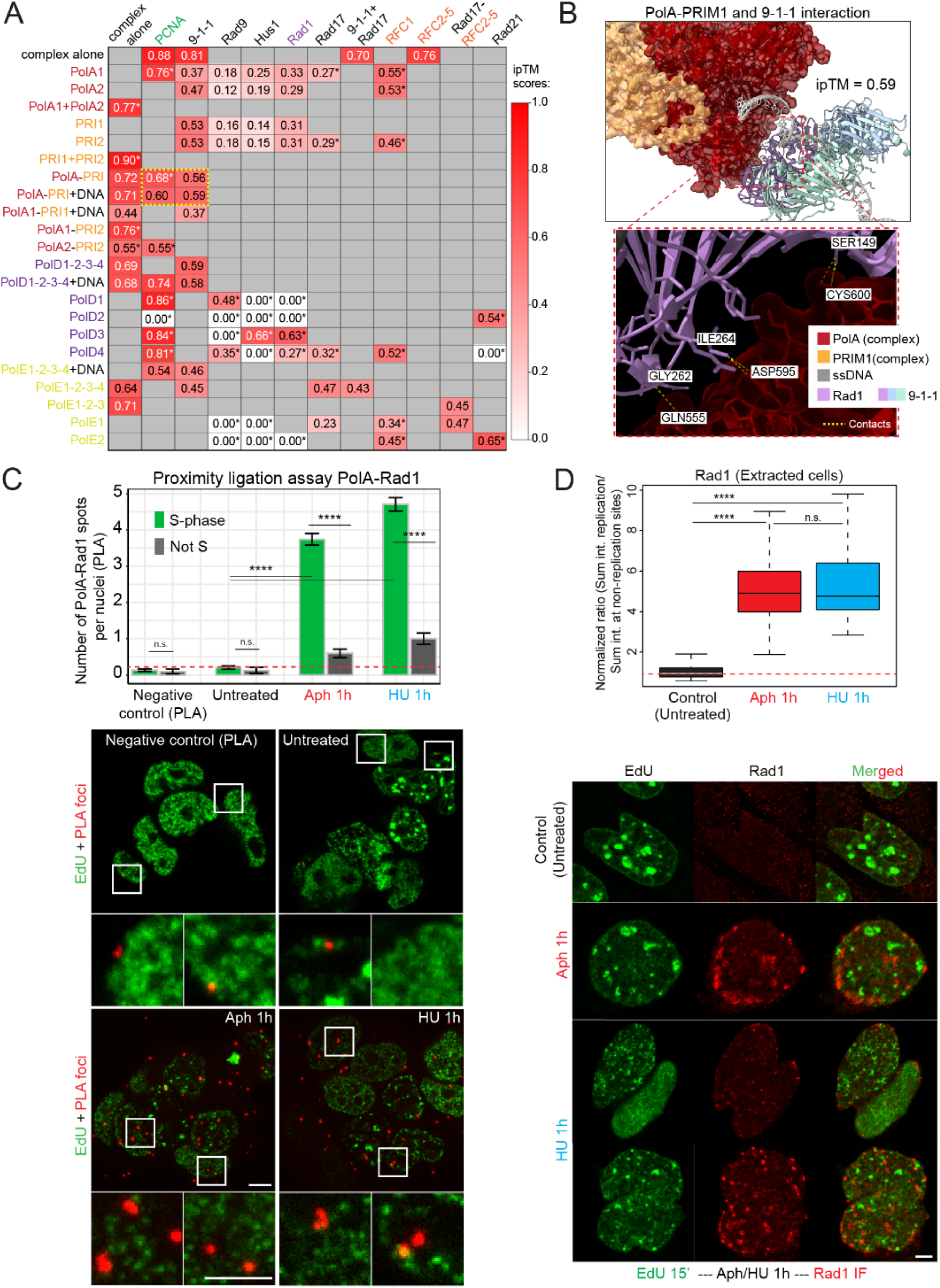
Computational predictions and in situ validation of the interaction between polymerase alpha and the 9-1-1 clamp at replication forks upon stress induction. (A) ipTM confidence scores generated by AlphaFold 3 for the interaction between components of the 9-1-1 clamp, the clamp loader RFC, and the 3 replicative polymerase complexes (including primase). Predicted scores of known interactions (e.g., PolA/PCNA, 9-1-1/Rad17, RFC2-3-4/Rad17) are included for benchmarking. (B) Three-dimensional representation of the predicted complex between the 9-1-1 clamp, PolA, and PRIM1 bound to single-stranded DNA, generated by AlphaFold 3. The predicted interface with PolA is located in the Rad1 subunit of the 9-1-1 clamp; predicted contact residues are indicated (Rad1: Ser149, Ile264, Gly262; PolA: Gln555, Asp595, Cys600). (C) Barplots showing the quantification of a proximity ligation assay (PLA) detecting interactions between the N-terminally GFP-tagged catalytic subunit of polymerase alpha (anti-GFP) and Rad1 (anti-Rad1), with and without 1-hour induction of replication stress (50 µM aphidicolin or 10 mM hydroxyurea). The negative control omitted the addition of the anti-GFP primary antibody. Representative images are shown at the bottom. Each PLA focus (red) corresponds to a site where the 2 targets were less than 40 nm apart. Scale bar: 5 µm. Two replicates were performed, including between 75 and 102 cells. (D) Quantification of an immunostaining for Rad1, showing the Rad1 levels inside replication sites divided by the nuclear Rad1 signal outside of replication sites, normalized to the average of untreated control. Representative images (bottom) show the abundance and distribution of Rad1 (red) throughout the nucleus in untreated, aphidicolin-, and hydroxyurea-treated cells. EdU (green) was pulsed before stress induction to mark pre-stress replication sites. Scale bar: 5 µm. Three replicates were performed, including between 67 and 87 cells. Statistical significance was tested with a paired two-sample Wilcoxon test (n.s. is given for p-values ≥ 0.05; 4 stars (****) are given for values < 0.00005).

Based on these structural predictions, we investigated PolA and Rad1 interaction. We performed proximity ligation assays (PLA) in mESCs with and without replication stress induction (1 h incubation with Aph or HU). Quantification revealed almost no PLA foci for untreated cells (and the negative control without one of the primary antibodies), while stress-induced samples showed a significantly heightened number of nuclear PLA foci in comparison. This increase, indicative of PolA-Rad1 interaction, occurred exclusively in S-phase cells under replicative stress (Figure 2C), validating the AlphaFold predictions under these conditions. In addition, we performed immunostaining and quantification of Rad1 levels (chromatin-bound levels, as performed in Figures S3). Compared with untreated cells, replication stress induced with Aph or HU was associated with a significant increase in the accumulation of Rad1 (Figure 2D). This accumulation was observed near EdU foci that mark stalled forks (EdU pulse before drug addition). As an additional line of evidence, we performed live cell time-lapse on HeLa cells expressing GFP-Rad9, showing that another one of the 9-1-1 subunits also accumulates in response to stress, concomitantly with PCNA dissociation and increased polymerase loading (Figure S5). Based on these data, we propose a “clamp switch” mechanism in which dissociation of PCNA from replication sites gives way to 9-1-1 loading by interaction with PolA. However, this loading does not seem to occur exactly at stalled forks (marked by EdU before stress), but in nearby and partially overlapping positions (Figure 2D bottom). This suggests a repositioning of replisome components under stress and points to their recruitment to the previously hypothesized newly activated origins rather than mere accumulation at pre-existing stalled forks.

### RNA primer synthesis continues as polymerases accumulate upon stress

Based on the data, 2 possible scenarios could be envisioned: in scenario 1, more polymerases accumulate at existing stalled forks, whereas in scenario 2, new (dormant) origins increasingly fire during stress induction in the presence of stalled forks and the absence of DNA synthesis. To test and distinguish between these hypotheses, we first measured the levels of DNA/RNA hybrids, since priming of ssDNA could occur at newly activated sites in the presence of Aph/HU. Immunostaining against DNA/RNA hybrids in extracted cells (removal of DNA/RNA hybrids due to transcription) surprisingly showed a significant increase in their levels for both stress treatments compared to the untreated sample (Figure 3A and Figure S6A). To verify these results and exclude background noise, we performed PLA analyses between DNA/RNA hybrids and either PRIM1 or PolA. In both cases, we again observed a significantly higher number of nuclear PLA foci in S-phase cells under Aph or HU treatment (Figure 3B-C and Figure S6B). A feasible explanation for this observation relies on the fact that, during unaltered replication, primers are transitory structures rapidly removed (Harrington and Perrino, 1995; Yuan et al., 2023). However, during stress, priming would occur either after the firing of dormant origins or at the existing lagging-strand stalled forks, but the replication machinery would not progress further to DNA synthesis. The PolA-PRIM1 complex would remain at these structures for a longer period of time, explaining the increased number of PLA foci and the increased accumulation observed under stress (Figure 1C, Figure S3B). AlphaFold structural analysis of PolA-PRIM1-DNA/RNA interaction is shown in Figure 3D, with an ipTM of 0.55 for PolA1 and PRIM1 subunits forming an interface with primed ssDNA.

**Figure 3.**
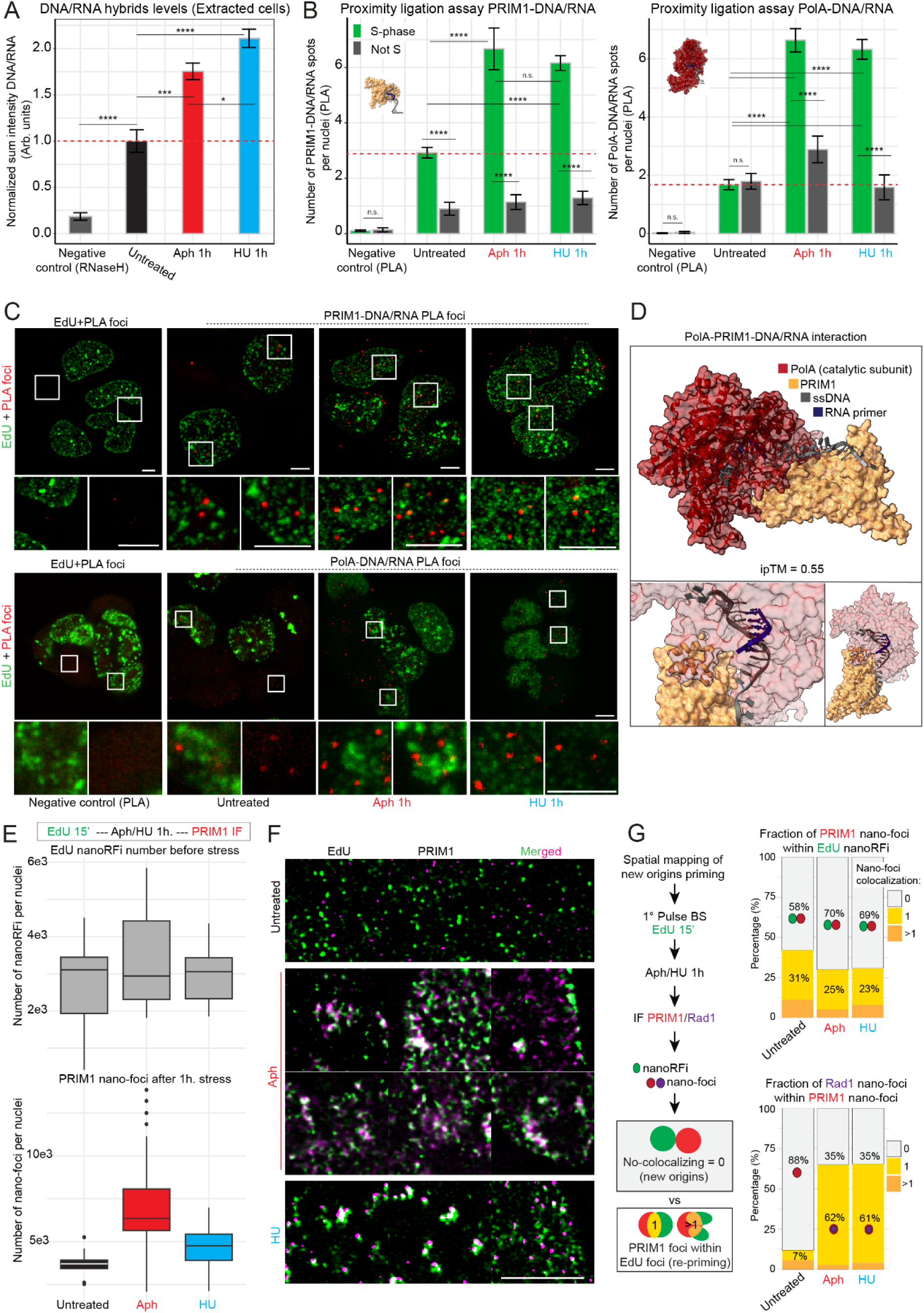
Assessment of primase activity and DNA/RNA hybrid (primer) formation upon stress induction. (A) Quantification of total nuclear DNA/RNA hybrid levels (S9.6 antibody) measured by immunostaining in pre-extracted mESCs before and after 1 hour of replication stress with aphidicolin or hydroxyurea. 2 independent replicates were performed, representing a total of 30-40 cells per condition. (B) Quantification of the average number of nuclear PLA foci per cell corresponding to interactions between DNA/RNA hybrids and PRIM1 (left) or PolA (right) in untreated, aphidicolin-, and hydroxyurea-treated cells. Negative controls were performed using only the primary antibody against DNA/RNA hybrids (S9.6). 2 independent replicates were performed, representing a total of 70-90 cells per condition. For experiments in (A) and (B), statistical significance was tested with a paired two-samples Wilcoxon test (n.s. is given for p-values ≥ 0.05; one star (*) for p-values < 0.05 and ≥ 0.005; two stars (**) are given for values < 0.005 and ≥ 0.0005; three stars (***) are given for values < 0.0005 and ≥ 0.00005; four stars (****) are given for values < 0.00005). (C) Representative images of the proximity ligation assays examining DNA/RNA hybrid colocalization with PRIM1 (top) and PolA (bottom). PLA foci are shown in red; EdU signal (pulse before stress) is shown in green. Scale bar: 5 µm. (D) AlphaFold 3 structural prediction of the PolA-PRIM1 complex bound to primed single-stranded DNA. The ipTM score for this prediction is 0.55. (E) Top: number of EdU nano-foci per nucleus before stress in pre-extracted mESCs. Bottom: number of PRIM1 nano-foci per nucleus after 1 hour of treatment with aphidicolin or hydroxyurea, compared with untreated controls. Foci were quantified by superresolution microscopy followed by 3D segmentation. (F) Representative superresolution images showing the distribution of EdU (green, pulse before stress) and PRIM1 (magenta) signals in pre-extracted mESC nuclei before and after stress induction. Scale bar: 5 µm. (G) Left: schematic representation of the analysis pipeline. Top right: quantification of overlap between PRIM1 nano-foci and EdU-labelled nanoRFi (stalled forks). Bottom right: quantification of overlap between Rad1 nano-foci and PRIM1 nano-foci. For (E) and (G), three independent replicates were performed, corresponding to 77-103 total cells per condition.

To further investigate the priming events during stress, we performed PRIM1 immunostaining followed by superresolution microscopy and single-cell quantification of nano-foci. The experiments were performed in extracted cells, which removes non-bound PolA-PRIM1 complexes, allowing us to exclusively detect DNA-bound primase. An EdU pulse before stress induction was used to mark replication sites and showed a similar number of nano-replication foci (nanoRFi, which correspond to replicons (Rausch et al., 2020) (Chagin et al., 2016) for all samples (Figure 3E top). However, quantification of PRIM1 nano-foci after 1 hour of treatment with Aph or HU revealed significant differences between untreated and stressed cells. We found an approximate 2-fold increase in the number of PRIM1 nano-foci after Aph treatment and a 1.5-fold increase after HU treatment (Figure 3E bottom). Representative superresolution images illustrate this increase and showcase the relative spatial pattern of both EdU and PRIM1 signals (Figure 3F). We next calculated the colocalization between both signals to discriminate whether the PRIM accumulation took place at the forks stalled upon stress or at newly fired (previously dormant) origins. Most of the PRIM1 nano-foci (around 70%) did not overlap or only partially overlapped with the nanoRFi corresponding to stalled forks (Figure 3G top). This outcome indicates that the majority of priming takes place at newly fired origins and only a small percentage is re-priming at the existing stalled forks. This also explains our observation in PLA experiments showing increased interaction of DNA/RNA hybrids with both PolA and PRIM1 upon stress, and it is in agreement with increasing phosphorylation of the DNA helicase (Figure 1E).

Furthermore, we analyzed the colocalization between PRIM1 and Rad1 nano-foci, showing that more than 60% of Rad1 foci are located within PRIM1 foci, in contrast with untreated samples with significantly lower Rad1 signal (Figure 3G bottom). This supports a model in which new origins fire near the stress-stalled forks and recruit polymerases as well as the 9-1-1 clamp. In agreement with the domino model of origin firing, and with the approximately two-fold increase in PRIM1 nano-foci observed for Aph, 2 adjacent origins could be firing forward and backwards of the stalled one. These results raise the next question: whether newly fired origins are within the same replication domains or not.

### Increased accumulation of the active DNA helicase and polymerases upon stress is abolished by CDK inhibition

To validate our previous results pointing to firing of dormant origins and differentiate between the 2 possible scenarios described, we inhibited *de novo* origin activation by treatment with the cyclin-dependent kinase (CDK) inhibitor roscovitine, which prevents phosphorylation of Mcm proteins by CDKs (particularly CDK2). We performed the same stress treatments (1 hour Aph or HU) with the simultaneous addition of the CDK inhibitor, followed by immunostaining and quantification by image analysis and flow cytometry (Figure 4A). In immunofluorescence images, we observed that the increase in Mcm2pS108 (ATR/ATM target during replication stress) and PolA levels upon stress induction did not occur when stress was induced in the presence of the CDK inhibitor (Figure 4B). Both image analysis and flow cytometry quantification confirmed this observation for Mcm2pS108 (Figure 4C). Therefore, de novo phosphorylation of Mcm2 in response to stress does not occur in the absence of CDK activity. We found a similar lack of accumulation upon combined stress and CDK inhibitor treatment for PolA, PolD, and PolE (Figure 4D).

**Figure 4.**
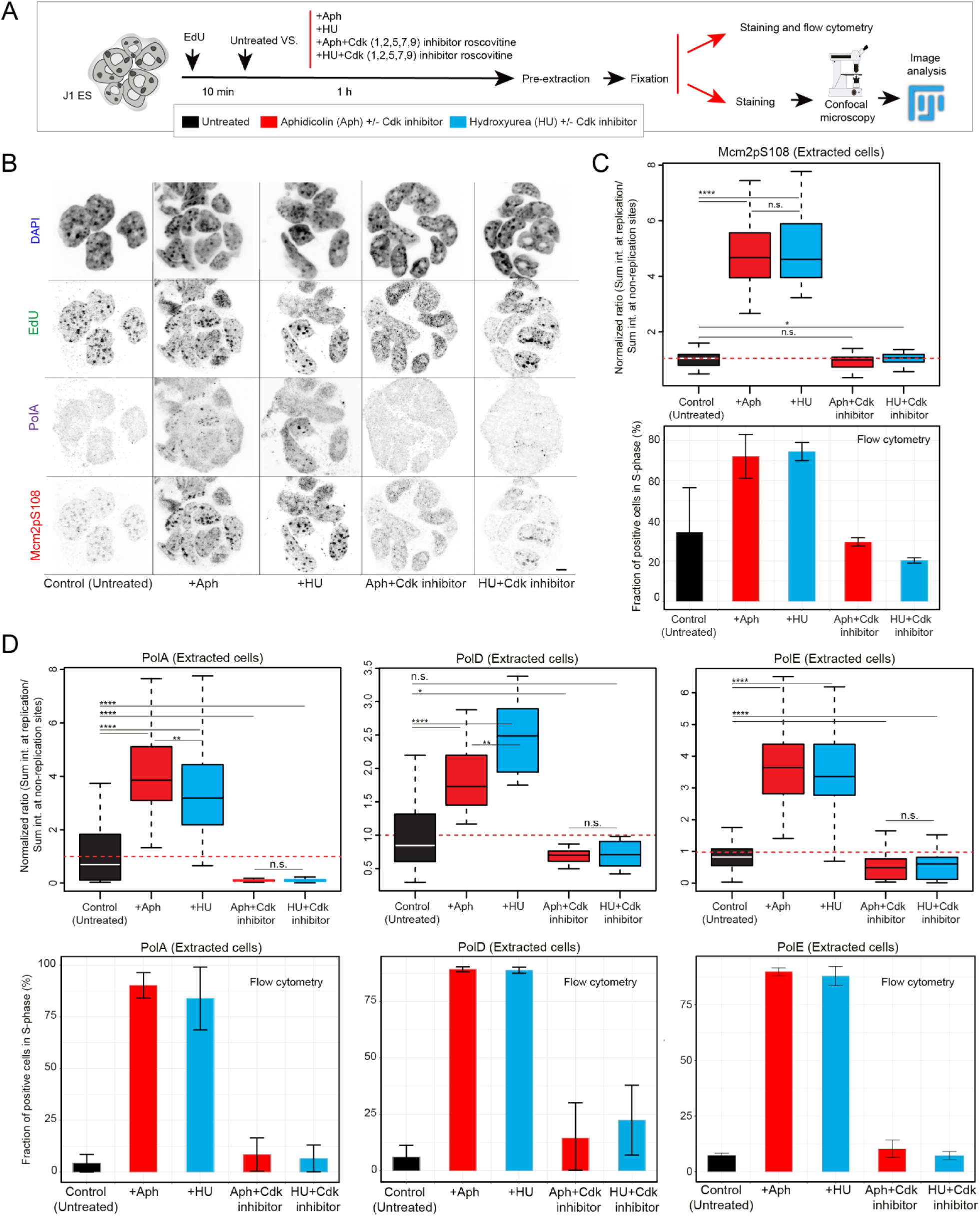
Effect of simultaneous CDK inhibition on replicative polymerase accumulation and MCM phosphorylation at replication sites under stress. (A) Treatment and analysis pipeline. To label replication sites, cells were treated with EdU for 10 minutes prior to the addition of a replication inhibitor (50 µM aphidicolin or 10 mM hydroxyurea), CDK inhibitor (5 µM roscovitine), both, or neither, for 1 hour. Unbound protein was removed by pre-extraction, and cells were fixed, stained, and analyzed through fluorescence microscopy or processed for flow cytometry. (B) Representative images of chromatin-bound Mcm2pS108 and PolA at replication sites under all conditions described in (A). Scale bar: 5 µm. (C) Top: box plots representing chromatin-bound Mcm2pS108 detected at replication sites by immunofluorescence. Bottom: bar plots showing the fraction of Mcm2pS108-positive cells detected by flow cytometry. (D) The same experimental approach and quantification as in (C) were performed for chromatin-bound PolA, PolD, and PolE. For flow cytometry experiments, following pre-extraction, fixation, and immunostaining of the protein of interest, cells were counterstained with propidium iodide and sorted. Barplots show the data corresponding to mean ± standard deviation from 3 independent replicates, including approximately 5000 cells total. Box plots represent 2-4 replicates, including between 80-95 total cells per condition. Statistical significance was tested with a paired two-samples Wilcoxon test (n.s. is given for p-values ≥ 0.05; one star (*) for p-values < 0.05 and ≥ 0.005; two stars (**) are given for values < 0.005 and ≥ 0.0005; three stars (***) are given for values < 0.0005 and ≥ 0.00005; four stars (****) are given for values < 0.00005).

Altogether, our data show that replication stress induces a significant increase in the levels of the active DNA helicase (phosphorylated form of the MCM complex: Mcm2pS108, Mcm2pS41, and Mcm2pS27), and the loading of all 3 replicative polymerases. This does not occur when cells are simultaneously subjected to CDK inhibition, confirming that the stress-induced recruitment of replication factors (active DNA helicase, PRIM1, and polymerases) relies on *de novo* activation of dormant origins. The fact that CDK inhibition led to polymerase levels that were similar or even below those of the untreated cells could be attributed to reduced loading from stalled forks. In these conditions, similar Mcm2pS108 levels to those of untreated cells agree with previous reports showing that the DNA helicase continues DNA unwinding activity in the presence of replicative stress (Devbhandari and Remus, 2020; Kose et al., 2019; Zafar and Nakagaw, 2011). CDK inhibition, simultaneously with stress induction, is not expected to affect this accumulation since it depends on previously phosphorylated MCMs. Following this reasoning, if the observed accumulation of polymerases occurs at pre-existing stalled forks, CDK inhibition would not be expected to affect it. However, because Mcm2pS108/PolA/PolD/PolE accumulation relies on the de novo activation of dormant origins, inhibiting CDK alongside stress induction prevents this activation and their accumulation.

### Replication stress increases the number of active replicons during recovery

As our results indicated firing and priming of new origins in the absence of checkpoint activation, we performed a detailed mapping of dormant origin activation in response to stress. To extend our previous results and map the location of new firing origins relative to stalled forks, we labelled nascent DNA synthesis before stress and either immediately after stress removal (AS) or after 1 hour recovery (Figure 5A). We selected 1 hour as the estimated time in which most replicons would have completed replication and new ones would have fired, based on mESC replication parameters (replicon size: median of 77 kbp; and replication fork speed: median of 1.59 nucleotides per minute) (Rausch et al., 2020). At these time points, we performed correlative microscopy analysis of confocal versus superresolution images. This method allows us to map replicons (nanoRFi) in relation to replication foci/domains (RFi) in single cells, and to measure parameters like replication synchrony (number of active replicons within a replication domain at the same time) and recovery dynamics. Following this approach (Pradhan et al., 2024), we compared the number and the location of replication foci before stress (BrdU-labelled), with replication foci after stress removal and after 1 hour of recovery (EdU-labelled). Compared with pre-stress measurements (BrdU foci), we found that the average number of RFi and nanoRFi after stress removal (EdU foci) was increased, and this increase was still present after 1 hour of recovery (Figure 5B and Figure S7A). Accordingly, the ratios of EdU/BrdU nanoRFi per nucleus were increased in both recovery time points, indicating a higher number of active replicons during stress recovery (Figure 5C). The number of RFi was lower for HU than Aph right after stress removal, though this was no longer the case after 1 hour of recovery. This was to be expected, as HU-induced nucleotide depletion requires more time to be alleviated than the simple competitive binding inhibition provided by Aph.

**Figure 5.**
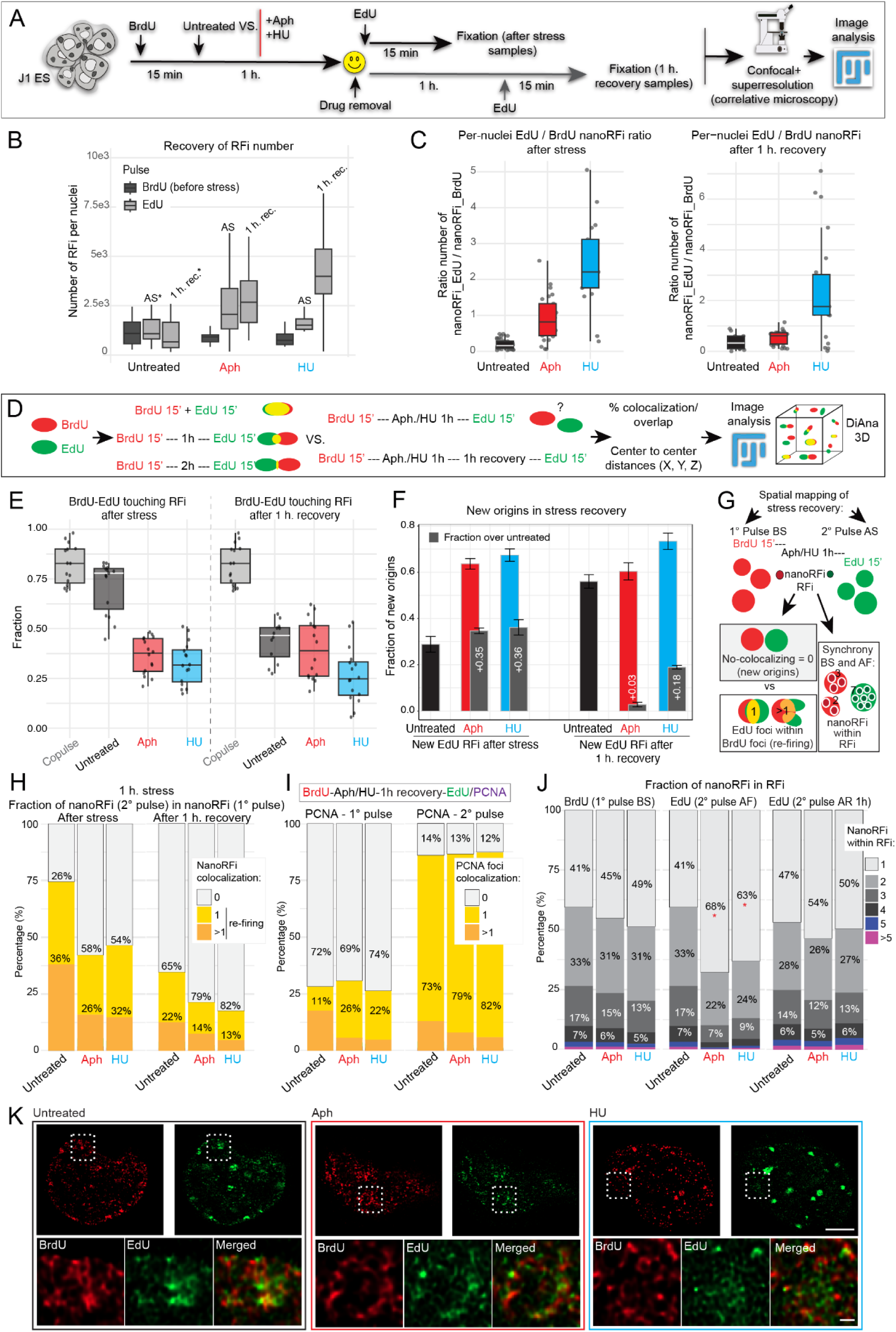
Three-dimensional mapping of replication patterns in S-phase cells before and after replication stress induction. (A) Treatment and imaging pipeline for pulse-stress-pulse experiments. Cells were pulsed with BrdU for 15 minutes, then either left in culture medium or treated with 50 µM aphidicolin or 10 mM hydroxyurea for 1 hour. The drug was washed off, and cells were pulsed with EdU for 15 minutes either immediately or after 1 hour of recovery. They were subsequently fixed, stained, imaged, and analyzed in Fiji. Nano-foci corresponding to individual resolved replication sites (nanoRFi) were determined by superresolution microscopy. Clusters of multiple replication sites at lower spatial resolution were defined as replication foci (RFi) using confocal/widefield imaging. (B) Number of RFi per nucleus before stress (BrdU, dark gray) and after stress (EdU, light gray). (C) Per-nucleus ratio of EdU/BrdU nanoRFi. Left: EdU pulse immediately after stress removal; right: EdU pulse 1 hour after stress removal. (D) Analysis scheme illustrating how the overlap of EdU and BrdU foci was quantified using three-dimensional distance analysis with the DiAna plugin (Fiji). For each pair of segmented BrdU–EdU objects within the same nucleus, center-to-center distance, individual object volume, absolute overlap volume, and fraction of overlapping volume were extracted. Pairs sharing at least 1 voxel were classified as touching. (E) Fraction of post-stress EdU foci touching pre-stress BrdU foci. In the co-pulse control, both BrdU and EdU were administered simultaneously and used as a positive control for the maximum overlap given the spatial resolution of confocal microscopy. (F) Fraction of post-stress EdU foci not touching any pre-stress BrdU focus. Left: EdU pulse immediately after stress removal; right: EdU pulse 1 hour after stress removal. (G) Analysis scheme for correlative microscopy of superresolution and confocal images, used to quantify nanoRFi within RFi (replication synchrony) and to map the colocalization of pre- and post-stress nanoRFi. BrdU nanoRFi were segmented in three dimensions and used as masks to quantify EdU nanoRFi within the same nuclei. (H) Distribution of the number of post-stress EdU nanoRFi found per pre-stress BrdU nanoRFi. Left: EdU pulse immediately after stress removal; right: EdU pulse 1 hour after stress removal. (I) Distribution of the number of PCNA foci colocalizing with BrdU-labelled stalled forks (left) versus EdU-labelled post-stress forks (right), quantified 1 hour after stress removal. (J) Distribution of the number of nanoRFi contained within individual RFi (replication synchrony) before stress, immediately after stress removal, and 1 hour after stress removal, for untreated, aphidicolin-, and hydroxyurea-treated cells. (K) Representative superresolution images of pre-stress (BrdU, red) and post-stress (EdU, green) replicating DNA in untreated, aphidicolin-, and hydroxyurea-treated mESC nuclei. Scale bar: 5 µm. For all experiments, 3 independent replicates were performed, including 79-95 cells total per condition.

Next, applying correlative image analysis, we investigated the position of the new replicons in relation to replication domains observed before stress. We found that most of the new replication foci did not show overlap with pre-stress replication sites, 80% in Aph and 67% in HU (Figure S7B). To exclude the possibility that the differences in EdU and BrdU detection strategies affected these data, we also performed the same experiments with both nucleosides switched (Figure S7C). Using EdU to label stalled forks and BrdU to label new firing origins during stress recovery yielded similar numbers of replication foci (Figure S7D) and 2nd pulse / 1st pulse ratios (Figure S7E). Representative images of BrdU/EdU nanoRFi and an example of the segmentation approach are shown in Figure S8. These findings are in line with our previous observation that stress leads to increased PRIM1-DNA interaction as well as priming, as this would be expected to result in the described increase in replicon number during recovery.

### Mapping of dormant origins firing and stress recovery response shows replication restart at new domains adjacent to stalled forks

We further analyzed the colocalization between stalled forks (BrdU-labelled) and after-stress origins (EdU-labelled) using a different image analysis approach, 3D colocalization using the FIJI plugin DiAna 3D (Figure 5D) (Gilles et al., 2017). This allowed us to map the relative position and distance between BrdU and EdU foci, obtain detailed information about their volumes, and the fraction of overlap. For touching BrdU-EdU foci, the average fraction of volumes that show overlap in Aph/HU-treated samples was below 10%, a percentage even lower than that of untreated cells, in which replication progressed normally during 1 hour. Therefore, the spatial separation of pre- and post-stress replicons is higher than the separation achieved during 1 hour of unaffected replication progression. Co-pulse samples used as reference showed the highest percentage of colocalization (Figure S7F). Also, the distances center-to-center between BrdU and EdU 3D foci were much higher for stress samples, indicating more separation between stalled forks and post-stress replicons (Figure S7G). But importantly, these values were calculated for “touching” BrdU-EdU foci showing some degree of overlap. In agreement with the correlative analysis of Figure S7B, this analysis also showed that less than 40% of post-stress replication foci are touching stalled forks (Figure 5E). Over 60% of foci represent new origins after stress compared to just 30% in untreated cells, meaning that new origin firing under stress conditions greatly exceeds the expected amount of origin firing during unperturbed replication (Figure 5F and Figure S7H). Similar results were obtained by analyzing the fraction of EdU nanoRFi overlapping with BrdU RFi (Figure S7I-J). It can be posited that the aforementioned increase in primase activity upon stress occurs at these additional newly fired origins.

To quantify the fraction of replicons corresponding with newly fired origins after stress (AS) versus re-firing of stalled forks labelled before stress (BS), we performed correlative microscopy image analysis in superresolution 3D stacks. For this analysis, BrdU nanoRFi were three-dimensionally segmented and used for masking and quantification of EdU nanoRFi, returning values of 0 if no EdU nanoRFi falls within the BrdU mask. The same approach was used to quantify replication synchrony before and after stress, in this case comparing the number of nanoRFi (superresolution) that fall within the same RFi domain (confocal) for each separate pulse (Figure 5G). Spatial mapping of stress recovery shows that 58% to 54% of nanoRFi after stress removal do not colocalize at all with pre-stress nanoRFi, which must therefore correspond with newly fired origins. In contrast, such newly fired origins only corresponded to 26% of post-stress nanoRFi in untreated cells. These percentages are even higher after 1 hour of recovery, highlighting how replication progression affects the spatial resolution of labelled replicons and showing that replication predominantly continues at new origins after stress (Figure 5H). In line with this, correlative analysis showed that 69-74% of PCNA foci do not colocalize with BrdU-labeled stalled forks, but instead accumulate at EdU-labeled newly replicating origins after 1 hour of recovery (79 to 82%) (Figure 5I). When we analyzed replication synchrony (number of nanoRFi per RFi, or number of origins firing simultaneously), we found that replication synchrony is decreased immediately after stress removal, showing higher percentages of domains with 1 single replicon after Aph or HU treatments. However, synchrony returns to almost baseline levels after 1 hour of recovery (Figure 5J). These analyses showcase the dynamics of stress response and replication restart after stress removal: the newly fired origins commonly occur as single replicons (nanoRFi) instead of clusters, localized in replication domains (RFi) that are separate from the ones replicating before stress. This matches the higher number of RFi and nanoRFi (around double that before stress) during recovery. In agreement with the domino effect theory (Löb et al., 2016; Maya-Mendoza et al., 2010; Sporbert et al., 2002), after 1 hour of recovery, replication synchrony increased due to these single origins facilitating the firing of neighboring ones.

To assess whether the locations of newly fired origins (EdU) are randomly distributed or dependent on the locations of previously stalled forks (BrdU), we performed the Kolmogorov-Smirnov (KS) test for complete spatial randomness (Figure S9). This non-parametric test quantifies the maximum difference between the empirical cumulative distribution function (CDF) of the data and the theoretical cumulative distribution function for complete spatial randomness (CSR). Spatial randomness is calculated using nearest neighbor distance (r), the shortest distance from each point (nanoRFi) to its closest neighbor. Applying the KS test to the distribution of nanoRFi 3D coordinates for each cell, we found that the empirical line for BrdU nanoRFi before stress was above the theoretical line for complete spatial randomness, meaning points were clustered, and neighboring foci were on average closer than expected for random distribution. Untreated samples and +Aph/HU showed almost identical distributions (Figure S9A-left). Similar trends were observed for EdU nanoRFi 1 hour after stress removal (Figure S9A-right), with curves above the theoretical distribution and close to untreated samples. In addition, neither stress nor recovery time had a significant impact on differences in the distance between empirical and theoretical curves (KS-D, maximum vertical difference between CDF and CSR). This shows that replication stress did not increase the degree of randomness of nanoRFi distribution after stress removal (Figure S9B). Instead, the similar position of the empirical curves observed pointed to similar patterns of nanoRFi positions before and after stress. These data agree with our previous visual judgment: while they did not appear to overlap, the EdU and BrdU patterns of nanoRFi appeared highly correlated, with EdU foci appearing in the vicinity of BrdU foci. Representative images of pre- and post-stress replicating DNA compared with untreated cells are shown in Figure 5K, illustrating both the low level of overlap and the proximity of BrdU and EdU labelled DNA before and after stress.

### Stress recovery response showing replication restart at new proximal domains is conserved between mouse and human cells

In the previous sections, we have shown the replisome response to stress prior to checkpoint activation, followed by its recovery upon stress removal, in mouse ESCs. In addition, we investigated these scenarios in HeLa cells to elucidate whether the stress response is similar in human cancer cells. Following the same experimental pipeline, stress treatments, and image analysis (Figure 6A), we found a similar increase in the number of RFi after 1-hour stress treatment and after 1-hour recovery. This increase was less pronounced for HU after stress removal but reached higher levels after 1 hour of recovery, indicating a slower recovery dynamic for HU treatments (Figure 6B). In line with these results, per-nuclei ratiometric analysis of EdU and BrdU nanoRFi numbers showed an increased EdU/BrdU ratio compared with untreated cells, also higher for HU after 1 hour recovery (Figure 6C). These results reflect a similar stress recovery dynamic, with an increased number of nanoRFi corresponding to active replicons firing after stress removal. Similarly to mESCs, representative images illustrate how the new nanoRFi (EdU) do not colocalize with stalled forks (labelled with BrdU before stress), but are in their vicinity (Figure 6D). In contrast with untreated cells, in which replication progresses unchallenged for 1 hour, stress-treated samples seemed to show lower overlap. In our previous study (Pabba et al., 2023), we found comparable dynamics in HeLa cells for the replisome factors RPA, PCNA, Mcm2pS108, PolA, and PolD (Pabba et al., 2023). Furthermore, the data on PCNA and 9-1-1 using HeLa cells (Figure S5) also support the proposal of a clamp switch. All this evidence supports similar stress response dynamics in murine and human cells.

**Figure 6.**
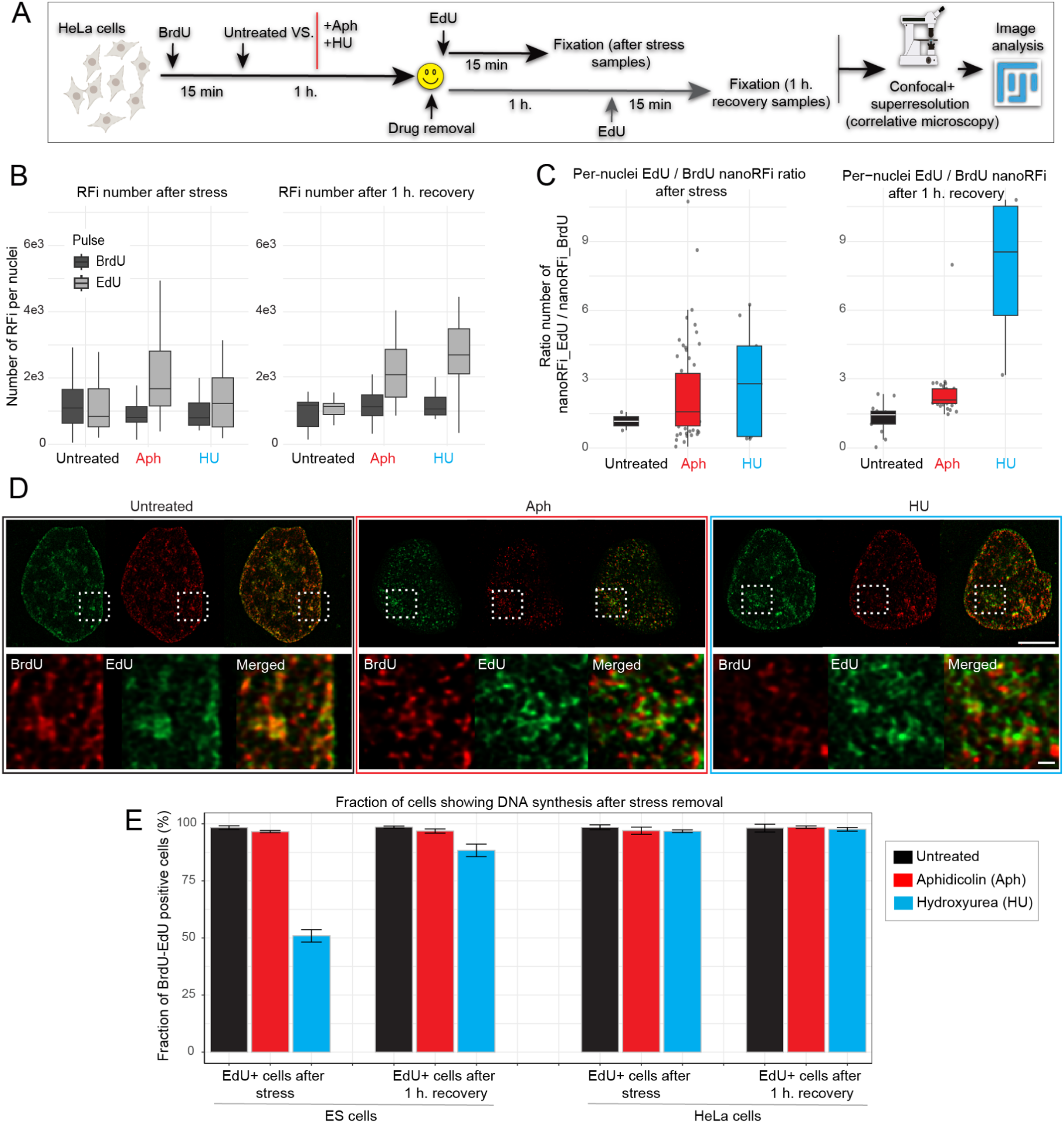
Stress recovery dynamics in HeLa Kyoto cells. A) Treatment and imaging pipeline for pulse-stress-pulse experiments in human cells. Cells were pulsed with BrdU for 15 minutes, then either left untreated or treated with 50 µM aphidicolin or 10 mM hydroxyurea for 1 hour. After drug removal, cells were pulsed with EdU for 15 minutes either immediately or following 1 hour of recovery. They were then fixed, stained, imaged by confocal and superresolution microscopy, and analyzed in Fiji. (B) Number of replication foci (RFi) per nucleus before stress (BrdU, dark gray) and after stress (EdU, light gray) in HeLa cells. Left: EdU pulse performed immediately after stress removal; right: EdU pulse performed 1 hour after stress removal. Each box plot represents at least 20 cells from 2 independent replicates. (C) Per-nucleus ratio of EdU/BrdU replication nano-foci (nanoRFi) in HeLa cells. Left: EdU pulse immediately after stress removal; right: EdU pulse 1 hour after stress removal. (D) Representative superresolution images of HeLa nuclei showing the spatial distribution of BrdU-labelled (red) and EdU-labelled (green) replication foci in untreated, aphidicolin-, and hydroxyurea-treated cells. Scale bar: 5 µm. (E) Percentage of S-phase cells negative for the second nucleotide pulse (EdU) in mESCs (left) and HeLa cells (right), quantified immediately after stress removal and after 1 hour of recovery. Two independent replicates were performed, totaling 60-71 cells per condition.

The slower recovery dynamics after HU treatment compared to Aph we found earlier likely result from the more indirect mechanism of inhibition requiring upstream processes to occur before replication can resume. However, this response to HU differed between mESCs and HeLa cells. In mESCs, we observed a higher fraction of cells that were not labelled (or had almost undetectable levels) by the second pulse of modified nucleotides (EdU negative) after the removal of HU. We quantified these phenomena in all conditions, finding that around 50% of S-phase mESCs were negative for the second pulse after stress removal, corresponding with cells that did not restart DNA synthesis immediately. However, after 1 hour of recovery, more than 90% of S-phase mESCs were positive for the second pulse. For HeLa cells, recovery occurred immediately after both Aph or HU stress removal, showing a faster restart of the nucleotide pathway and DNA synthesis (Figure 6E). From these experiments, we can extract that apart from changes in the speed of DNA synthesis restart, likely due to differences between the ESC and cancer cell metabolism, stress recovery response occurs similarly in mouse and human cells, with a higher number of replicons active after stress removal that predominantly do not correspond to the previously stalled forks.

### Recovery after checkpoint activation follows a similar replication restart dynamic to recovery pre-checkpoint

Most of the studies focused on DNA replication stress using Aph or HU have historically been performed under conditions that induce S-phase checkpoint activation. However, the earlier response of the replisome components to these challenges and the dynamics of DNA synthesis restart in the absence of checkpoint signaling were not characterized. In this study, we focused on these conditions and exhaustively mapped the dynamic response of the different replisome components as early as possible upon stress induction. We monitored this response during 1 hour of DNA synthesis hijacking, and described the dynamics and characteristics of recovery at the nanoscale and single-cell level. Given the unique approaches of our study compared with other studies in the field, it was important to contrast with conditions that induce checkpoint activation. As shown in Figure S1, in our cellular models, Chk1 phosphorylation as a marker of checkpoint activation was seen in around 50% of mESCs after 3 hours and in nearly 100% of cells after 12 hours. Because of this, we also performed the previous analysis of foci count and localization under these conditions (Figure 7A). In general, we observed similar recovery results to 1-hour treatments, with some interesting differences between 3 hours and 12 hours. For 3-hour stress treatments, we found that the number of RFi was increased after stress removal compared to untreated cells, along with the previously observed increase in nanoRFi ratios (EdU/BrdU), and the slower recovery dynamic for HU (Figure 7B). To exclude the possibility of a partial checkpoint activation (only for a fraction of cells, around 50% for Aph and 25% for HU, Figure S1) affecting our results, we looked at 12-hour treatments. In this case, we still observed a similar response: an increased number of RFi after stress removal, and increased EdU/BrdU ratios. However, the dynamics of response seemed slower, with a much smaller increase in these parameters after stress removal, which then increased after 1 hour of recovery. The latter was more obvious in the EdU/BrdU ratios when comparing after-stress versus 1-hour recovery samples. Still, HU-treated samples showed the slowest recovery dynamic (Figure 7C).

**Figure 7.**
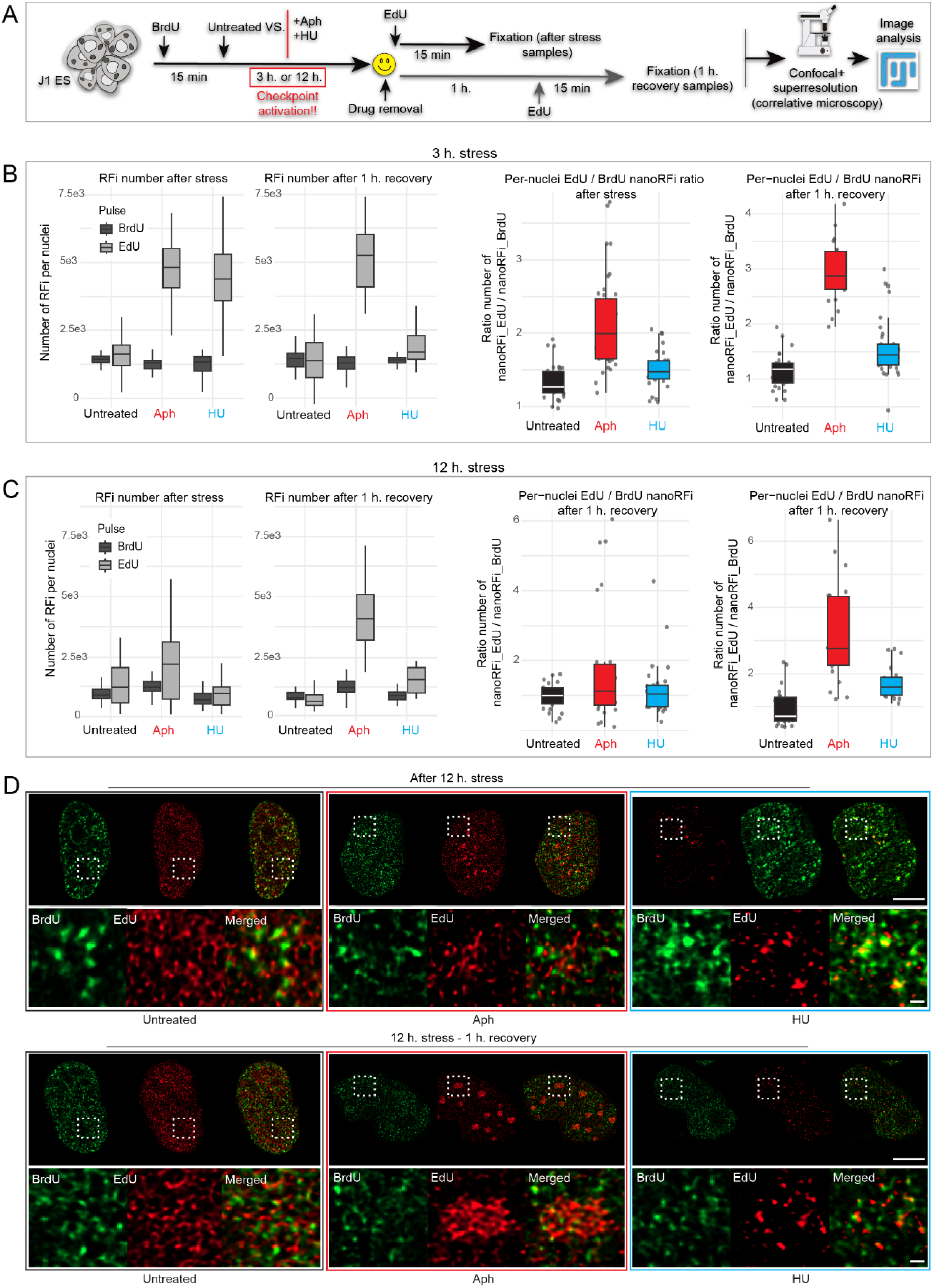
Replication recovery following prolonged stress treatment that induces intra-S checkpoint activation. (A) 3-12 hours treatment pipeline for pulse-chase-pulse experiments. mESCs were pulsed with BrdU for 15 minutes, then treated with 50 µM aphidicolin or 10 mM hydroxyurea for either 3 hours or 12 hours. Following drug removal, cells were pulsed with EdU for 15 minutes either immediately or after 1 hour of recovery, fixed, stained, and analyzed by confocal and superresolution microscopy. (B) Quantification of replication foci (RFi) per nucleus (top), as well as per-nucleus EdU/BrdU nanoRFi ratios (bottom) for 3-hour stress treatments. Left: EdU pulse immediately after stress removal; right: EdU pulse 1 hour after stress removal. Each box plot represents at least 20 cells total from 3 independent replicates. (C) Same quantifications as shown in (B) for 12-hour stress treatments. (D) Representative superresolution images of mESC nuclei showing BrdU-labelled foci (red) before and EdU-labelled foci (green) after 12 hours of stress treatment, either including or excluding 1 hour of recovery. Scale bar: 5 µm. Two independent replicates were performed, including 74-86 total cells per condition.

Furthermore, representative superresolution images of 12-hour treatments show similar patterns to those of 1 hour, with a low level of overlap between after-stress replicons and stalled forks (Figure 7D). The recovery of replication synchrony after 12 hours also showed a trend similar to that without checkpoint activation, with a higher fraction of single replication domains (only 1 nanoRFi) after stress removal, which then decreased at 1 hour of recovery time (Figure S10A). Quantifying colocalization between after-stress nanoRFi (EdU) and nanoRFi before stress (BrdU-labeled stalled forks), we can infer that 66-62% and 64-56% of replicons after 3 hours of treatment do not overlap with stalled forks, and therefore represent newly fired origins, while the rest were found to overlap and thus could represent re-firing stalled forks. Similar percentages were observed for 12-hour stress treatments (Figure S10B).

In summary, 3D mapping of pre- and post-stress replicons shows that stress recovery dynamics in the absence of checkpoint activation ultimately do not change after checkpoint signaling activation and/or extended stress timing. This may reflect an essential pathway or fundamental mechanism orchestrating DNA synthesis restart by activation of dormant origins near stalled forks.

## CONCLUSIONS AND OUTLOOK

In this study, we provide a high-resolution, single-cell view of the replisome’s immediate response to replication stress, before and beyond intra-S checkpoint activation (summarized in Figure 8). By combining live-cell imaging of endogenously tagged replication factors with super-resolution microscopy, structural modeling, and pulse-stress-pulse experiments, we uncovered a sequence of events that reshapes the current view of how cells cope with stalled forks. Upon stress induction with aphidicolin or hydroxyurea, the CMG helicase remains stably loaded and shows increased activating phosphorylation (Mcm2pS108, -pS41, -pS27), while PCNA dissociates concomitantly with 9-1-1 clamp loading, which interacts with the bound PolA-Prim1 complexes, as we have shown both computationally and empirically. Strikingly, all 3 replicative polymerases accumulate at replication sites during stress, and primase remains catalytically active, generating DNA/RNA hybrids despite the absence of detectable DNA synthesis. CDK inhibition abolishes both the stress-associated MCM phosphorylation and the polymerase accumulation, demonstrating that these events are driven by *de novo* firing of dormant origins in the vicinity of stalled forks rather than by reinforcement of pre-existing forks.

**Figure 8.**
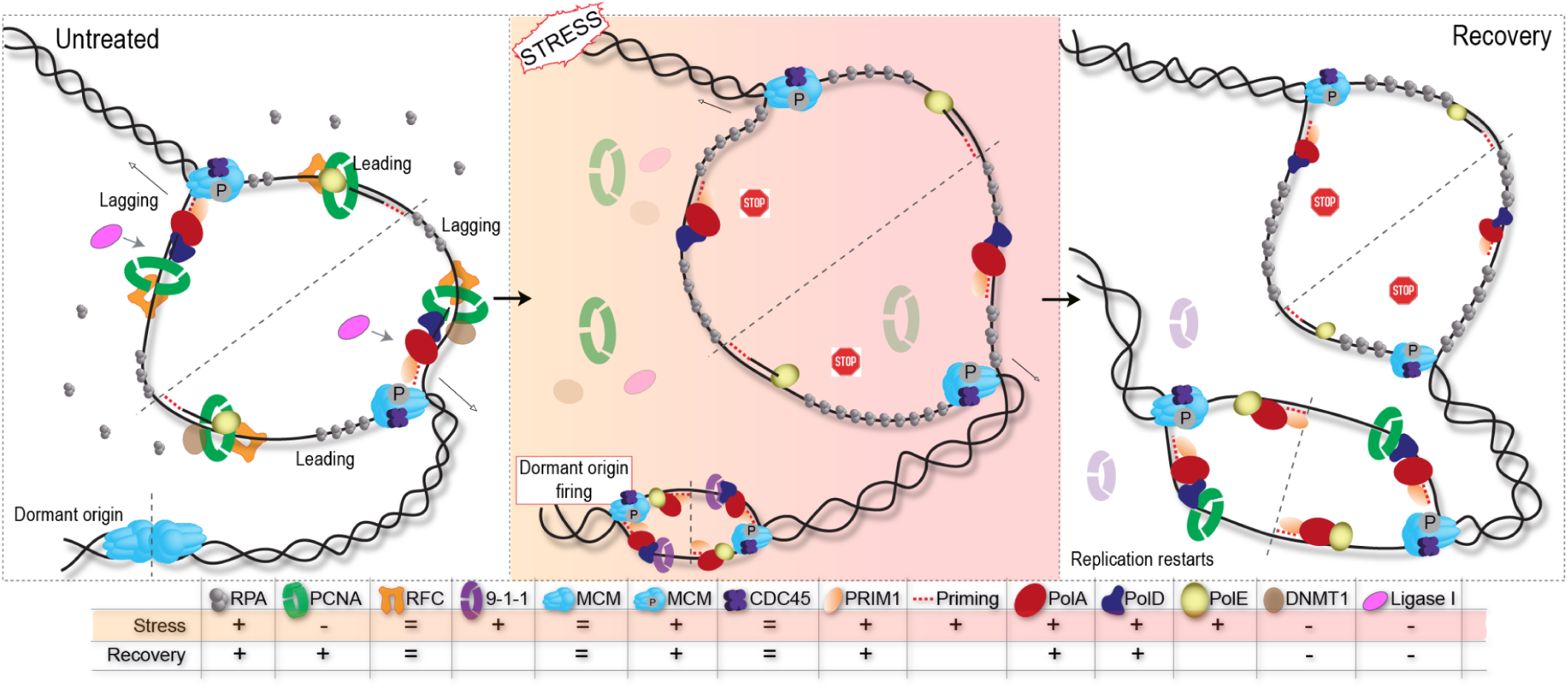
Proposed model of proximal dormant origin activation in response to replication stress. Unperturbed replication (untreated): active replication bubbles, characterized by phosphorylated CMG helicase and replicative polymerases (PolA-PRIM1, PolD, PolE) along with PCNA, expand bidirectionally along the DNA. Inactive MCM complexes remain dormant at nearby licensed origins. Replication stress: polymerase activity at the original fork is blocked while the CMG helicase continues unwinding DNA, generating excess RPA-coated ssDNA. Dormant origins in the vicinity of stalled forks are activated through MCM phosphorylation, leading to replication bubble formation and the recruitment of replicative polymerases and the 9-1-1 alternative clamp, as well as synthesis of RNA primers by PRIM1. Stress recovery: DNA synthesis resumes at the newly primed proximal origins, leading to an increase in observable nanoRFi adjacent to the original stalled forks, which predominantly fail to restart replication themselves.

Building on these observations, our spatial mapping of replicons before and after stress reveals that, upon stress removal, DNA synthesis preferentially restarts at newly fired origins located adjacent to, but largely non-overlapping with, the previously stalled forks. This pattern is conserved between mouse embryonic stem cells and human HeLa cells, and is maintained even after prolonged stress that triggers full Chk1-mediated checkpoint activation, suggesting that proximal dormant origin firing is a fundamental, checkpoint-independent feature of the replisome stress response. Together, these data support a model in which the helicase first unwinds DNA at adjacent dormant origins, PolA-PRIM1 loads and interacts with the 9-1-1 clamp, RNA primers are synthesized and retained while polymerases remain stalled, and, upon stress relief, DNA synthesis resumes from these primed proximal origins. The initial recovery is dominated by single replicons, but synchrony within replication domains is re-established within 1 hour, consistent with a domino-like propagation of firing to neighboring origins.

Several questions remain open and define the direction of our future work. The fate of the originally stalled forks - whether they eventually resume synthesis, are passively rescued by converging forks from neighboring origins, or are processed as substrates for repair - could not be resolved here and will require single-molecule or DNA fiber approaches in combination with the live-cell tools developed in this study. The molecular basis of the PCNA to 9-1-1 clamp switch, including the signals that trigger PCNA unloading and the structural determinants of polymerase retention, also warrants direct biochemical and structural validation. Furthermore, the stress response and loading dynamics of other regulatory replisome factor complexes like Treslin, MTBP, and TopBP1 at stalled forks should be investigated. In summary, the kinetic maps generated of the replisome under stress that were generated in this study uncovered the firing of new origins as a compensatory mechanism to overcome replication stress.

## MATERIALS AND METHODS

### Cell culture, transfection, and treatments

All cells were regularly tested for mycoplasma using PCR and are deemed free of contamination. All cells used in the study were authenticated via ATCC authentication services. mESC J1 (Stevens, 1973) were cultured in Dulbecco’s modified Eagle’s medium (DMEM) high glucose (Cat. No.: D6429, Sigma-Aldrich Chemie GmbH, Steinheim, Germany) supplemented with 16% fetal calf serum (FCS) (Cat. No.: FBS 12 A, Capricorn scientific, Germany), 1× non-essential amino acids (Cat. No.: M7145, Sigma-Aldrich Chemie GmbH, Steinheim, Germany), 1× penicillin/streptomycin (Pen/Strep) (Cat. No.: P4333, Sigma-Aldrich Chemie GmbH, Steinheim, Germany), 1× L-glutamine (Cat. No.: G7513, Sigma-Aldrich Chemie GmbH, Steinheim, Germany), 0.1 mM betamercaptoethanol (Cat. No.: 4227, Carl Roth, Karlsruhe, Germany), 1000 U/ml recombinant mouse LIF (Cat. No.: ESG1107, Merck, Steinheim, Germany) and 2i (PD032591 and CHIR99021 (Cat. Nos.: 1408 and 1386 respectively, Axon Medchem, Netherlands)) on gelatin-coated culture dishes (0.2% gelatin; Cat. No.:G2500, Sigma-Aldrich Chemie GmbH, Steinheim, Germany). Cells were split every 2-4 days. Human cervical cancer cell line HeLa Kyoto provided by Jan Ellenberg (EMBL, Heidelberg, Germany) (Erfle et al., 2007) were maintained in Dulbecco’s modified Eagle medium high glucose (Cat.No.: D6429, Sigma- Aldrich Chemie GmbH, Steinheim, Germany) supplemented with 10% fetal calf serum, 1× glutamine (Cat.No.: G7513, Sigma- Aldrich, St Louis, MO, USA) and 1 µM gentamicin (Cat.No.: G1397, Sigma- Aldrich, St Louis, MO, USA) in a humidified atmosphere with 5% CO2 at 37 °C. Transient transfections were performed using the Neon transfection system (Cat. No.: N10096, Thermo Fisher) in µ-dish chambers (Ibidi), after which cells were maintained at 37 °C with an atmosphere of 5% CO_2_ using an Okolab environmental chamber. In order to block DNA replication, hydroxyurea (HU) (Cat.No.: H8627, Sigma-Aldrich, Steinheim, Germany) or aphidicolin (Aph) (Cat. No.: A0781, Sigma-Aldrich, Steinheim, Germany) was added to the media at final concentrations of 10 mM and 50 µM, respectively. Cells were subsequently examined for periods of 15 minutes following drug treatment. Additionally, the DNA synthesis inhibition was confirmed by labeling the cells with 10 μM EdU (5-ethynyl-2’-deoxyuridine) for 10 minutes and detecting the DNA synthesis. For broad-spectrum CDK inhibition, roscovitine (Cat. No.: 557360-1, Merck, Steinheim, Germany) was used during 1-hour stress treatments (HU or Aph), at a concentration of 5 μM. The expected effect of roscovitine on cell cycle progression at the concentration used was confirmed by flow cytometry.

### Materials for cell line generation and expression constructs

Several targets were fluorescently tagged with EGFP using CRISPR-Cas9 genome engineering in the mouse embryonic stem cell line J1 (mESCJ1). All cell line characteristics are summarized in Supplementary Table 2. Specific gRNAs (Supplementary Table 3, Extended data 1) were cloned into a puromycin-selectable vector expressing both SpCas9 and gRNA (pSpCas9(BB)-2A-Puro (PX459) V2.0 was a gift from Feng Zhang (Addgene plasmid # 62988) (Ran et al., 2013). CRISPR gRNAs were designed using http://crispr.mit.edu/. Cells were transfected with Cas9-gRNA vectors and the homologous recombination template (HRT), including GFP-coding sequences and a linker, using the Neon electroporation system (Cat. No: N10096, Thermo Fisher) according to the manufacturer’s instructions. For CRISPR-Cas9-mediated Homology-Directed Repair (HDR), approximately 3·10^6^ cells were transfected in a volume of 100 μl using 2.5 μg of gRNA/Cas9 plasmids and 20 μg of the homologous recombination template (Supplementary Table 3). The cells were grown for 2 days in gelatin-coated culture plates containing ES medium with 10 μM L755507 (Cat. No.: SML1362, Merck, Steinheim, Germany) to promote homology-directed repair. The medium was then replaced with fresh ES medium containing 1 μg/ml puromycin (Cat.No.: ant-pr-1, Invitrogen) for 48 hours to eliminate cells that were not transfected. The cells were cultured for 7-10 days until individual clones could be picked from plates seeded at low confluency. These clones were then cultured separately, and their fluorescence was examined periodically to narrow down the selection. The most promising clones were further characterized via Western blots and immunofluorescence staining using antibodies that targeted the tagged protein as well as GFP. The clones were further verified by isolating genomic DNA, and Sanger sequencing was performed using the GFP primers to verify the insertion at the target locus. These experiments confirmed that the tagged protein co-localized with the endogenous protein and was present at sites of active replication (Extended data 1 and Figure S3).

To facilitate the generation of triple-labelled stem cell lines for time-lapse microscopy, a plasmid was established allowing the insertion of fluorescently tagged RPA and PCNA into cells using the Sleeping Beauty transposon system. Fragments encoding the desired plasmid backbone, the two fluorescently tagged replication factors, and the internal ribosome entry site were amplified from preexisting plasmids via polymerase chain reaction, during which overlapping regions of 20 base pairs were added to both ends of all fragments to facilitate cloning. The fragments were purified using the Nucleospin PCR and Gel Clean-up kit (Cat.No.: 740609.50, Macherey-Nagel, Düren, Germany) and assembled in one hour at 50 °C using NEBuilder HiFi DNA assembly mix (Cat.No.: E2621S).

Using the previously established GFP-tagged mESCJ1 lines as a basis, approximately 10^6^ cells were transfected in a volume of 100 μl using 15 μg of the vector carrying fluorescently tagged RPA and PCNA (pc5128) and 5 μg of Sleeping Beauty transposase plasmid (pc5129). After 2 days, the medium was supplemented with 1-2 μg/ml puromycin (Cat.code: Ant-pr-1, InvivoGen) until colonies with low or missing RFP and miRFP670 signals (representing tagged RPA and PCNA, respectively) were no longer present upon 24 hours of treatment with 2 μg/m doxycycline (Cat. No.: D9891-1G, Merck, Steinheim, Germany).

### DNA replication and (immuno)fluorescence

Cells were seeded on gelatin-coated glass coverslips. To label sites of DNA synthesis/replication before stress (BS) and after stress (AS), cells were grown for 10 or 15 minutes in a medium supplemented with modified nucleosides (EdU (5-ethynyl-2’-deoxyuridine, Cat. No.: 7845.4, Carl Roth, Karlsruhe, Germany) or BrdU (5-bromo-2-deoxyuridine, Cat. No.: 600-401-C29, Invitrogen, Carlsbad, USA)) at a concentration of 20 µM for the first pulse and 100 µM for the second pulse after stress. For most of the experiments, BrdU was used for the first labelling and EdU for the second. Both BrdU and EdU were proven to be exchangeable, and both were used before and after stress, yielding similar results. Incubation with EdU or BrdU was followed by washing with warm medium before stress treatment or with cold 1x PBS before fixation in 3.7% formaldehyde (Cat. No.: F8775, Sigma-Aldrich Chemie GmbH, Steinheim, Germany) in 1x PBS for 10 minutes. For pre-extraction of MCM proteins, samples were extracted before fixation following the protocol from (Mašata et al., 2011) to eliminate soluble fractions and ensure the detection of chromatin-loaded DNA helicase. Briefly, cells were extracted with 0.1% Triton X-100 in CSK buffer (10 mM Pipes-KOH, pH 7.0, 100 mM NaCl, 300 mM sucrose, 3 mM MgCl2) on ice for 5 minutes. Then, the extraction solution was carefully replaced with 2% formaldehyde in the CSK buffer for 30 minutes. For the other replisome factors, cells were extracted with 10 mM Tris-HCl (pH 7.5), 2.5 mM MgCl2, 0.5% Nonidet P40 substitute (IGEPAL CA-630), and 1 mM PMSF in ddH2O for 2 minutes on ice. Then, the extraction buffer was replaced with 4% formaldehyde in 1xPBS. After fixation and 3 washing steps with PBS-T (1x PBS, 0.01% Tween-20), cells were permeabilized with 0.5% Triton X-100 in 1x PBS for 20 minutes. Only for DNA-RNA hybrid immunostaining (anti-S9.6), incubation with RNaseH (1U/μL, Cat. No.: M0297S, New England Biolabs) was performed for 1 hour at 37°C. Only for PCNA immunostaining, after permeabilization, cells were incubated in ice-cold 88% methanol in ddH2O (v/v) for 5 minutes for antigen retrieval, and washed again. Before incubation with primary antibodies, blocking was performed for 40 minutes in 2% bovine serum albumin in 1× PBS at 37 °C in a humid chamber. Primary antibody incubation was performed in 2 % BSA in 1xPBS for 1.5 hours at 37 °C. BrdU was detected with a rat anti-BrdU antibody (1:100) diluted in buffer consisting of a 1:1 mixture of blocking and 2× DNase I reaction buffer (60 mM Tris/HCl, pH 8.1, 0.66 mM MgCl2, 1 mM beta-mercaptoethanol) and 25 U/ml DNase I (Cat. No.: D5025, Sigma-Aldrich Chemie GmbH, Steinheim, Germany). DNase I digestion was stopped by washing with PBS–TE (PBS-T with 1 mM EDTA). After incubation with the primary antibodies, cells were washed 3 times with PBS-T, and for the detection of the primary antibodies, cells were incubated with fluorescently tagged secondary antibodies diluted in 2% BSA. After 45 minutes of incubation with the secondary antibody solution at room temperature, cells were washed 3 times with PBS-T. The detection of EdU was performed using the Click-IT assay following the manufacturer’s instructions, preparing a dilution containing 1:200 3-azido-7-hydroxycoumarin, 1:1,000 6-carboxyfluorescein (6-FAM azide), 1:2,000 5/6-sulforhodamine azide, or Eterneon-Red 645 Azide (Cat. No.: 7811, 7806, 7776, and 1Y73.2, respectively, Carl Roth, Karlsruhe, Germany). The samples were incubated with the Click-IT mix for 45 minutes at room temperature, followed by 3 washing steps in PBS-T. Finally, DNA was counterstained with DAPI (4,6-diamidino-2-phenylindole, 10 g/ml, Cat. No.: D27802, Sigma-Aldrich Chemie GmbH, Steinheim, Germany) for 10 minutes and samples were mounted in Mowiol 4-88 (Cat. No.: 81381, Sigma-Aldrich Chemie GmbH, Steinheim, Germany) containing 2.5% DABCO (1,4-diazabicyclo[2.2.2]octane, Cat. No.: D27802, Sigma-Aldrich Chemie GmbH, Steinheim, Germany), or in Vectashield (Cat. No.: VEC-H-1000, Biozol Diagnostica Vertrieb). All relevant information on modified nucleotides is shown in Supplementary Table 4, and primary and secondary antibodies are described in Supplementary Table 5.

### Proximity ligation assay (PLA)

*In situ* interaction between PolA- and PRIM1-DNA/RNA hybrids was quantified by proximity ligation assay using the DuolinkⓇ kit (Cat. No.: DUO92101, Sigma-Aldrich, Steinheim, Germany), following the manufacturer’s instructions and the protocol described in (Derangère et al., 2016), (Arroyo et al., 2024), and (Lin et al., 2015). In this technique, small oligonucleotide probes (+) and (-), conjugated to secondary antibodies, specifically recognize the primary antibodies against the proteins of interest. When the 2 probes are closer than 40 nm, ligation by ligase incubation can occur. This generates circular DNAs that will be amplified by a polymerase incorporating fluorescently labeled nucleotides. Afterward, fluorescent spots can be detected and quantified using microscopy and image analysis, considering each spot an interaction site between the 2 proteins. Briefly, immunostaining was performed as described before, but after incubation with primary antibodies and subsequent washing steps, fluorescent-conjugated secondary antibodies were replated with DuolinkⓇ PLA reagents (*in situ* complementary oligonucleotide probe MINUS (-) and PLUS (+)). Incubation with only one of the primary antibodies (anti-DNA/RNA hybrids) was used as a negative control for the assay. The PLA probes were mixed and diluted (1:5) in antibody diluent (2% BSA in 1x PBS), incubated at room temperature for 20 minutes, and then incubated with the samples of interest for 1 hour at 37 °C in a humid chamber. Then, samples were washed 2 times with washing buffer A (0.01 M Tris, 0.15 M NaCl, 0.05% Tween-20), and the probes were ligated with 2 other circle-forming DNA oligonucleotides by ligation-ligase solution for 30 minutes at 37 °C. After this incubation and washing steps, amplification of the oligonucleotides was performed via the rolling circle by incubation with amplification-polymerase solution (nucleotides and fluorescently labeled oligonucleotides together with polymerase) for 90 minutes at 37 °C. During this incubation, the fluorescent oligonucleotides hybridize into the rolling-circle amplification product, making the signal visible as a fluorescent spot by microscopy. Finally, samples were washed with washing buffer B (0.2 M Tris, 0.1 M NaCl) 2 times x 10 minutes, once in 1xPBS, and DNA was counterstained with DAPI and mounted with Mowiol as described for immunostainings.

### Flow cytometry

After the corresponding treatments, cells were trypsinized, fixed using 70% ethanol, and stored at -20 °C before staining and analysis. For pre-extracted cells, the same extraction protocol described before was used before trypsinization. For immunostaining, cells were centrifuged at 1000xg for 5 minutes and washed twice with 1xPBS, followed by permeabilization with 0.7 % Triton X-100 for 20 minutes at room temperature. Blocking was performed by resuspending the cells in 4 % BSA in 1xPBS for 30 min. Incubation with primary antibodies (Supplementary Table 5) was performed for 3 hours at room temperature, followed by washing and incubation with secondary antibodies for 1 hour. Then, cells were washed and incubated with 50 μg/mL propidium iodide in 1× PBS and 0.3% Triton X-100 supplemented with 100 μg/mL RNaseA for 3 hours, washed again with 1xPBS, and sorted using an S3e Cell sorter (Bio-Rad). Data corresponding to 3 independent replicates were analyzed using FloJo software (https://www.flowjo.com/). Propidium iodide was used to sort the cells by cell cycle stage based on DNA content. Negative control (without primary antibody staining) was used to distinguish positive from negative cell fractions.

### Microscopy

Characteristics of the microscope systems, including laser, filters, and objectives used, are summarized in Supplementary Table 6.

### Live cell microscopy

Super-resolution live cell imaging of fluorescently tagged mESCJ1 was performed using a Zeiss LSM 900 Airyscan system (Zeiss, Germany) equipped with a C Plan- Apochromat 63x oil objective (NA: 1.4) and an Axiocam 820 mono SONY IMX541 CMOS camera with an Airyscan detector for high-speed stack acquisition at a lateral resolution of approximately 120 nm. To induce the expression of RFP-RPA and miRFP670-PCNA in triple-tagged cell lines, the culture medium was supplemented with 2 µg/ml doxycycline 24 hours before imaging. For the duration of the experiment, the microscope’s incubation chamber was kept at 37 °C and 5% CO_2_. The fluorophores GFP, RFP, and miRFP670 were excited using the 470 nm, 561 nm, and 640 nm laser lines, respectively, and detected in ranges of 490-575 nm, 565-620 nm, and 650-700 nm. For each sample, 10-20 regions containing S-phase cells were selected based on the replication patterns that could be observed via PCNA fluorescence, and a z-stack of 34 image slices with an interval of 0.15 µm was acquired for each region. The culture medium was then supplemented with either 50 µM aphidicolin or 10 mM hydroxyurea, after which the selected regions were imaged every 15 minutes for 1 hour in the same manner. The medium was subsequently replaced with a fresh medium containing no replication inhibitors, and the imaging continued in 15-minute intervals for 2 additional hours.

### Superresolution microscopy of fixed cells

Samples were prepared as described in section “DNA replication and (immuno)fluorescence”. 3D z-stack images were acquired with the Zeiss LSM 900 Airyscan system (Zeiss, Germany) described above, using the C Plan- Apochromat 63x oil objective (NA: 1.4) and Axiocam 820 mono SONY IMX541 CMOS camera. The fluorophores were excited using the 470 nm, 561 nm, and 640 nm laser lines, respectively, and detected in ranges of 490-575 nm, 565-620 nm, and 650-700 nm. For each sample, regions containing S-phase cells were selected based on the BrdU/EdU signal, and a z-stack of 40 to 60 image slices with an interval of 0.15 µm was acquired for each region. For correlative microscopy analysis, raw images were taken in Airyscan mode and further processed with Airyscan joint deconvolution to generate superresolved images in Zen software.

### Confocal microscopy

To analyze the accumulation of replisome factors and replication foci in fixed cells by staining, samples were prepared as described above. Confocal images were collected in a Leica TCS SP5II confocal laser scanning microscope (Leica Microsystems, Wetzlar, Germany) equipped with an oil immersion Plan-Apochromat 100×/1.44 NA objective lens (pixel size in XY set to 50 nm, Z-step = 290 nm). The latter microscope was also used for the acquisition of images in the PLA assays.

For characterization of the cell lines, an UltraVIEW VoX spinning disc system (PerkinElmer, UK) on a Nikon Ti microscope (Nikon, Japan) was used, equipped with an objective 100×/1.49 NA CFI Apochromat TIRF oil immersion (voxel size, 0.071 × 0.071 × 0.5–1 μm; Nikon, Tokyo, Japan) and a cooled 14-bit CCD camera.

### Image analysis

All image analysis was performed on Fiji, Volocity, Zeiss software, and R. A list of analysis software and associated plugins/packages is found in Supplementary Table 7. All scripts used for image analysis are uploaded here https://doi.org/10.48328/tudatalib-2251

### Quantification of replisome factors dynamics by live-cell microscopy

Time-lapse data obtained by live-cell superresolution microscopy underwent joint Airyscan deconvolution with 20 iterations of processing in Zeiss Zen 3.9, and the z-stacks were orthogonally projected into two-dimensional images using maximum projection for further analysis in Volocity 6.3. 5 to 10 cells exhibiting a replication pattern in the miRFP670-PCNA channel during the earliest time point were chosen manually for further analysis. Within these cells, a mask of replication foci represented by strong RPA or PCNA signal intensity was generated for each time point. Any foci smaller than 0.05 µm^2^ were omitted from the plots as they largely corresponded to transient imaging artifacts rather than persistent replication sites. Additionally, a mask of the entire cell nucleus was generated based on the nuclear background (soluble) PCNA signal. To account for fluctuations in fluorescence intensity due to photobleaching or slight shifts in the focal plane, the mean signal intensity of the GFP-tagged factor of interest within each replication focus was normalized to the mean nuclear GFP intensity of the corresponding cell at each time point. For visual data representation, the average of the normalized intensity values and the corresponding 95 % confidence interval was calculated at each time point.

### High-throughput image analysis of replisome factors accumulation

Confocal images were processed and analyzed with semi-automatic Fiji scripts as described in (Arroyo et al., 2023) with some modifications. First, we performed nucleus extraction and segmentation from multichannel fluorescence images using a semi-automatic Fiji macro. The macro recursively scanned a user-defined input directory for image files. Then, the user was prompted to manually select nuclei of interest using Fiji’s selection tools and add them to the ROI (Region of Interest) Manager. For each selected ROI, the corresponding nuclear region was duplicated as a full z-stack, signal outside the ROI was cleared, and the cropped nucleus was saved. After nuclei segmentation, quantification of signal intensity within and outside the replication foci was performed. Images were opened using the Bio-Formats Importer, and a maximum intensity z-projection was generated for multi-z stacks. A duplicate of the projected image was converted to 8-bit and subjected to Gaussian blur (sigma = 1) to reduce noise, followed by automated binary thresholding using the Otsu method with dark background correction and hole filling to generate a nuclear mask. The user was then prompted to select the nucleus of interest using the wand tool, and the resulting ROI was added to the ROI Manager. To segment EdU-positive regions, channels were split, and the EdU was used to generate an intensity-averaged image, creating a binary mask for ROI selection. Mean, Sum, standard deviation, minimum, and median fluorescence intensities were then measured across all 4 channels within the EdU-positive selection. To quantify signals in the EdU-negative nuclear area, the EdU-mask selection was cleared from the nuclear-masked image, and measurements were repeated across all 4 channels within the remaining nuclear ROI. Results were exported as a .csv file and further processed using RStudio. Ratio of EdU/EdU-negative regions for all channels was calculated by dividing the sum intensities of EdU-segmented areas by the sum intensities in EdU-negative regions. The obtained ratios for each cell nucleus were normalized by the average ratio of untreated samples. Normalized ratios were plotted.

### Quantification of PLA assay

Proximity ligation assay was analyzed in confocal images using Fiji. Replicating cells in any S-phase substage (EdU positive) and non-replicating cells were selected. First, maximum intensity projections were generated from multicolor Z-stacks, and nuclei were segmented using the DAPI signal. In this step, we obtained nuclear ROIs, which were applied to the channel for the PLA signal as a selection. This removed the background outside the nuclei and the spots corresponding to additional cells in the image. Once the nuclear selection was applied to the ‘PLA channel’, we found the local maxima that corresponded with the fluorescent PLA spots, maintaining the same value of ‘Prominence’ for all samples. In all our data sets, this value was set to >40, and for accurate counting, the function ‘Exclude edge maxima’ was used. The final output of this quantification was the number of maxima/spots per nucleus versus cytoplasm. The number of spots for each condition was plotted using RStudio.

### Quantification of nanoRFi/RFi and correlative microscopy analysis

Supersolution and correlative microscopy analysis were performed as described in (Pradhan et al., 2024), using a set of automated scripts in Fiji. These scripts are available at https://doi.org/10.48328/tudatalib-2251. First, all 16-bit images were converted to 8-bit using the script “ASJD_16_to_8_bit.ijm” before analysis. 3D nuclei were segmented using the DAPI signal to define the nuclear ROI. The 3D stack of the DAPI channel was processed with a Gaussian filter (pixel radius = 2) and normalized (process > enhance contrast > saturated pixels = 0). The nucleus was segmented with 3D nuclei segmentation in the 3Dsuite plugin, a binary image was created, and further processed with dilations, fill holes, and erode (see script “Batch_DAPI_Segmentation_Process_3D.ijm”). This ROI was applied to all the channels to define the ROI using the image calculator, applying “min” between the DAPI mask and the channel of interest (see script “Image_Calculator_Min_Stack.ijm”). All DAPI masks were manually cross-checked for suboptimal DAPI segmentation. For quantification of RFi and nanoRFi corresponding to replicons (3D spot quantification), a combined approach of threshold and cluster-based segmentation was performed for the 3D spot segmentation. First, the 3D stack was processed using a mean filter (pixel radius = 1) and normalized. The FindStackMaxima Macro (https://imagej.nih.gov/ij/macros/FindStackMaxima.txt) was used to find all the local maxima. The corresponding voxels of 3D Maxima were extracted from the processed image using the Image Calculator, which was used as a seed (“Image_Calculator_Min_Stack.ijm”). The seed and processed images were used to segment the foci in 3D spot segmentation (3D suite) using a Gaussian fit to determine the intensity value, used as the threshold to stop the voxel clustering. The segmented objects were further processed with a 3D watershed split (3D suite) to separate the closely clustered RFi/replicons/spots. This was imported into the 3D ROIManager (3D suite) for further quantification and measurements. For 3D spot quantification, the 3D numbering plugin was used (RFI_or_Object_ Quantification_in_ROI_DAPI_MASK.ijm), where the number of 3D spots was measured inside a single nucleus. The RFi distance analysis and features were extracted using the 3D ROIManager, where the first object was the DAPI mask, and the rest of the objects were individual RFi/nanoRFi. All possible combinations of distances (for further analysis) were measured using the “distances” function. The volume of the individual foci was measured using the Measure 3D function. For correlative microscopy analysis, confocal versus processed images (Airyscan Joint Deconvolution) were used. Segmented replication foci in both resolutions were quantified as described above and compared with each other.

### NanoRFi within RFi analysis of replication synchrony by correlative microscopy

To quantify replication synchrony, correlative image analysis was performed as described above. The EdU or BrdU-labeled replication foci were segmented in both confocal and superresolved images. Then, the masked replication foci in confocal resolution (RFi, corresponding to replication domains) were used as individual masks for quantification of nanoRFi (single replicons) in superresolved images. This quantification yielded the number of nanoRFi within individual RFi for each segmented RFi in a single cell nucleus. Then, the replicons were further classified as single (1 nanoRFi within 1 RFi), or multiple, with 2, 3, 4, 5, and more than 5 nanoRFi within a single RFi. The fraction of each of these categories per condition was plotted using RStudio.

### Spatial mapping of stress recovery response by correlative microscopy

The correlative microscopy approach described before was used to map the location of stress response and replication restart after stress removal. Replication labeling before stress (BrdU or EdU) and after stress removal (EdU or BrdU) was performed, in addition to PRIM1 or PCNA immunofluorescence. First, confocal and superresolved images were used to quantify the number of nanoRFi (after stress removal, second labelling) within RFi before stress. Segmented RFi were used as a mask to quantify the number and the location of nanoRFi during stress recovery, yielding the fraction of pre-stress replication domains (RFi) containing 0, 1, or >1 nanoRFi. Next, superresolved images of pre-stress and post-stress labelled nanoRFi were used to calculate the degree of overlap between replicons, using the segmented nanoRFi labelled after stress removal to quantify the number of pre-stress nanoRFi (corresponding to stalled forks) within them. This analysis yielded the fraction of stalled forks overlapping with 0, 1, or >1 of the new replicons after stress removal. The same approach was used to map the location of new origins priming, 9-1-1 loading during stress, and PCNA loading after stress removal in relation to stalled forks and/or new fired origins.

### 3D colocalization and distance analysis with DiAna

Three-dimensional colocalization between replication foci labelled before stress (BrdU pulse) and after stress removal (EdU pulse) was performed using the DiAna plugin for Fiji (Gilles et al., 2017). Prior to analysis, all 16-bit confocal z-stacks were converted to 8-bit using the script “ASJD_16_to_8_bit.ijm”, and nuclear regions of interest (ROIs) were defined based on DAPI signal as described in the previous section (see script “Batch_DAPI_Segmentation_Process_3D.ijm”). The DAPI mask was applied to both BrdU and EdU channels using the image calculator function with the “min” operation to restrict the analysis to intranuclear signal (“Image_Calculator_Min_Stack.ijm”). For each nucleus, BrdU and EdU foci were independently segmented in 3D using the DiAna segmentation module. A combined threshold and spot-based segmentation approach was applied, with local maxima detection used as seed points for iterative voxel clustering. Segmentation parameters (minimum and maximum object volumes, threshold values) were kept consistent across all samples. Closely clustered foci were further separated using 3D watershed splitting. Segmented BrdU and EdU objects were then passed to the DiAna analysis module for pairwise colocalization quantification. The following parameters were extracted for each pair of BrdU–EdU objects within the same nucleus: center-to-center distance between objects in 3D (in µm), volume of each individual object (in voxels and µm³), the absolute volume of overlap between paired objects, and the fraction of each object’s volume that overlapped with its partner. Object pairs were considered “touching” when their segmented volumes shared at least 1 voxel (i.e., overlap volume > 0). For each cell, we computed the total number of BrdU and EdU foci, the percentage of EdU foci touching at least 1 BrdU focus and vice versa, the mean fraction of overlapping volume for touching pairs, and the center-to-center distance distribution for touching pairs. Non-touching EdU foci were classified as newly fired origins, while touching pairs were interpreted as either re-firing stalled forks or spatially adjacent neighboring origins, depending on their fractional overlap. As a positive control for maximum colocalization, samples in which BrdU and EdU were co-pulsed simultaneously (co-pulse condition) were processed in parallel and used as a reference for the expected upper limit of overlap given the spatial resolution of confocal microscopy. Untreated samples subjected to 2 sequential nucleotide pulses separated by 1 hour of unperturbed replication were used as a control for the degree of overlap expected from natural replication progression in the absence of stress. All DiAna output tables were exported as CSV files and further processed in RStudio. Scripts used for DiAna batch processing and downstream R analysis are available in the TUdatalib repository (https://doi.org/10.48328/tudatalib-2251).

### Kolmogorov-Smirnov spatial randomness analysis

To assess whether the three-dimensional distribution of replication nano-foci (nanoRFi) within nuclei deviated from complete spatial randomness (CSR), we applied the one-sample Kolmogorov-Smirnov (KS) test to the nearest-neighbor distance distributions of segmented foci. NanoRFi were segmented in 3D from superresolution z-stacks as described in the previous section, and their centroid coordinates (x, y, z, in µm) were extracted per nucleus using the 3D ROIManager (3D suite) within nuclear ROIs defined by DAPI segmentation. The coordinates of each nucleus were exported as an individual CSV file for downstream analysis in RStudio. For each nucleus, the nearest-neighbor distance (*r*) was computed for every nanoRFi as the minimum Euclidean distance to any other nanoRFi, using the *nndist* function from the *spatstat.geom* package applied to the matrix of 3D coordinates. From these per-cell distance distributions, the empirical cumulative distribution function (ECDF) was calculated, representing, for any distance *r*, the fraction of nanoRFi whose nearest neighbor was located within that distance. The ECDF was then compared to the theoretical cumulative distribution function expected under complete spatial randomness in 3 dimensions, defined as:

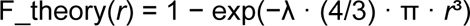

where λ is the point density, calculated per nucleus as the number of segmented nanoRFi divided by the volume of the three-dimensional axis-aligned bounding box enclosing all foci within that nucleus, computed as (x_max − x_min) · (y_max − y_min) · (z_max − z_min). This theoretical distribution assumes that points are independently and uniformly distributed throughout the available volume. The one-sample KS test was applied to compare the ECDF with the theoretical CSR distribution using base R’s ks.test function with F_theory passed as the reference distribution, yielding 2 parameters: the KS statistic (KS-D), defined as the maximum vertical distance between the ECDF and the theoretical CDF, which quantifies the magnitude of deviation from randomness; and the associated p-value (KS-p), representing the probability that the observed distribution could have arisen under CSR. ECDFs positioned above the theoretical curve indicate spatial clustering (nearest neighbors closer than expected by chance), whereas ECDFs below the theoretical curve indicate spatial dispersion (nearest neighbors farther than expected by chance). This analysis was performed separately on BrdU-labelled nanoRFi (pre-stress) and EdU-labelled nanoRFi (immediately after stress removal or after 1 hour of recovery), in both untreated and stress-treated samples (aphidicolin or hydroxyurea). Plots showing ECDF curves overlaid with theoretical CSR curves were generated using *ggplot2*. Scripts for KS analysis and plotting are available in the TUdatalib repository (https://doi.org/10.48328/tudatalib-2251).

### Protein samples preparation, SDS PAGE, and Western blot

For Western blot analysis, cells were trypsinized, counted to 5×10^5^ cells per lane, and lysed for 15 minutes on ice in 800 mM NaCl containing lysis buffer (20 mM Tris-HCl (pH 8), 1.5 mM MgCl2, 0.2 mM EDTA, 0.4% NP-40, and protease inhibitors). The lysate was drawn 10 times through a 21-G needle, incubated on ice for 25 more minutes, and cleared by centrifugation for 15 minutes at 13,000 xg and 4 °C. This lysate was diluted in 6x SDS loading buffer (200 mM Tris/HCl, pH 6.8, 400 mM DTT, 8% SDS, 0.4% bromophenol blue, and 40% glycerol) to 1x and heated to 95 °C for 5 minutes. SDS-PAGE and Western blotting were performed as described in (Mortusewicz et al., 2006), with SDS-PAGE (sodium dodecyl sulfate–polyacrylamide) gel concentrations ranging from 6 % to 12 % depending on the mass of the proteins of interest. The protein was subsequently transferred onto a nitrocellulose membrane (GE Healthcare, München, Germany). Blocking of membranes was performed for 1 hour in 3% low-fat milk in 1x PBS, followed by incubation with primary antibodies diluted in blocking buffer overnight at 4 °C. The membrane was incubated with the respective secondary antibodies after washing using 1xPBS supplemented with 0.01% Tween-20. Horseradish peroxidase (HRP) conjugated sheep anti-mouse IgG (Amersham Pharmacia Biotech, United Kingdom), goat anti-rabbit IgG (Sigma-Aldrich, USA), and goat anti-rat IgG (The Jackson Laboratory, USA) were used (1:5000). Alternatively, incubation with fluorescently tagged secondary antibodies was performed using Cy5-conjugated donkey anti-mouse IgG (H+L), Cy3-conjugated donkey anti-mouse IgG (H+L), Cy5-conjugated donkey anti-rabbit IgG (H+L), and Cy3-conjugated donkey anti-rabbit IgG (H+L) (1:2000; The Jackson Laboratory, Bar Harbor, ME, USA). Characteristics of all primary and secondary antibodies, as well as dilutions used, are described in Supplementary Table 5. For imaging, the Amersham AI600 Imager was used (GE Healthcare, Chicago, II, USA). Cutouts of the membranes were made in some cases for better composition of the figures. Uncropped and unprocessed scans for all the blots are available, including replicates, and are provided with the data sets uploaded to TUdatalib (https://doi.org/10.48328/tudatalib-2251).

### Computational modeling with AlphaFold 3 and AlphaFold 2-Multimer

Protein complexes interaction modeling was performed with AlphaFold as described in (Hastert et al., 2025). The AlphaFold3 web interface was used to screen protein-protein-RNA interactions (https://alphafoldserver.com/) (Abramson et al., 2024). To rank the predictions, both pTM and ipTM scores were used. The predicted template modeling score (pTM) and the interface-predicted template modeling (ipTM) are measures of the accuracy of the entire structure (Xu and Zhang, 2010; Zhang and Skolnick, 2004). As a guide, pTM scores above 0.5 indicate that the overall predicted fold for the complex might be similar to the true structure. Complementary to pTMs, the ipTM score measures the interface and the accuracy of the predicted relative positions of the subunits within the complex. For ipTM scores, values higher than 0.8 represent high-quality predictions with high confidence, while values below 0.4 suggest a likely failed prediction. PAE and pLDDT were also taken into consideration after a close inspection of the models. PAE plots for full-length protein predictions and pre-existing models in the 3D structures databases (AlphaFoldDB) were used. A local installation of AlphaFold-Multimer 2.3.2 was used to perform selected structural modeling with AMBER relaxation (Evans et al., 2021). 5 predictions were generated per model, and the prediction with the highest model confidence was used for verification of AlphaFold 3 predictions. All protein sequences were extracted from UniProt (UniProt Consortium, 2023). Data analysis and plotting were done with Python (pandas and numpy packages). Protein structure images were generated with ChimeraX (Pettersen et al., 2021). All the structural models, generated “.pbd” files, and associated data, including PAE values, “.json”, and scores, are deposited and available at the repository TUdatalib (https://doi.org/10.48328/tudatalib-2251).

### Statistics and data availability

RStudio (versions V1.2.5033 and V2023.03.1-446, https://rstudio.com/) and MicrosoftⓇ ExcelⓇ for Mac 2011 (Version 14.7.7) were used for data visualization, analysis, statistics, and plotting (unless stated otherwise). In all barplots, the median value of the distribution is shown, with the whiskers representing the standard deviation (95% confidence interval). For graphs showing box plots, in all cases, the box represents the distribution of 50% of the data, starting at the first quartile (25%) and ending at the third (75%), with the line inside the box representing the median value and with whiskers representing the upper and lower quartiles not included in the box. Outliers are defined as 1.5 times the interquartile range and are excluded from most of the plots. Statistical comparisons were applied when considered necessary using an independent two-group comparison test: Wilcoxon-Mann-Whitney or One-Way ANOVA. Related to statistical significance, not significant (n.s.), is assigned to p-values > or equal to 0.05; 1 star (*) is assigned to p-values < 0.05 and > or equal to 0.005; 2 stars (**) is assigned to values < 0.005 and > or equal to 0.0005; 3 stars (***) is given for values < 0.0005; 4 stars (****) is given for values < 0.00005. This information is located in the plots between the top of the 2 samples subjected to comparison. Statistical values (number (#) of cells (N), mean, median, standard deviation (SD), and p-values are summarized in the corresponding figure legends.

All the data supporting this study are available in the repository TUdatalib, registered with the following DOI: https://doi.org/10.48328/tudatalib-2251

## ACKNOWLEDGMENTS

We are indebted to Anne Lehmkuhl and Manuela Milden-Appel for excellent technical help, Alexander Rapp and Sunil K. Pradhan for advice on imaging and image analysis, and Conrad E. Knies and Selina D. Buchwald for their help with cell line generation and immunostaining analysis. We thank Katja Luck for initial help with AlphaFold modeling, and Johannes Walter and Chunlong Chen for discussions. L.S., M.K.P. and M.K. have been members of the Graduate School of Life Science Engineering of the TU Darmstadt.

Molecular graphics and analyses performed with UCSF ChimeraX, developed by the Resource for Biocomputing, Visualization, and Informatics at the University of California, San Francisco.

## DATA AVAILABILITY STATEMENT

Renewable biological materials will be made available upon request from the corresponding author M. Cristina Cardoso. All our data sets have been deposited and are available at https://doi.org/10.48328/tudatalib-2251

## AUTHORS CONTRIBUTIONS

LS: Data curation, formal analysis, investigation, methodology, visualization, writing original draft, writing - review & editing

MKP: Formal analysis, investigation, methodology, visualization, writing - review & editing

MK: Investigation, methodology, writing - review & editing

PP: Investigation, methodology, writing - review & editing

MA: Data curation, formal analysis, investigation, methodology, visualization, writing original draft, project administration, writing - review & editing

MCC: Funding acquisition, project administration, writing - review & editing

## SUPPLEMENTARY INFORMATION

### Supplementary figures

**Figure S1.**
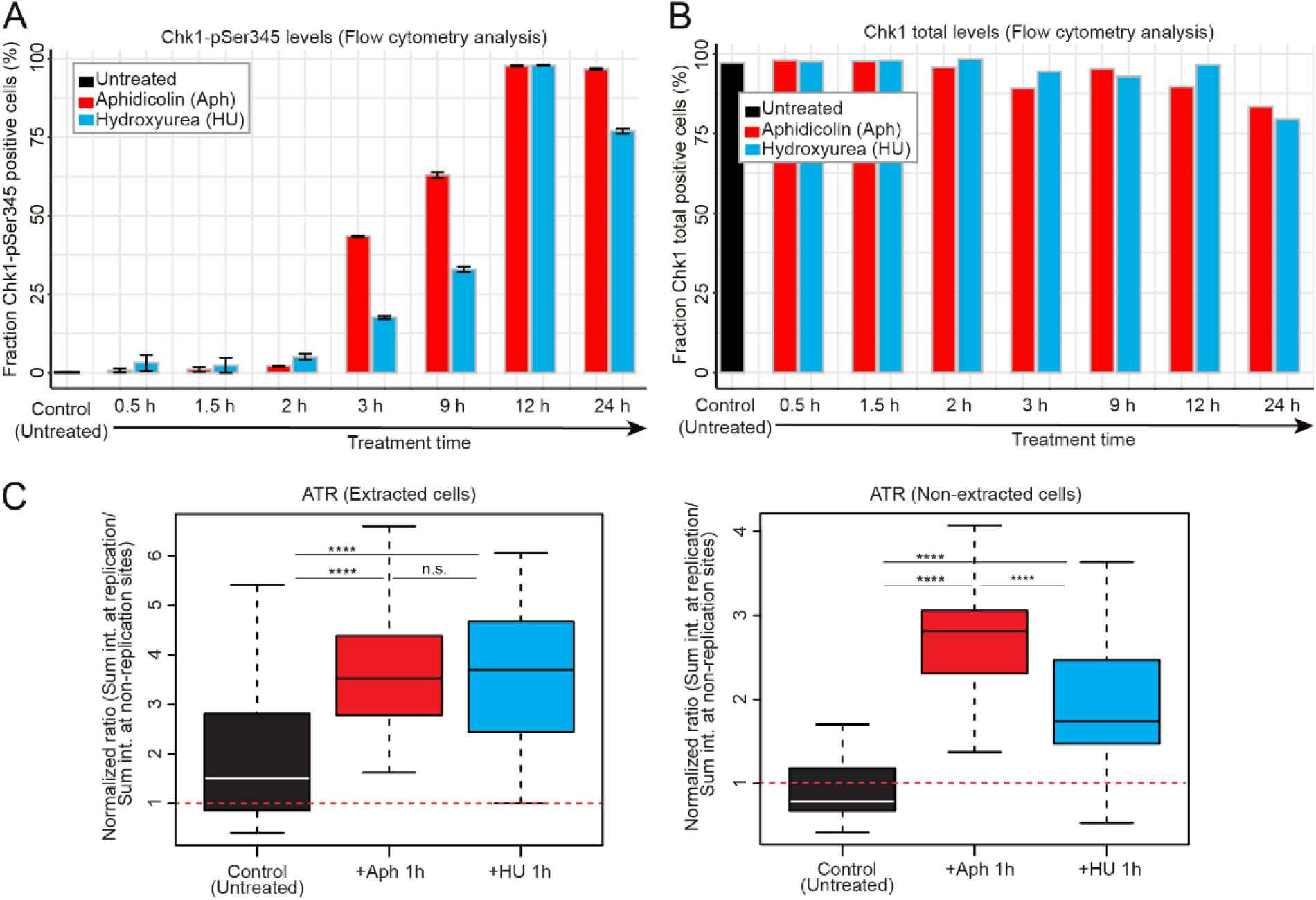
Time course of different stages of the intra-S checkpoint activation under replication stress in mESCs. (A) Percentage of cells positive for Chk1-pSer345, measured by flow cytometry over 24 hours of aphidicolin (red) or hydroxyurea (blue) treatment. (B) Total Chk1 levels over the same 24-hour period, measured by flow cytometry. For (A) and (B), 2 independent replicates were performed, quantifying at least 5000 cells in total. The whiskers represent the standard deviation with a 95% confidence interval. (C) Quantification of total nuclear ATR levels (left) and chromatin-bound ATR levels (right) in mESCs after 1 hour of aphidicolin or hydroxyurea treatment compared to untreated controls. Cells were immunostained against ATR and imaged using confocal microscopy. Nuclei were segmented based on the DAPI signal, and sum intensities for the ATR channel were normalized to the sum intensity of DAPI and to the average of untreated cells. Each box plot represents at least 20 cells from 2 independent replicates. Statistical significance was tested with a paired two-sample Wilcoxon test (n.s. is given for p-values ≥ 0.05; 4 stars (****) are given for values < 0.00005).

**Figure S2.**
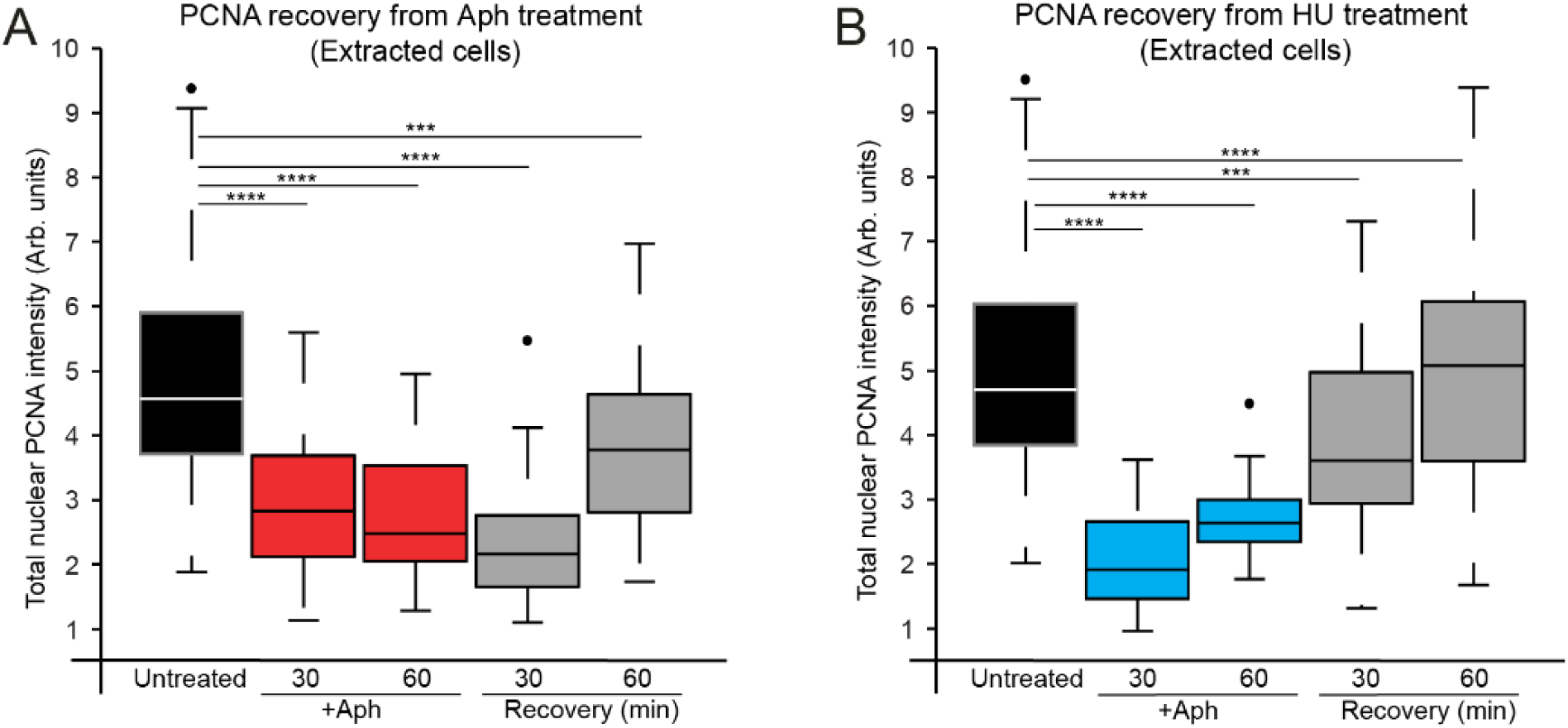
Measurements of chromatin-bound nuclear PCNA in S-phase cells during stress induction and subsequent recovery in fixed, pre-extracted cells. S-phase cells were identified based on an EdU pulse performed before stress induction. Each box plot represents 2 replicates, consisting of 50–60 cells. Bound nuclear PCNA levels were quantified before, during, and after aphidicolin treatment (A) or hydroxyurea treatment (B). Each box plot represents 50% of the data, starting in the first quartile (25%) and ending in the third (75%). The line inside represents the median. The whiskers represent the upper and lower quartiles. Statistical significance was tested with a paired two-sample Wilcoxon test (three stars (***) are given for values < 0.0005 and ≥ 0.00005; 4 stars (****) are given for values < 0.00005).

**Figure S3.**
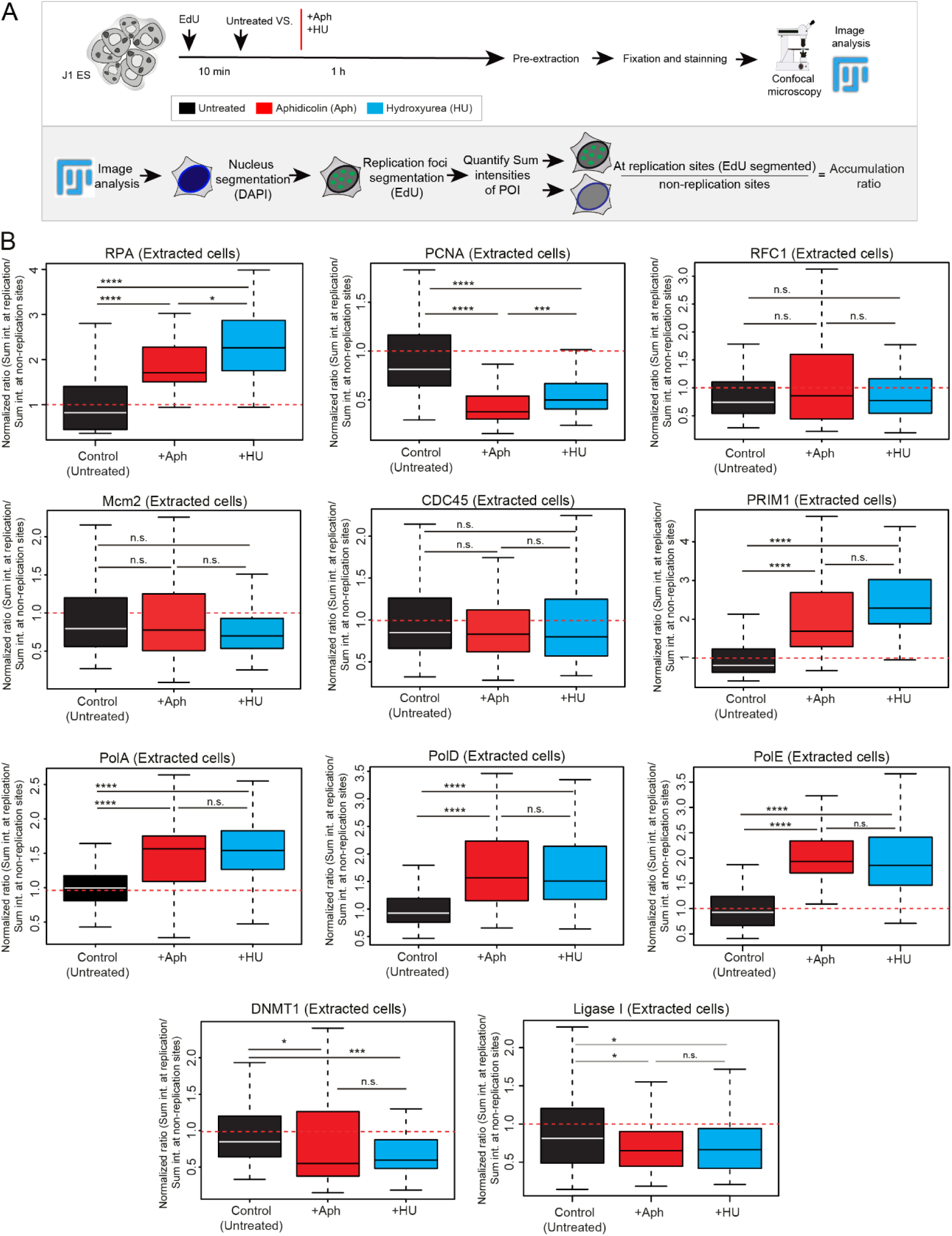
Fixed cell immunofluorescence analysis of the abundance of chromatin-bound replisome factors at replication sites after 1 hour of aphidicolin or hydroxyurea treatment relative to an untreated control. Cells were pre-extracted before fixation to remove the soluble protein pool from the nuclei. (A) Experimental and image analysis pipeline for obtaining the accumulation ratio at replication sites. The sum signal intensity within replication foci (segmented based on EdU signal) was divided by the sum signal intensity within the nucleus (segmented based on DAPI signal), in line with the live cell data shown in Figure 1. All values were normalized by the average ratio of untreated samples. Dashed red lines show the average ratio in the respective control sample. (B) Box plots depicting the chromatin-bound abundance of the indicated replisome factors. 2-4 replicates were performed for each replisome factor, including 45-90 cells in total. Each box plot represents 50% of the data, starting in the first quartile (25%) and ending in the third (75%). The line inside represents the median. The whiskers represent the upper and lower quartiles. Statistical significance was tested with a paired two-samples Wilcoxon test (n.s. is given for p-values ≥ 0.05; 1 star (*) for p-values < 0.05 and ≥ 0.005; 2 stars (**) are given for values < 0.005 and ≥ 0.0005; 3 stars (***) are given for values < 0.0005 and ≥ 0.00005; 4 stars (****) are given for values < 0.00005).

**Figure S4.**
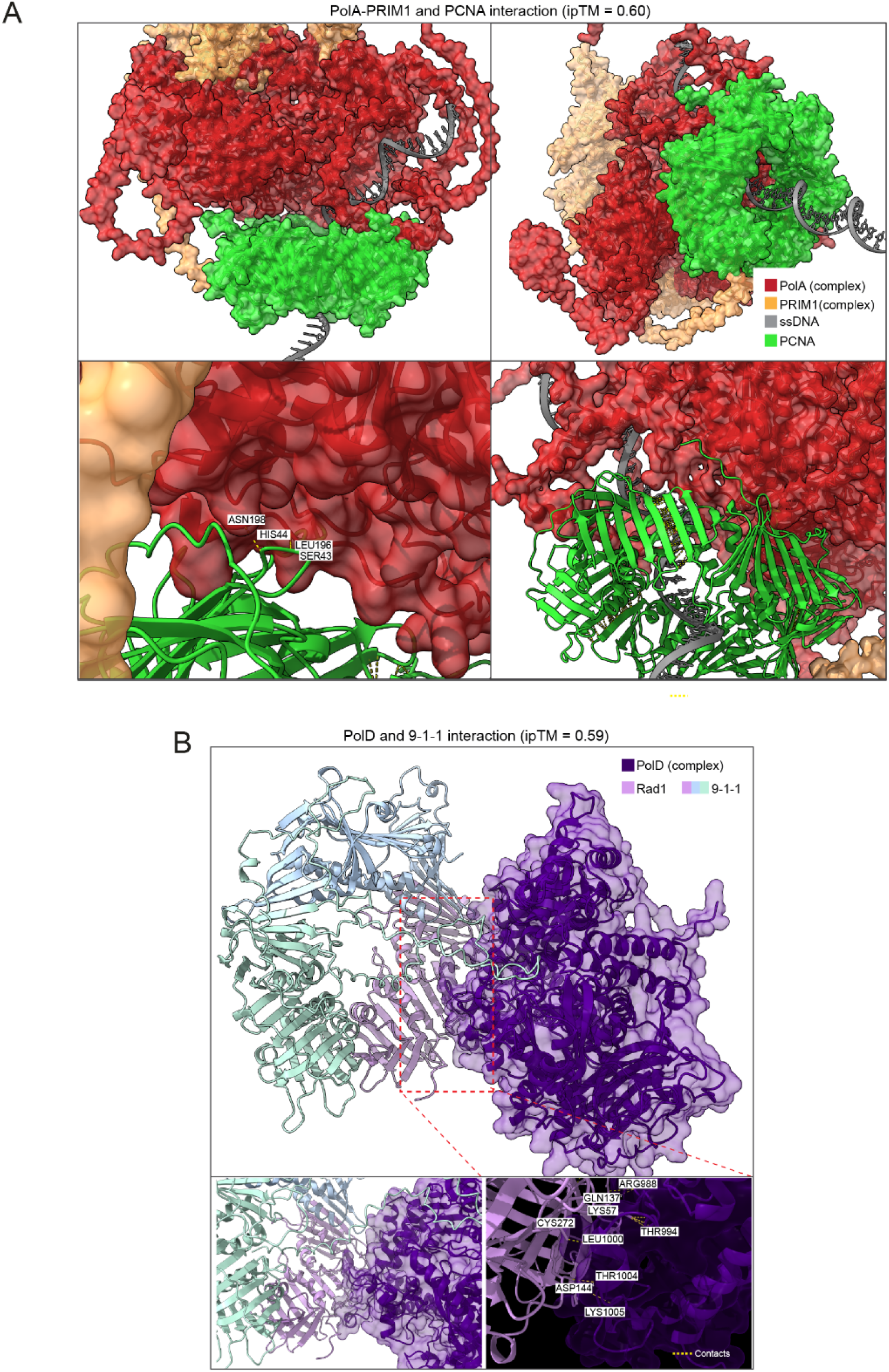
Structural models of the predicted PolA-PRIM1/PCNA and PolD/9-1-1 complex interactions. (A) 3D surface model of the PolA-PRIM1/PCNA/ssDNA complex generated by AlphaFold 3. (B) 3D surface model of the PolD/9-1-1/ssDNA complex generated by AlphaFold 3. For (A) and (B), predicted contact residues at the protein-protein interfaces and ipTMs are indicated.

**Figure S5.**
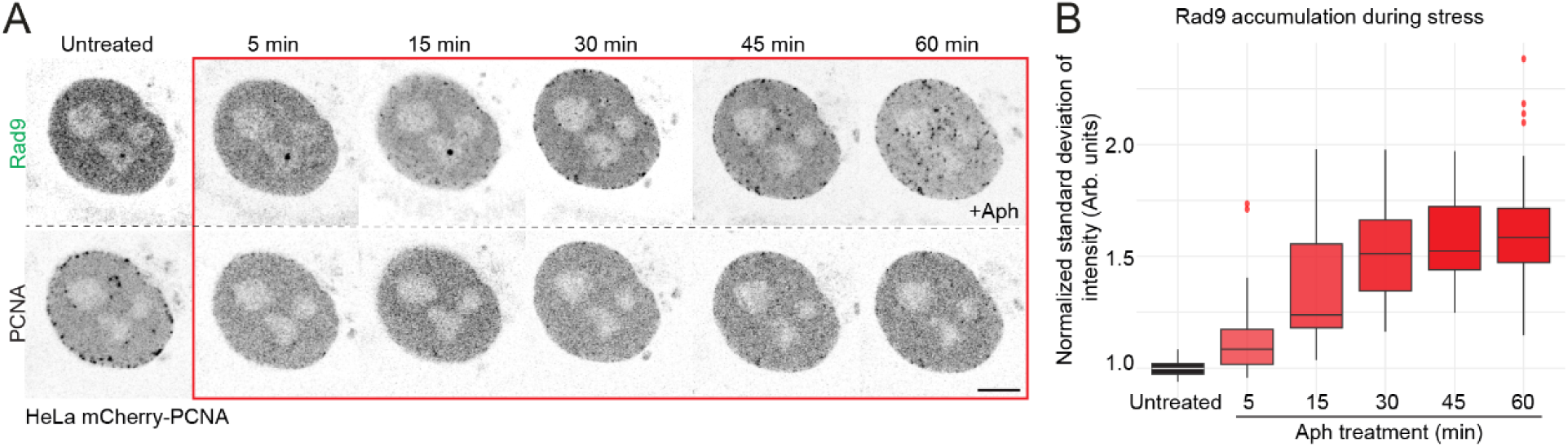
Live-cell imaging of GFP-tagged Rad9 dynamics during replication stress in HeLa cells. (A) Representative live-cell time-lapse images of HeLa mCherry-PCNA cells transiently expressing GFP-Rad9, captured before and during 1 hour treatment with 50 µM aphidicolin. Scale bar: 5 µm. (B) Quantification of GFP-Rad9 standard deviation at each time point normalized by the average standard deviation of untreated cells. Two replicates were performed encompassing a total of 15 cells. 7 ROIs were randomly selected for each nuclei, and their change in brightness was quantified over time.

**Figure S6.**
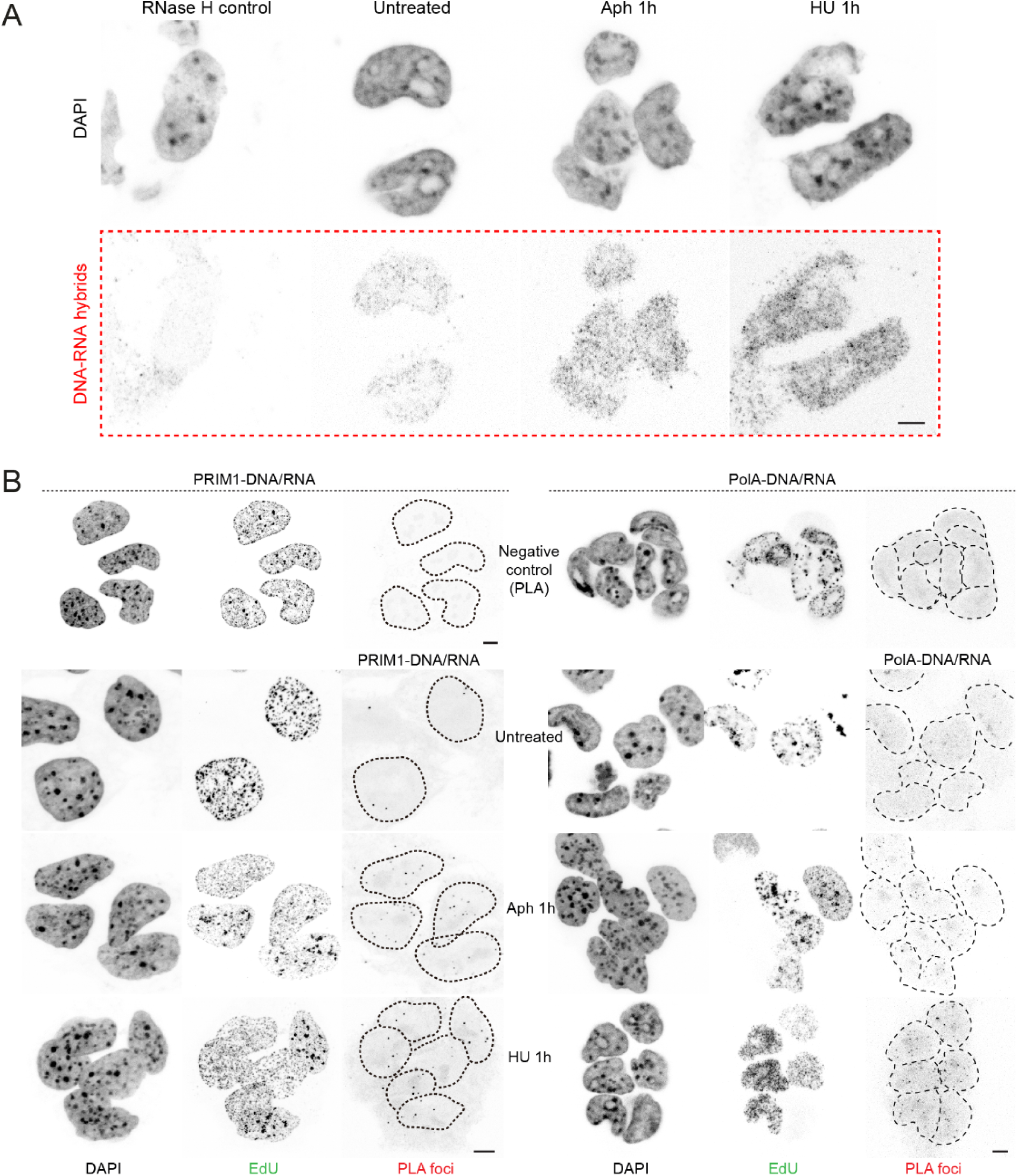
Example images showing the DNA/RNA hybrid formation and interaction with the PolA/Prim1 complex under untreated and stress conditions. (A) Representative immunofluorescence staining against DNA/RNA hybrids in pre-extracted cells. The negative control was performed by treatment with RNase H, which specifically digests DNA/RNA hybrids. Pre-extraction removes soluble DNA/RNA hybrids derived from transcription. (B) Representative images of proximity ligation assays shown in Figure 3B-C (DNA/RNA hybrids colocalizing with PRIM1 or PolA), including incubation with only the anti-DNA/RNA hybrid primary antibody as negative control. EdU was added before replication stress treatments and was used to select S-phase cells. Scale bar: 5 µm.

**Figure S7.**
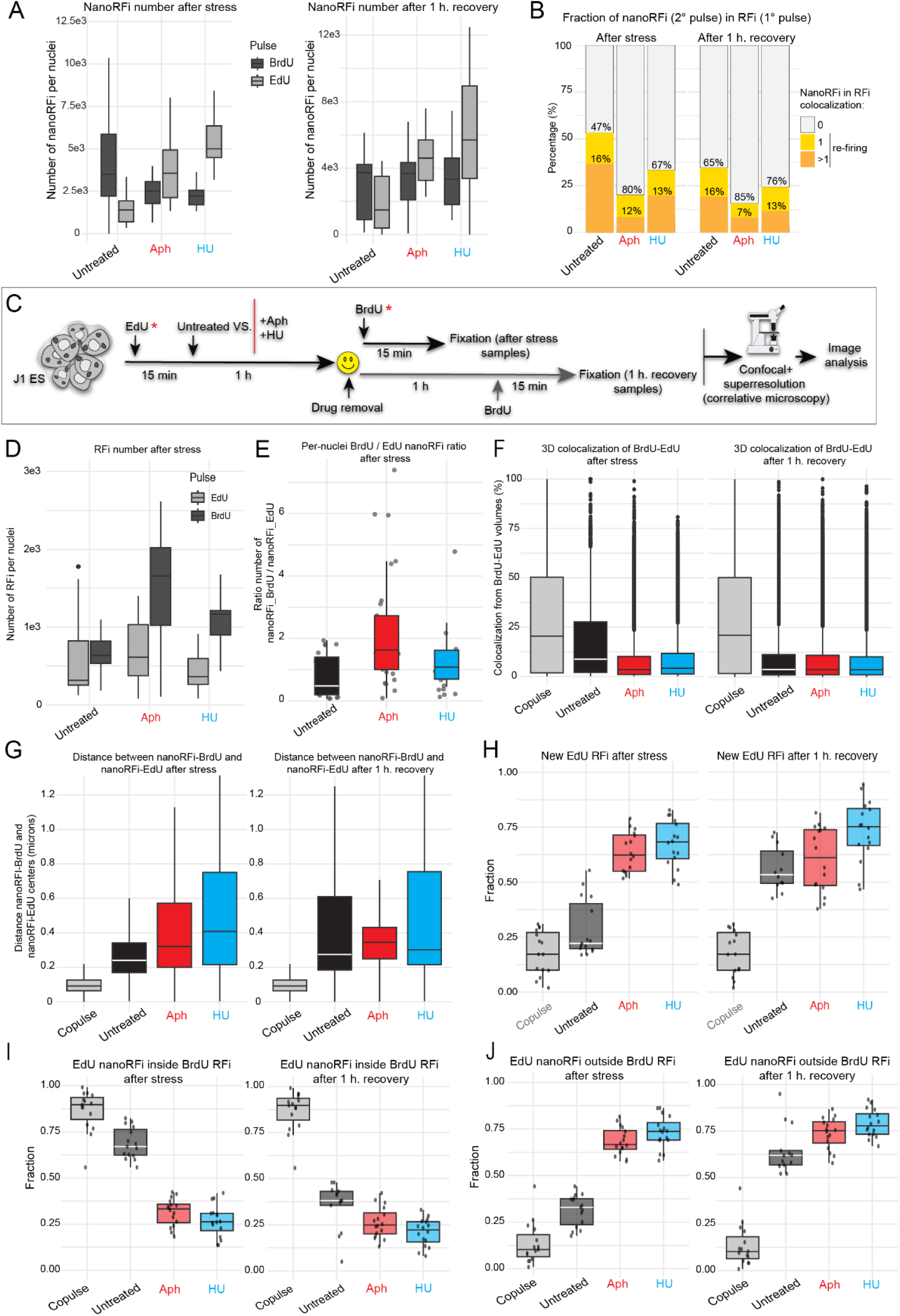
Colocalization analysis of pre- and post-stress replication foci. (A) Number of replication nano-foci (nanoRFi) per nucleus before (BrdU, dark gray) and after (EdU, light gray) stress induction. Left: EdU pulse immediately after stress removal; right: EdU pulse 1 hour after stress removal. (B) Adjusted treatment and imaging pipeline with inverted pulse order (EdU before stress, BrdU after stress). (C) Number of RFi per nucleus before (EdU, light gray) and after (BrdU, dark gray) stress induction using the inverted pulse order. The second pulse occurred immediately (left) or 1 hour (right) after stress removal. (D) Per-nucleus ratio of second-pulse nanoRFi per first-pulse nanoRFi using the inverted pulse order. (E) Number of EdU and BrdU nanoRFi per nucleus with inverted pulse, immediately and 1 hour after stress removal. (F) Percentage of three-dimensional overlap between replication foci before and after stress induction, quantified using the DiAna plugin (Fiji). Co-pulse samples (BrdU and EdU administered simultaneously) are included as a positive control for maximum overlap. Left: second pulse immediately after stress removal; right: second pulse 1 hour after stress removal. (G) Center-to-center distances between paired BrdU and EdU 3D foci classified as “touching” (sharing at least 1 voxel). (H) Fraction of EdU foci not touching any BrdU foci. (I-J) Complementary analysis showing the percentage of EdU nanoRFi falling within BrdU RFi. For all experiments, 3 independent replicates were performed, including 79-95 total cells per condition.

**Figure S8.**
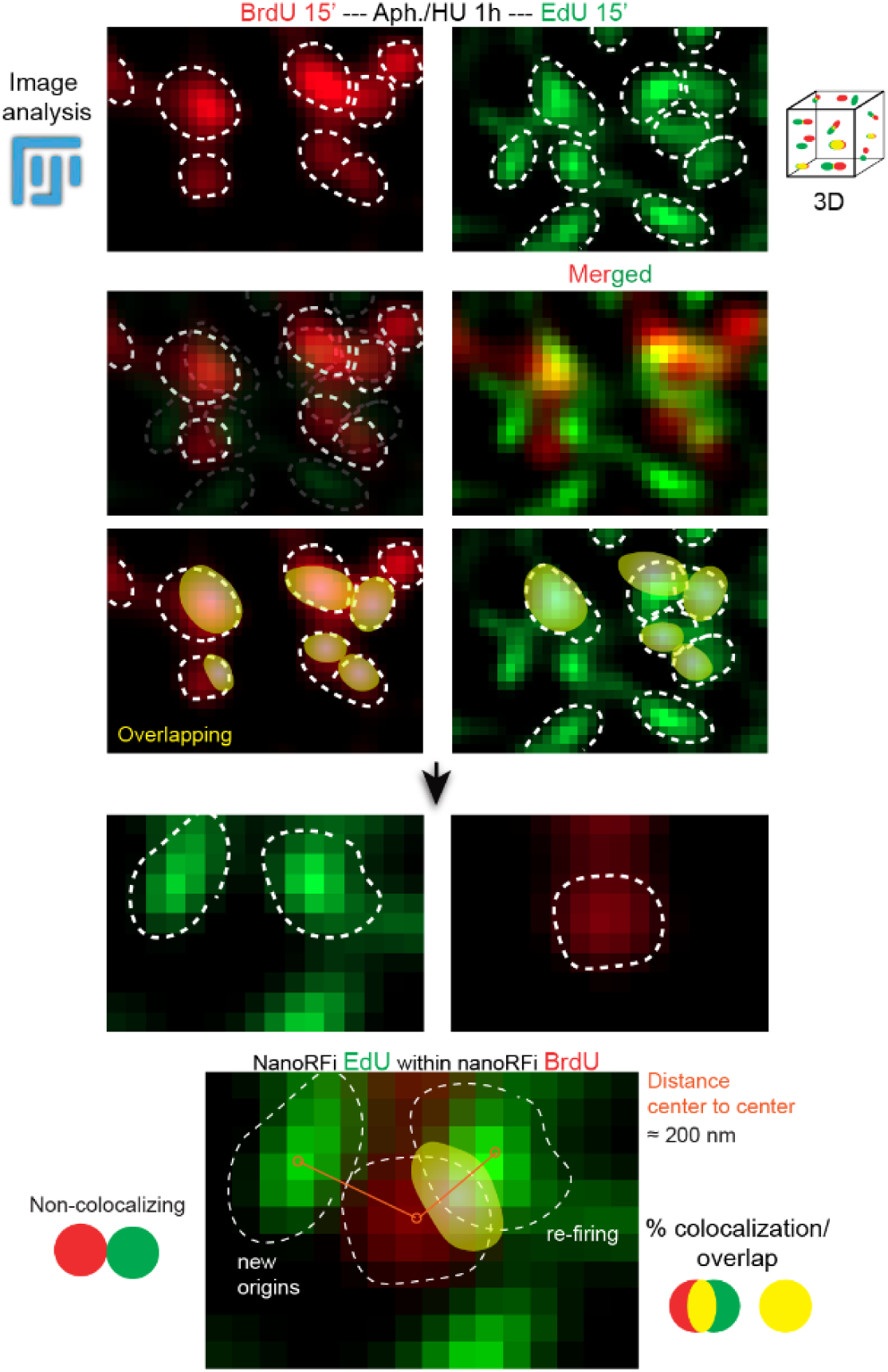
Representative images and segmentation pipeline for BrdU/EdU nanoRFi quantification. Superresolution images of nanoRFi labelled with BrdU before stress (red) and EdU after stress (green). The analysis pipeline utilized to determine the degree of overlap between BrdU and EdU foci is shown in a stepwise manner.

**Figure S9.**
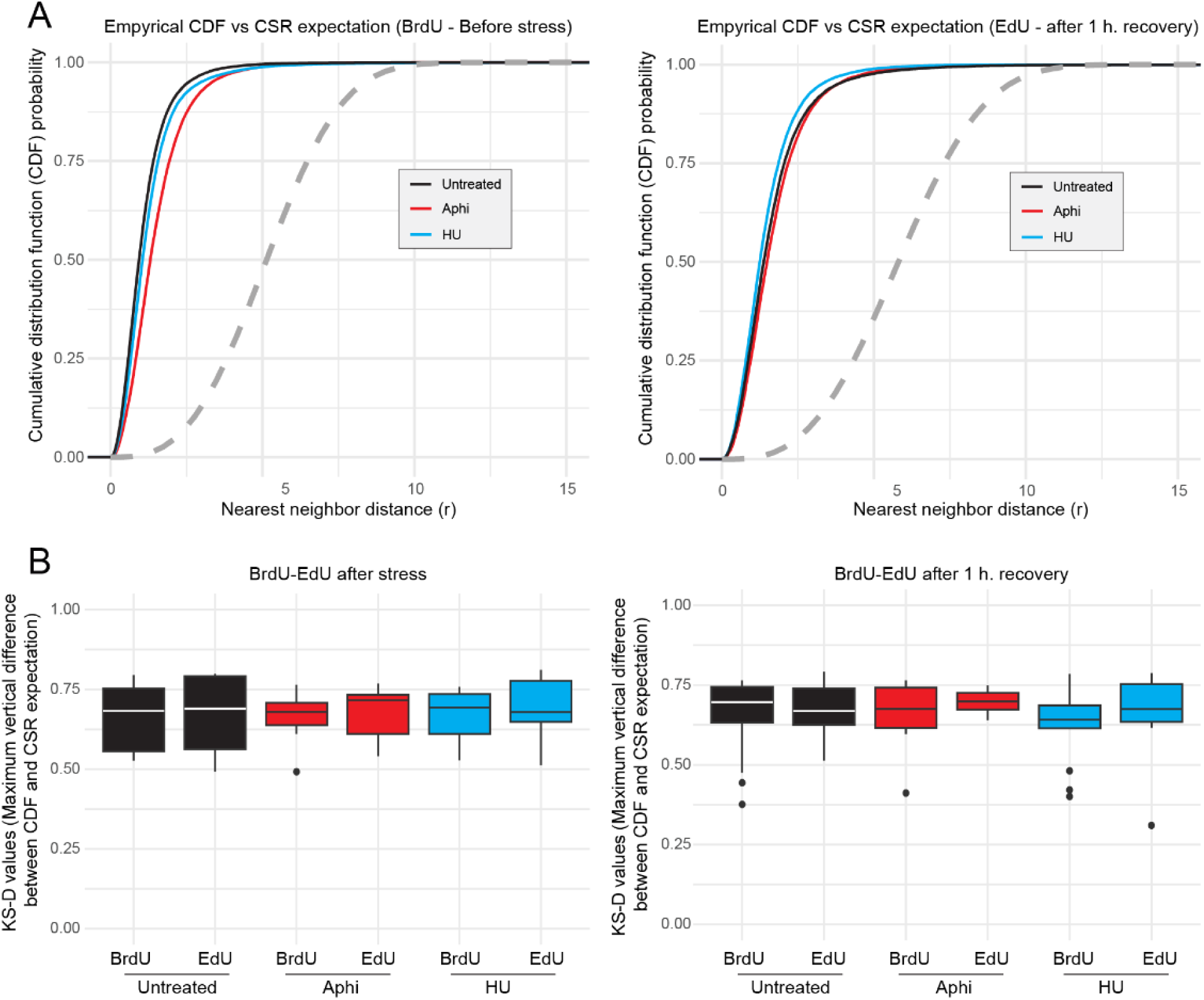
Spatial randomness analysis of nanoRFi distribution before and after stress. Empirical cumulative distribution functions (ECDFs, solid lines) of nearest-neighbor distances between segmented nanoRFi within individual nuclei were compared to the theoretical CDF expected under complete spatial randomness (CSR, dashed lines) in 3 dimensions. Untreated samples (black), aphidicolin-treated (red), and hydroxyurea-treated (blue) cells are compared. (A) Foci distance distributions in samples where the post-stress pulse was performed 1 hour after stress removal. Left: BrdU foci (pre-stress); right: EdU foci (post-stress). (B) Maximum vertical distance (KS-D statistic) between empirical and theoretical CDFs. For all experiments, 3 independent replicates were performed, including 79-95 cells per condition in total.

**Figure S10.**
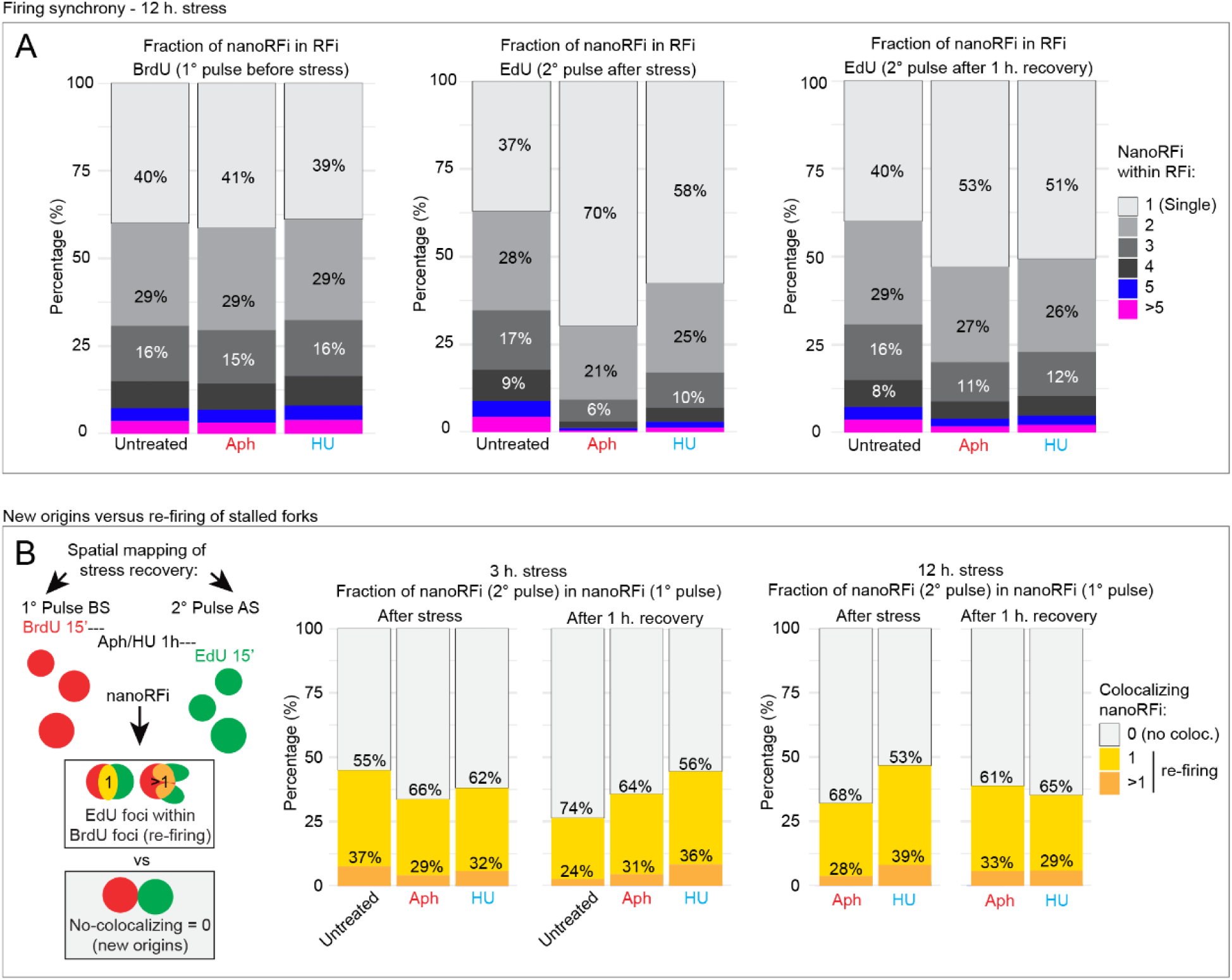
Replication synchrony and proportion of newly fired origins under conditions of intra-S checkpoint activation. (A) Distribution of the number of nanoRFi contained within individual RFi (replication synchrony) before stress and immediately or 1 hour after the end of 12-hour aphidicolin or hydroxyurea treatment. (B) Percentage of new and re-firing origins in the second pulse as determined by overlap with first pulse nanoRFi, quantified after 3-hour and 12-hour stress treatments. Across both treatment durations, 56-66% of post-stress replicons correspond to newly fired origins rather than re-firing stalled forks. Two independent replicates were performed, totaling 74-86 cells per condition.

### Supplementary tables

**Supplementary table 1:**
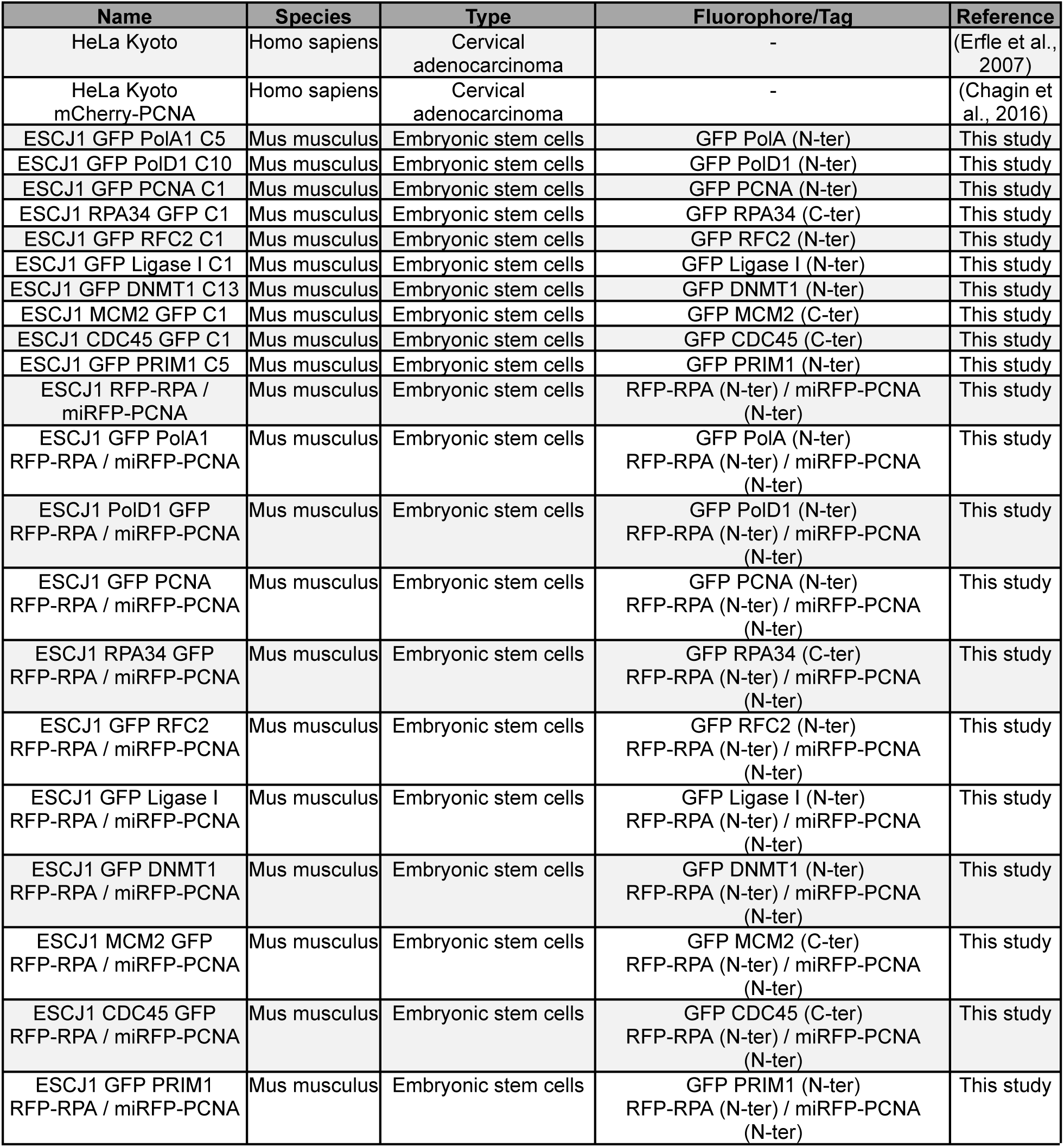
Cell line characteristics.

**Supplementary table 2:**
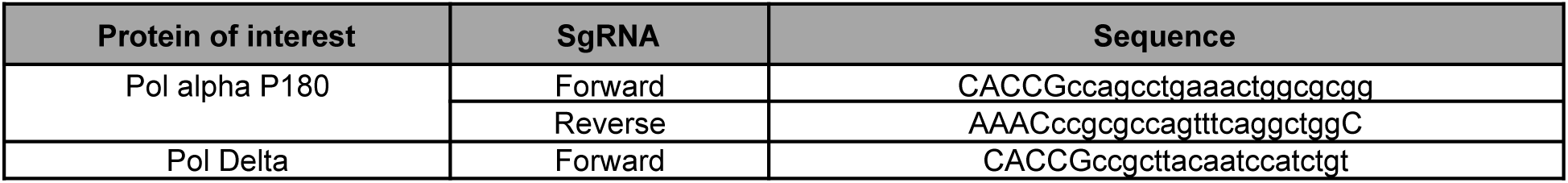
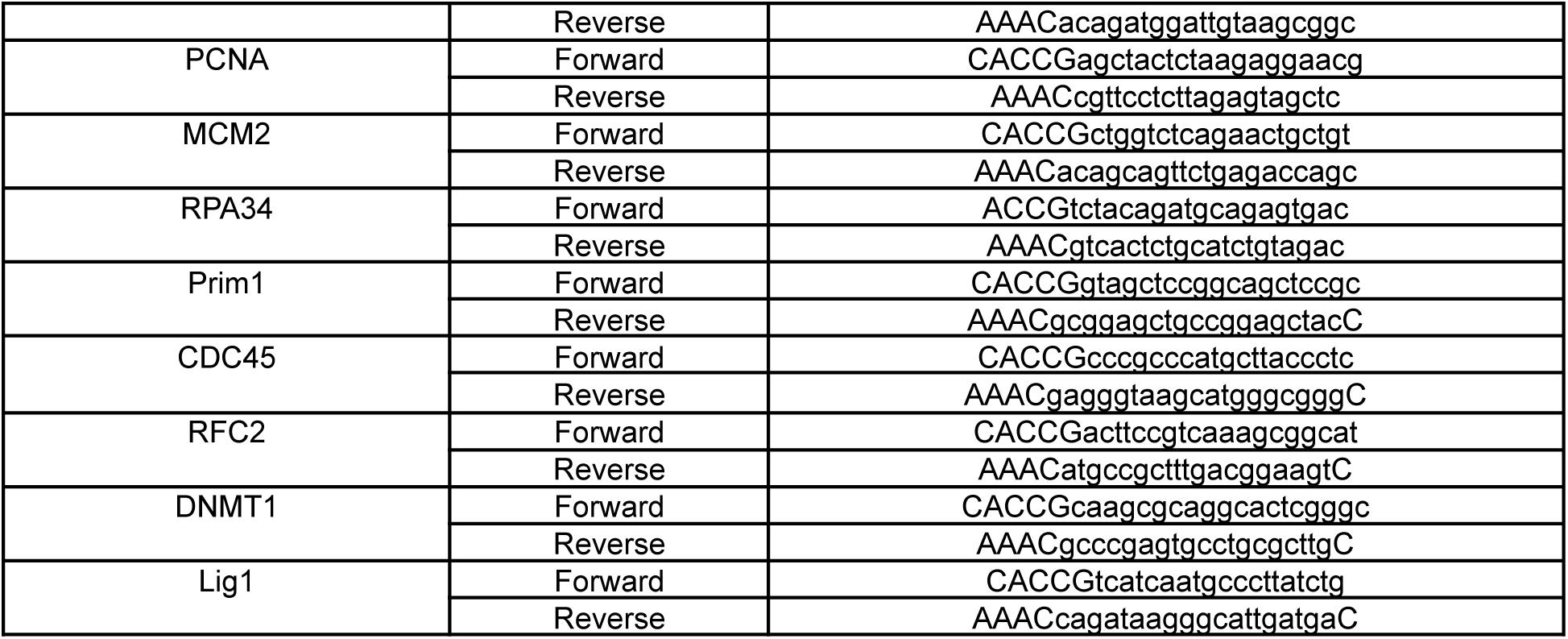
sgRNA sequences for CRISPR cell lines.

**Supplementary table 3:**
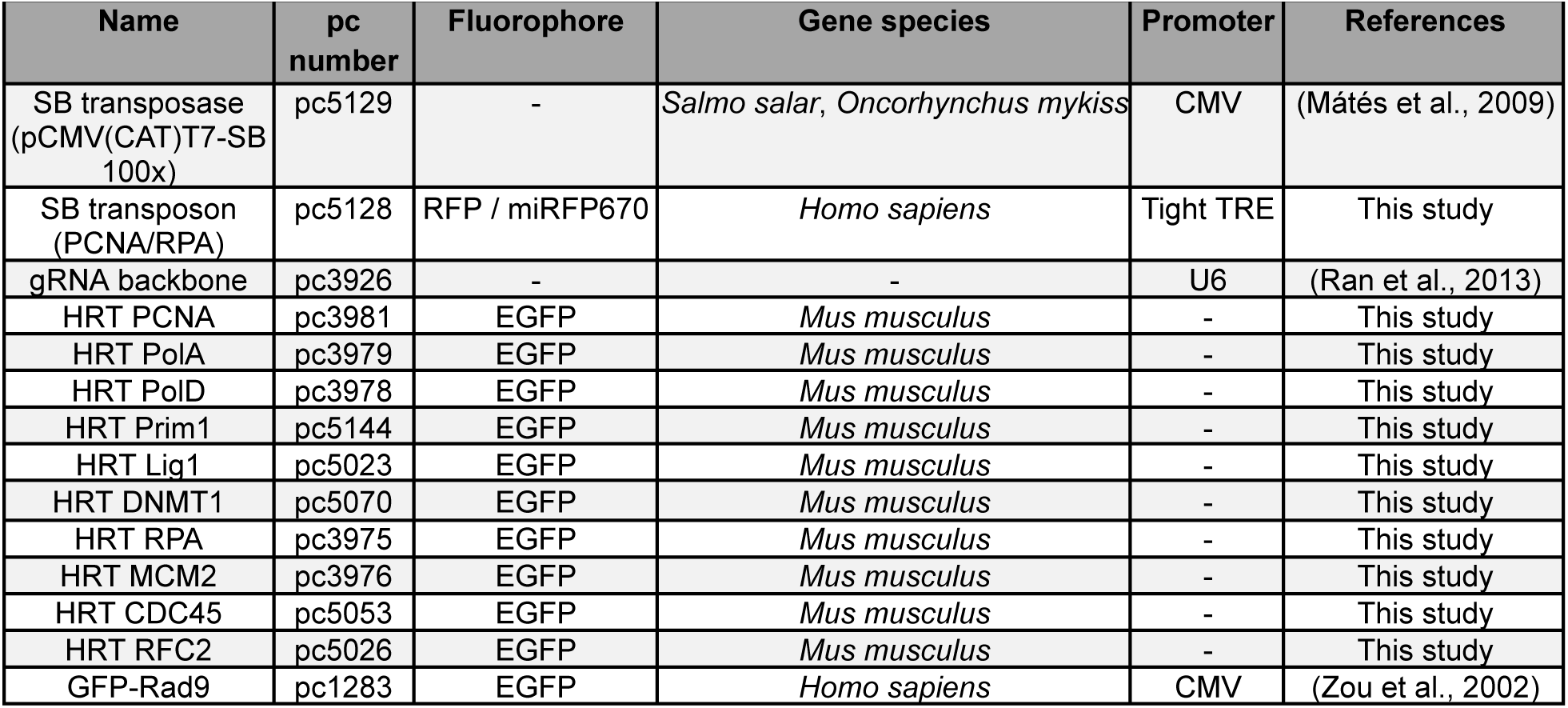
Plasmid characteristics.

**Supplementary table 4:** Nucleotide and inhibitor characteristics.

| Name | Application | Detection | Cat # | Company |
| --- | --- | --- | --- | --- |
| Hydroxyurea | Nucleotide synthesis blocking | - | H8627 | Sigma-Aldrich, St Louis, MO, USA |
| Aphidicolin | Polymerase inhibition | - | A0781-1MG | Sigma-Aldrich, St Louis, MO, USA |
| 5-ethynyl-2'-deoxyuridine (EdU) | Labeling of nascent DNA in pulse (chase) experiments | ClickIT chemistry | E10415 | Thermo Fisher Scientific, Waltham, MA, USA |
| 5-Bromo-2'-deoxyuridine (BrdU) | Labeling of nascent DNA in pulse (chase) experiments | Immunocytochemistry | B5002 | Sigma-Aldrich, St Louis, MO, USA |
| Roscovitine | CDK inhibition | - | 557360 | Merck Sigma-Aldrich Chemie |

**Supplementary table 5:** Primary and secondary antibody characteristics.

| Reactivity | Host | Clonality | Dilution | Application | Catalogue /clone no | Company |
| --- | --- | --- | --- | --- | --- | --- |
| anti-PCNA | Mouse | Monoclonal | 1:100 | IF* | M0879/PC 10 | Dako, Hamburg, Germany |
| anti-RPA70A serum | Mouse | Monoclonal | 1:2 | IF | 7G9E3 | Hybridoma (Kenny et al., 1990) |
| anti-RFC1 | Mouse | Monoclonal | 1:50 | IF | sc-271658 | Santa Cruz Biotechnology |
| anti-MCM2 | Rabbit | Monoclonal | 1:200 | IF | EPR4120 | Abcam, Cambridge, UK |
| anti-MCM2pS27 | Rabbit | Monoclonal | 1:200 | IF | EPR4119 | Epitomics, USA |
| anti-MCM2pS41 | Rabbit | Monoclonal | 1:100 | IF | 3252-1 | Epitomics, USA |
| anti-MCM2pS53 | Rabbit | Monoclonal | 1:200 | IF | EP4120 | Epitomics, USA |
| anti-MCM2pS108 | Rabbit | Monoclonal | 1:1000 | IF | 3267-1 | Epitomics, USA |
| anti-CDC45 | Rabbit | Monoclonal | 1:50 | IF | 556-1 | Epitomics, USA |
| PRIM1 | Rabbit | Polyclonal | 1:100 | IF/PLA | 10773-1-AP | Proteintech, Germany |
| anti-PolA | Mouse | Monoclonal | Undiluted | IF | SJK287-38 | Hybridoma (Tanaka et al., 1982) |
| anti-PolD | Mouse | Monoclonal | 1:500 | IF, WB | 610972 | BD Biosciences, Germany |
| anti-PolE | Rabbit | Polyclonal | 1:500 | IF, WB | GTX132100 | Gene Tex, Germany |
| anti-Rad1 | Rabbit | Polyclonal | 1:50 | IF/PLA | 11726-2-AP | Proteintech, Germany |
| anti-DNMT1 | Rat | Monoclonal | 1:10 | IF | 2E8(AA171-256) | Ludwig-Maximilians-Universität München |
| anti-Ligasel | Mouse | Monoclonal | 1:100 | IF | 084K1062 | Sigma-Aldrich, St Louis, MO, USA |
| anti-BrdU | Rabbit | Polyclonal | 1:500 | IF | 600-401-C29 | Rockland Immunochemicals, USA |
| anti-mouse IgG Cy3 | Donkey | Polyclonal | 1:800 | IF (secondary) | 715-165-151 | The Jackson Laboratory, USA |
| anti-rabbit IgG Cy3 | Donkey | Polyclonal | 1:800 | IF (secondary) | 711-165-152 | The Jackson Laboratory, USA |
| anti-rabbit IgG (H+L) AF594 | Goat | Polyclonal | 1:500 | IF (secondary) | 111-585-144 | The Jackson Laboratory, USA |
| anti-rat IgG AF594 | Donkey | Polyclonal | 1:500 | IF (secondary) | 712-585-153 | The Jackson Laboratory, USA |
| anti-rat IgG AF488 | Donkey | Polyclonal | 1:800 | IF (secondary) | A11006 | Invitrogen, USA |
| anti-mouse IgG AF488 | Goat | Polyclonal | 1:800 | IF (secondary) | 2120125 | Invitrogen, USA |
| anti-rabbit IgG AF488 | Donkey | Polyclonal | 1:800 | IF (secondary) | A11034 | Invitrogen, USA |

\* requires methanol treatment

**Supplementary table 6:** Imaging systems characteristics.

| Microscope/<br>Company | Lasers/lamps | Filters (ex. & em.<br>[nm])* | Objectives/lenses used | Detection system |
| --- | --- | --- | --- | --- |
| Ultra-View VoX<br>spinning disk on<br>an inverted Nikon<br>Ti-E microscope,<br>PerkinElmer Life<br>Sciences, UK | Diode lasers<br>(405 nm,<br>488 nm,<br>561 nm,<br>640 nm) | *405: 415–475<br>*488: 505–549<br>*561: 580–650<br>*640: 664–754<br>405/488/568/640** | oil immersion<br>60x Plan-Apochromat<br>(NA 1.45) | cooled 14-bit<br>Hamamatsu®<br>C9100-50<br>EMCCD |
| Widefield<br>Axiovert<br>200, Zeiss,<br>Germany | HBO100<br>mercury<br>amp | 488: 473-491 & 506-534<br>561: 550-580 & 590-650<br>640: 590-650 & 663-738 | oil immersion<br>63x Plan-Apochromat<br>(1.4 NA) | 12-bit<br>AxioCam<br>mRM |
| Leica SP5 II,<br>Wetzlar, Germany | 405 nm diode 488 nm<br>Argon, 561 nm DPSS,<br>633 nm HeNe gas | AOBS beam splitter | HCX PL APO 63x / 1.4-0.6<br>oil lambda blue & HCX PL<br>APO 100x / 1.44 oil Corr CS | 2 HyD Hybrid Detectors |
| Zeiss LSM 900<br>Airyscan, Zeiss,<br>Germany | LEDs modules: 385<br>nm, 470nm, 540-580<br>nm, 625 nm<br>Laser Power:<br>405 nm- 5 mW 488<br>nm- 10 mW 561 nm- 10<br>mW 640 nm- 5 mW | ex: 390/40 em: 450/40<br>ex: 470/40 em: 525/50<br>ex: 550/25 em: 605/70 &<br>ex: 640/30 em: 690/50 | EC Plan-Neofluar 2.5x<br>Plan-Apochromat 20x<br>Plan-Apochromat 40x<br>C Plan-Apochromat 63x<br>alpha Plan-Apochromat 100x | Axiocam 820 mono SONY<br>IMX541 CMOS back<br>illuminated pixel size 2.74 µm x<br>2.74 µm 4512 x 4512 pixel |

\*ex.: excitation & em.: emission, \*\* dichroic specification, \*\*\* WD: working distance

**Supplementary table 7:**
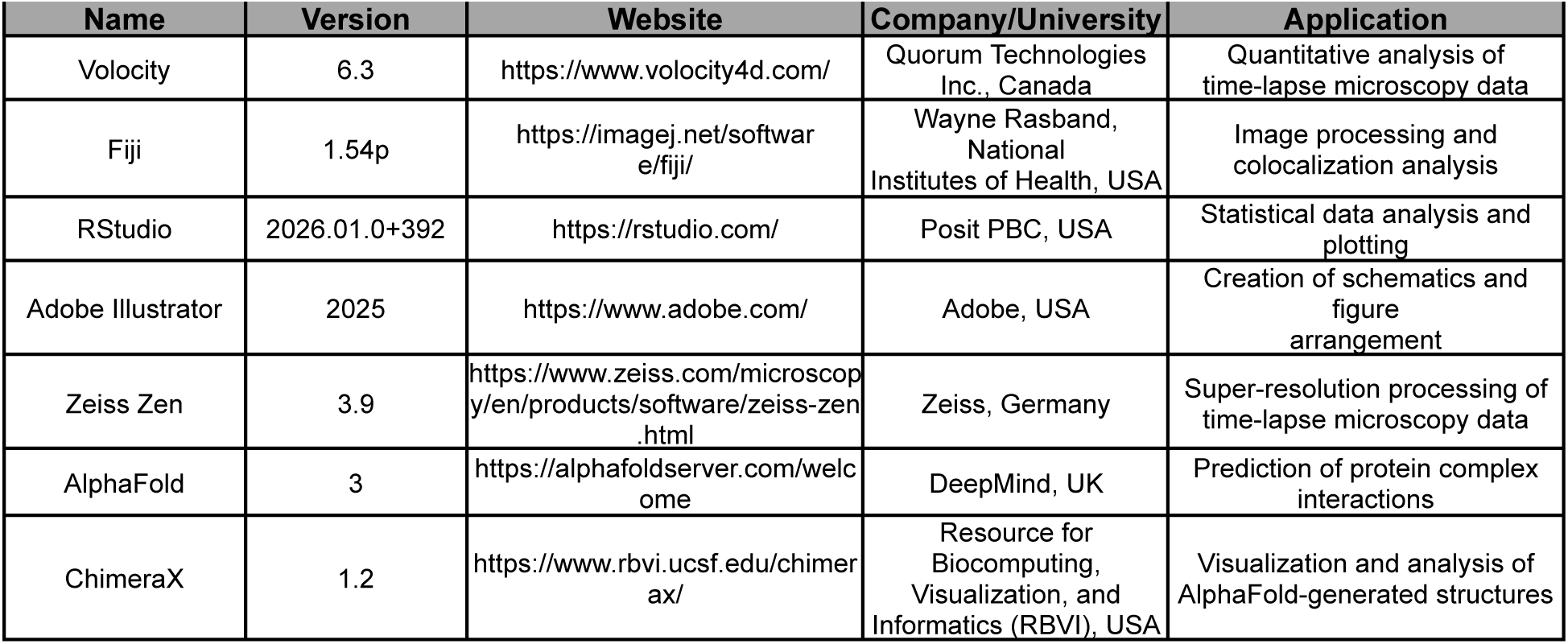
Software.

## REFERENCES

Abramson J, Adler J, Dunger J, Evans R, Green T, Pritzel A, Ronneberger O, Willmore L, Ballard AJ, Bambrick J, Bodenstein SW, Evans DA, Hung C-C, O’Neill M, Reiman D, Tunyasuvunakool K, Wu Z, Žemgulytė A, Arvaniti E, Beattie C, Jumper JM. 2024. Accurate structure prediction of biomolecular interactions with AlphaFold 3. Nature 630:493–500. doi:10.1038/s41586-024-07487-w

Acharya N, Prakash L, Prakash S. 2023. Yeast 9-1-1 complex acts as a sliding clamp for DNA synthesis by DNA polymerase ε. J Biol Chem 299:102727. doi:10.1016/j.jbc.2022.102727

Arroyo M, Cardoso CM, Hastert FD. 2023. In situ Quantification of Cytosine Modification Levels in Heterochromatic Domains of Cultured Mammalian Cells. Bio Protoc 13:e4716. doi:10.21769/BioProtoc.4716

Arroyo M, Casas-Delucchi CS, Pabba MK, Prorok P, Pradhan SK, Rausch C, Lehmkuhl A, Maiser A, Buschbeck M, Pasque V, Bernstein E, Luck K, Cardoso MC. 2024. Histone variant macroH2A1 regulates synchronous firing of replication origins in the inactive X chromosome. Nucleic Acids Res 52:11659–11688. doi:10.1093/nar/gkae734

Baranovskiy AG, Babayeva ND, Suwa Y, Gu J, Pavlov YI, Tahirov TH. 2014. Structural basis for inhibition of DNA replication by aphidicolin. Nucleic Acids Res 42:14013–14021. doi:10.1093/nar/gku1209

Bellelli R, Boulton SJ. 2021. Spotlight on the Replisome: Aetiology of DNA Replication-Associated Genetic Diseases. Trends Genet 37:317–336. doi:10.1016/j.tig.2020.09.008

Berezney R, Dubey DD, Huberman JA. 2000. Heterogeneity of eukaryotic replicons, replicon clusters, and replication foci. Chromosoma 108:471–484. doi:10.1007/s004120050399

Blow JJ, Dutta A. 2005. Preventing re-replication of chromosomal DNA. Nat Rev Mol Cell Biol 6:476–486. doi:10.1038/nrm1663

Blow JJ, Ge XQ, Jackson DA. 2011. How dormant origins promote complete genome replication. Trends Biochem Sci 36:405–414. doi:10.1016/j.tibs.2011.05.002

Blow JJ, Hodgson B. 2002. Replication licensing--defining the proliferative state? Trends Cell Biol 12:72–78. doi:10.1016/s0962-8924(01)02203-6

Bryant P, Pozzati G, Elofsson A. 2022. Improved prediction of protein-protein interactions using AlphaFold2. Nat Commun 13:1265. doi:10.1038/s41467-022-28865-w

Chagin VO, Casas-Delucchi CS, Reinhart M, Schermelleh L, Markaki Y, Maiser A, Bolius JJ, Bensimon A, Fillies M, Domaing P, Rozanov YM, Leonhardt H, Cardoso MC. 2016. 4D Visualization of replication foci in mammalian cells corresponding to individual replicons. Nat Commun 7:11231. doi:10.1038/ncomms11231

Cortez D, Glick G, Elledge SJ. 2004. Minichromosome maintenance proteins are direct targets of the ATM and ATR checkpoint kinases. Proc Natl Acad Sci USA 101:10078–10083. doi:10.1073/pnas.0403410101

Costa A, Diffley JFX. 2022. The initiation of eukaryotic DNA replication. Annu Rev Biochem 91:107–131. doi:10.1146/annurev-biochem-072321-110228

Day M, Oliver AW, Pearl LH. 2022. Structure of the human RAD17-RFC clamp loader and 9-1-1 checkpoint clamp bound to a dsDNA-ssDNA junction. Nucleic Acids Res 50:8279–8289. doi:10.1093/nar/gkac588

Derangère V, Bruchard M, Végran F, Ghiringhelli F. 2016. Proximity ligation assay (PLA) protocol using duolink® for T cells. Bio Protoc 6. doi:10.21769/BioProtoc.1811

Devbhandari S, Remus D. 2020. Rad53 limits CMG helicase uncoupling from DNA synthesis at replication forks. Nat Struct Mol Biol 27:461–471. doi:10.1038/s41594-020-0407-7

De March M, Merino N, Barrera-Vilarmau S, Crehuet R, Onesti S, Blanco FJ, De Biasio A. 2017. Structural basis of human PCNA sliding on DNA. Nat Commun 8:13935. doi:10.1038/ncomms13935

Diffley JFX. 2011. Quality control in the initiation of eukaryotic DNA replication. Philos Trans R Soc Lond B Biol Sci 366:3545–3553. doi:10.1098/rstb.2011.0073

Erfle H, Neumann B, Liebel U, Rogers P, Held M, Walter T, Ellenberg J, Pepperkok R. 2007. Reverse transfection on cell arrays for high content screening microscopy. Nat Protoc 2:392–399. doi:10.1038/nprot.2006.483

Evans R, O’Neill M, Pritzel A, Antropova N, Senior AW, Green T, Žídek A, Bates R, Blackwell S, Yim J, Ronneberger O, Bodenstein S, Zielinski M, Bridgland A, Potapenko A, Cowie A, Tunyasuvunakool K, Jain R, Clancy E, Kohli P, Hassabis D. 2021. Protein complex prediction with AlphaFold-Multimer. BioRxiv. doi:10.1101/2021.10.04.463034

Frisbie V, Bleichert F. 2026. License to replicate: mechanisms of licensing eukaryotic origins for DNA replication. Bioessays 48:e70095. doi:10.1002/bies.70095

Ganier O, Prorok P, Akerman I, Méchali M. 2019. Metazoan DNA replication origins. Curr Opin Cell Biol 58:134–141. doi:10.1016/j.ceb.2019.03.003

Garcia-Diaz M, Bebenek K. 2007. Multiple functions of DNA polymerases. CRC Crit Rev Plant Sci 26:105–122. doi:10.1080/07352680701252817

Gaudet F, Talbot D, Leonhardt H, Jaenisch R. 1998. A short DNA methyltransferase isoform restores methylation in vivo. J Biol Chem 273:32725–32729. doi:10.1074/jbc.273.49.32725

Gilles J-F, Dos Santos M, Boudier T, Bolte S, Heck N. 2017. DiAna, an ImageJ tool for object-based 3D co-localization and distance analysis. Methods 115:55–64. doi:10.1016/j.ymeth.2016.11.016

Gonzalez MA, Tachibana KK, Laskey RA, Coleman N. 2005. Control of DNA replication and its potential clinical exploitation. Nat Rev Cancer 5:135–141. doi:10.1038/nrc1548

Görisch SM, Sporbert A, Stear JH, Grunewald I, Nowak D, Warbrick E, Leonhardt H, Cardoso MC. 2008. Uncoupling the replication machinery: replication fork progression in the absence of processive DNA synthesis. Cell Cycle 7:1983–1990. doi:10.4161/cc.7.13.6094

Hanaoka F, Kato H, Ikegami S, Oashi M, Yamada M. 1979. Aphidicolin does inhibit repair replication in HeLa cells. Biochem Biophys Res Commun 87:575–580. doi:10.1016/0006-291x(79)91833-3

Harrington C, Perrino FW. 1995. Initiation of RNA-primed DNA synthesis in vitro by DNA polymerase alpha-primase. Nucleic Acids Res 23:1003–1009. doi:10.1093/nar/23.6.1003

Hastert FD, Weber J, Bauer C, Zhadan A, Singh DND, Dix TC, Arnold R, Bessonov S, Soller M, Leonhardt H, Cardoso MC, Arroyo M. 2025. TET dioxygenases localize at splicing speckles and promote RNA splicing. Nucleus 16:2536902. doi:10.1080/19491034.2025.2536902

Hozák P, Jackson DA, Cook PR. 1994. Replication factories and nuclear bodies: the ultrastructural characterization of replication sites during the cell cycle. J Cell Sci 107 ( **Pt 8**):2191–2202. doi:10.1242/jcs.107.8.2191

Johansson E, Macneill SA. 2010. The eukaryotic replicative DNA polymerases take shape. Trends Biochem Sci 35:339–347. doi:10.1016/j.tibs.2010.01.004

Jones ML, Aria V, Baris Y, Yeeles JTP. 2023. How Pol α-primase is targeted to replisomes to prime eukaryotic DNA replication. Mol Cell 83:2911–2924.e16. doi:10.1016/j.molcel.2023.06.035

Kenny MK, Schlegel U, Furneaux H, Hurwitz J. 1990. The role of human single-stranded DNA binding protein and its individual subunits in simian virus 40 DNA replication. J Biol Chem 265:7693–7700. doi:10.1016/S0021-9258(19)39170-7

Koç A, Wheeler LJ, Mathews CK, Merrill GF. 2004. Hydroxyurea arrests DNA replication by a mechanism that preserves basal dNTP pools. J Biol Chem 279:223–230. doi:10.1074/jbc.M303952200

Kose HB, Xie S, Cameron G, Strycharska MS, Yardimci H. 2019. Mechanism of RPA-Facilitated Processive DNA Unwinding by the Eukaryotic CMG Helicase. BioRxiv. doi:10.1101/796003

Koundrioukoff S, Carignon S, Técher H, Letessier A, Brison O, Debatisse M. 2013. Stepwise activation of the ATR signaling pathway upon increasing replication stress impacts fragile site integrity. PLoS Genet 9:e1003643. doi:10.1371/journal.pgen.1003643

Lee K-Y, Park SH. 2020. Eukaryotic clamp loaders and unloaders in the maintenance of genome stability. Exp Mol Med 52:1948–1958. doi:10.1038/s12276-020-00533-3

Lin M, Martin J, Baxter R. 2015. Proximity Ligation Assay (PLA) to Detect Protein-protein Interactions in Breast Cancer. Bio Protoc 5. doi:10.21769/BioProtoc.1479

Li E, Bestor TH, Jaenisch R. 1992. Targeted mutation of the DNA methyltransferase gene results in embryonic lethality. Cell 69:915–926. doi:10.1016/0092-8674(92)90611-f

Li N, Gao N, Zhai Y. 2023. DDK promotes DNA replication initiation: Mechanistic and structural insights. Curr Opin Struct Biol 78:102504. doi:10.1016/j.sbi.2022.102504

Löb D, Lengert N, Chagin VO, Reinhart M, Casas-Delucchi CS, Cardoso MC, Drossel B. 2016. 3D replicon distributions arise from stochastic initiation and domino-like DNA replication progression. Nat Commun 7:11207. doi:10.1038/ncomms11207

Mašata M, Juda P, Raška O, Cardoso MC, Raška I. 2011. A fraction of MCM 2 proteins remain associated with replication foci during a major part of S phase. Folia Biol (Praha) 57:3–11.

Mátés L, Chuah MKL, Belay E, Jerchow B, Manoj N, Acosta-Sanchez A, Grzela DP, Schmitt A, Becker K, Matrai J, Ma L, Samara-Kuko E, Gysemans C, Pryputniewicz D, Miskey C, Fletcher B, VandenDriessche T, Ivics Z, Izsvák Z. 2009. Molecular evolution of a novel hyperactive Sleeping Beauty transposase enables robust stable gene transfer in vertebrates. Nat Genet 41:753–761. doi:10.1038/ng.343

Maya-Mendoza A, Olivares-Chauvet P, Shaw A, Jackson DA. 2010. S phase progression in human cells is dictated by the genetic continuity of DNA foci. PLoS Genet 6:e1000900. doi:10.1371/journal.pgen.1000900

McGann PT, Ware RE. 2015. Hydroxyurea therapy for sickle cell anemia. Expert Opin Drug Saf 14:1749–1758. doi:10.1517/14740338.2015.1088827

Méchali M. 2001. DNA replication origins: from sequence specificity to epigenetics. Nat Rev Genet 2:640–645. doi:10.1038/35084598

Montagnoli A, Valsasina B, Brotherton D, Troiani S, Rainoldi S, Tenca P, Molinari A, Santocanale C. 2006. Identification of Mcm2 phosphorylation sites by S-phase-regulating kinases. J Biol Chem 281:10281–10290. doi:10.1074/jbc.M512921200

Mortusewicz O, Rothbauer U, Cardoso MC, Leonhardt H. 2006. Differential recruitment of DNA Ligase I and III to DNA repair sites. Nucleic Acids Res 34:3523–3532. doi:10.1093/nar/gkl492

Moyer SE, Lewis PW, Botchan MR. 2006. Isolation of the Cdc45/Mcm2-7/GINS (CMG) complex, a candidate for the eukaryotic DNA replication fork helicase. Proc Natl Acad Sci USA 103:10236–10241. doi:10.1073/pnas.0602400103

O’Donnell M, Li H. 2016. The eukaryotic replisome goes under the microscope. Curr Biol 26:R247–56. doi:10.1016/j.cub.2016.02.034

Pabba MK, Ritter C, Chagin VO, Meyer J, Celikay K, Stear JH, Loerke D, Kolobynina K, Prorok P, Schmid AK, Leonhardt H, Rohr K, Cardoso MC. 2023. Replisome loading reduces chromatin motion independent of DNA synthesis. eLife 12. doi:10.7554/eLife.87572

Pettersen EF, Goddard TD, Huang CC, Meng EC, Couch GS, Croll TI, Morris JH, Ferrin TE. 2021. UCSF ChimeraX: structure visualization for researchers, educators, and developers. Protein Sci 30:70–82. doi:10.1002/pro.3943

Pradhan SK, Lozoya T, Prorok P, Yuan Y, Lehmkuhl A, Zhang P, Cardoso MC. 2024. Developmental Changes in Genome Replication Progression in Pluripotent versus Differentiated Human Cells. Genes 15. doi:10.3390/genes15030305

Pradhan SK, Zhang H, Kolobynina KG, Rapp A, Arroyo M, Cardoso MC. 2025. Dynamic association of H3K36me3 with pericentromeric heterochromatin regulates its replication time. EMBO Rep 26:4950–4976. doi:10.1038/s44319-025-00575-6

Prorok P, Wolf E, Cardoso MC. 2024. Timeless-Tipin interactions with MCM and RPA mediate DNA replication stress response. Frontiers in Cell and Developmental Biology.

Pursell ZF, Isoz I, Lundström E-B, Johansson E, Kunkel TA. 2007. Yeast DNA polymerase epsilon participates in leading-strand DNA replication. Science 317:127–130. doi:10.1126/science.1144067

Qin W, Wolf P, Liu N, Link S, Smets M, La Mastra F, Forné I, Pichler G, Hörl D, Fellinger K, Spada F, Bonapace IM, Imhof A, Harz H, Leonhardt H. 2015. DNA methylation requires a DNMT1 ubiquitin interacting motif (UIM) and histone ubiquitination. Cell Res 25:911–929. doi:10.1038/cr.2015.72

Ran FA, Hsu PD, Wright J, Agarwala V, Scott DA, Zhang F. 2013. Genome engineering using the CRISPR-Cas9 system. Nat Protoc 8:2281–2308. doi:10.1038/nprot.2013.143

Rausch C, Weber P, Prorok P, Hörl D, Maiser A, Lehmkuhl A, Chagin VO, Casas-Delucchi CS, Leonhardt H, Cardoso MC. 2020. Developmental differences in genome replication program and origin activation. Nucleic Acids Res 48:12751–12777. doi:10.1093/nar/gkaa1124

Rosenkranz HS, Levy JA. 1965. Hydroxyurea: a specific inhibitor of deoxyribonucleic acid synthesis. Biochim Biophys Acta 95:181–183. doi:10.1016/0005-2787(65)90225-x

Savatier P, Lapillonne H, Jirmanova L, Vitelli L, Samarut J. 2002. Analysis of the cell cycle in mouse embryonic stem cells. Methods Mol Biol 185:27–33. doi:10.1385/1-59259-241-4:27

Schermelleh L, Haemmer A, Spada F, Rösing N, Meilinger D, Rothbauer U, Cardoso MC, Leonhardt H. 2007. Dynamics of Dnmt1 interaction with the replication machinery and its role in postreplicative maintenance of DNA methylation. Nucleic Acids Res 35:4301–4312. doi:10.1093/nar/gkm432

Sheu Y-J, Stillman B. 2010. The Dbf4-Cdc7 kinase promotes S phase by alleviating an inhibitory activity in Mcm4. Nature 463:113–117. doi:10.1038/nature08647

Spada F, Haemmer A, Kuch D, Rothbauer U, Schermelleh L, Kremmer E, Carell T, Längst G, Leonhardt H. 2007. DNMT1 but not its interaction with the replication machinery is required for maintenance of DNA methylation in human cells. J Cell Biol 176:565–571. doi:10.1083/jcb.200610062

Sporbert A, Domaing P, Leonhardt H, Cardoso MC. 2005. PCNA acts as a stationary loading platform for transiently interacting Okazaki fragment maturation proteins. Nucleic Acids Res 33:3521–3528. doi:10.1093/nar/gki665

Sporbert A, Gahl A, Ankerhold R, Leonhardt H, Cardoso MC. 2002. DNA polymerase clamp shows little turnover at established replication sites but sequential de novo assembly at adjacent origin clusters. Mol Cell 10:1355–1365. doi:10.1016/s1097-2765(02)00729-3

Stevens LC. 1973. A new inbred subline of mice (129-terSv) with a high incidence of spontaneous congenital testicular teratomas. J Natl Cancer Inst 50:235–242. doi:10.1093/jnci/50.1.235

Stillman B, Diffley JFX, Iwasa JH. 2025. Mechanisms for licensing origins of DNA replication in eukaryotic cells. Nat Struct Mol Biol 32:1143–1153. doi:10.1038/s41594-025-01587-5

Strzalka W, Ziemienowicz A. 2011. Proliferating cell nuclear antigen (PCNA): a key factor in DNA replication and cell cycle regulation. Ann Bot 107:1127–1140. doi:10.1093/aob/mcq243

Tanaka S, Hu SZ, Wang TS, Korn D. 1982. Preparation and preliminary characterization of monoclonal antibodies against human DNA polymerase alpha. J Biol Chem 257:8386–8390. doi:10.1016/S0021-9258(18)34343-6

Técher H, Koundrioukoff S, Nicolas A, Debatisse M. 2017. The impact of replication stress on replication dynamics and DNA damage in vertebrate cells. Nat Rev Genet 18:535–550. doi:10.1038/nrg.2017.46

Ubhi T, Brown GW. 2019. Exploiting DNA replication stress for cancer treatment. Cancer Res 79:1730–1739. doi:10.1158/0008-5472.CAN-18-3631

UniProt Consortium. 2023. UniProt: the universal protein knowledgebase in 2023. Nucleic Acids Res 51:D523–D531. doi:10.1093/nar/gkac1052

Woodward AM, Göhler T, Luciani MG, Oehlmann M, Ge X, Gartner A, Jackson DA, Blow JJ. 2006. Excess Mcm2-7 license dormant origins of replication that can be used under conditions of replicative stress. J Cell Biol 173:673–683. doi:10.1083/jcb.200602108

Xu J, Zhang Y. 2010. How significant is a protein structure similarity with TM-score = 0.5? Bioinformatics 26:889–895. doi:10.1093/bioinformatics/btq066

Yan S, Michael WM. 2009. TopBP1 and DNA polymerase-alpha directly recruit the 9-1-1 complex to stalled DNA replication forks. J Cell Biol 184:793–804. doi:10.1083/jcb.200810185

Yao NY, O’Donnell M. 2010. SnapShot: The replisome. Cell 141:1088, 1088.e1. doi:10.1016/j.cell.2010.05.042

Yin Y, Lee WTC, Gupta D, Xue H, Tonzi P, Borowiec JA, Huang TT, Modesti M, Rothenberg E. 2021. A basal-level activity of ATR links replication fork surveillance and stress response. Mol Cell 81:4243–4257.e6. doi:10.1016/j.molcel.2021.08.009

Yoo HY, Shevchenko Anna, Shevchenko Andrej, Dunphy WG. 2004. Mcm2 is a direct substrate of ATM and ATR during DNA damage and DNA replication checkpoint responses. J Biol Chem 279:53353–53364. doi:10.1074/jbc.M408026200

Yuan Z, Georgescu R, Li H, O’Donnell ME. 2023. Molecular choreography of primer synthesis by the eukaryotic Pol α-primase. Nat Commun 14:3697. doi:10.1038/s41467-023-39441-1

Zafar F, Nakagaw T. 2011. Regulation of minichromosome maintenance (MCM) helicase in response to replication stress In: Kusic-Tisma J, editor. Fundamental Aspects of DNA Replication. InTech. doi:10.5772/19284

Zegerman P, Diffley JFX. 2010. Checkpoint-dependent inhibition of DNA replication initiation by Sld3 and Dbf4 phosphorylation. Nature 467:474–478. doi:10.1038/nature09373

Zeman MK, Cimprich KA. 2014. Causes and consequences of replication stress. Nat Cell Biol 16:2–9. doi:10.1038/ncb2897

Zhang Y, Skolnick J. 2004. Scoring function for automated assessment of protein structure template quality. Proteins 57:702–710. doi:10.1002/prot.20264

Zou L, Cortez D, Elledge SJ. 2002. Regulation of ATR substrate selection by Rad17-dependent loading of Rad9 complexes onto chromatin. Genes Dev 16:198–208. doi:10.1101/gad.950302

