## Extended data 1 for "Dynamic anatomy of the replisome under stress"

**CRISPR genome-edited cell line generation**

mESC J1 were cultured in Dulbecco's modified Eagle's medium (DMEM) high glucose (Cat. No.: D6429, Sigma-Aldrich Chemie GmbH, Steinheim, Germany) supplemented with 16% fetal calf serum (FCS) (Cat. No.: FBS 12 A, Capricorn scientific, Germany), 1× non-essential amino acids (Cat. No.: M7145, Sigma-Aldrich Chemie GmbH, Steinheim, Germany), 1× penicillin/streptomycin (Pen/Strep) (Cat. No.: P4333, Sigma-Aldrich Chemie GmbH, Steinheim, Germany), 1× L-glutamine (Cat. No.: G7513, Sigma-Aldrich Chemie GmbH, Steinheim, Germany), 0.1 mM betamercaptoethanol (Cat. No.: 4227, Carl Roth, Karlsruhe, Germany), 1000 U/ml recombinant mouse LIF (Cat. No.: ESG1107, Merck, Steinheim, Germany) and 2i (PD032591 and CHIR99021 (Cat. Nos.: 1408 and 1386 respectively, Axon Medchem, Netherlands)) on gelatin-coated culture dishes (0.2% gelatin; Cat. No.:G2500, Sigma-Aldrich Chemie GmbH, Steinheim, Germany). Cells were split every 2-4 days.

The homologous recombination templates were generated by using genomic DNA as template DNA isolated from mouse embryonic stem cells J1. Briefly, for each CRISPR target, 500 bp right and left homology arms were amplified from genomic DNA using standard PCR procedures. The amplified homology arms were verified with Sanger sequencing to identify the correct genomic regions that were amplified. The homology arms together with the linker and GFP were then cloned into pc209 (pBluescript KS+) backbone vector for the generation of a homologous recombination template.

Transfection was performed by electroporation using the Thermo Fisher Neon transfection system. When using CRISPR-Cas9-mediated Homology-Directed Repair (HDR), approximately 1·10^6 cells were transfected in a volume of 100 µl using 2.5 µg of gRNA/Cas9 plasmids and 20 µg of the homologous recombination template. The cells were grown for 2 days in gelatin-coated culture plates containing ES medium with 10 µM L755507 (Cat. No.: SML1362, Merck, Steinheim, Germany) to promote homology-directed repair. The medium was then replaced with fresh ES medium containing 1 µg/ml puromycin for 48 hours to eliminate cells that are not transfected. The cells were cultured for 7-10 days until individual clones could be picked from plates seeded at low confluency. These clones were then cultured separately, and their fluorescence was examined periodically to narrow down the selection. The most promising clones were further characterized via Western blots, genomic PCR, and immunofluorescence staining using antibodies that targeted the tagged protein as well as GFP.

| Name | Species | Type | Fluorophore/Tag |
|---|---|---|---|
| ESCJ1 GFP PCNA C1 | Mus musculus | Embryonic stem cells | GFP PCNA (N-ter) |
| ESCJ1 RPA34 GFP C1 | Mus musculus | Embryonic stem cells | GFP RPA34 (C-ter) |
| ESCJ1 GFP RFC2 C1 | Mus musculus | Embryonic stem cells | GFP RFC2 (N-ter) |
| ESCJ1 MCM2 GFP C1 | Mus musculus | Embryonic stem cells | GFP MCM2 (C-ter) |
| ESCJ1 CDC45 GFP C1 | Mus musculus | Embryonic stem cells | GFP CDC45 (C-ter) |
| ESCJ1 GFP PolA1 C5 | Mus musculus | Embryonic stem cells | GFP PolA (N-ter) |
| ESCJ1 GFP PolD1 C10 | Mus musculus | Embryonic stem cells | GFP PolD1 (N-ter) |
| ESCJ1 GFP Ligase I C1 | Mus musculus | Embryonic stem cells | GFP Ligase I (N-ter) |
| ESCJ1 GFP PRIM1 C5 | Mus musculus | Embryonic stem cells | GFP PRIM1 (N-ter) |
| ESCJ1 GFP DNMT1 C13 | Mus musculus | Embryonic stem cells | GFP DNMT1 (N-ter) |

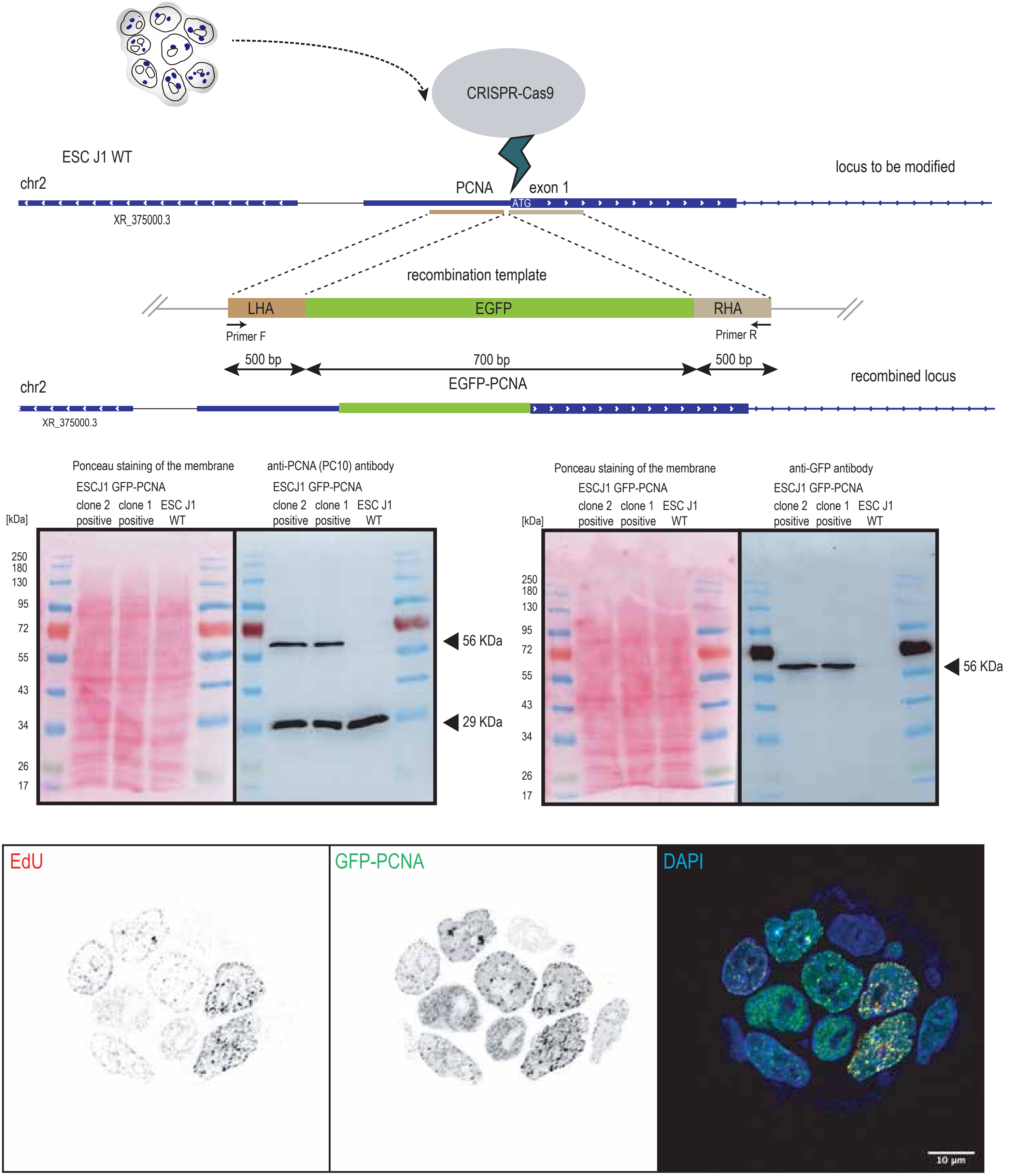

CRISPR-Cas9
ESC J1 WT
chr2
PCNA
exon 1
ATG
locus to be modified
XR_375000.3
recombination template
LHA
EGFP
RHA
Primer F
Primer R
500 bp
700 bp
500 bp
EGFP-PCNA
recombined locus
chr2
XR_375000.3
Ponceau staining of the membrane
anti-PCNA (PC10) antibody
ESCJ1 GFP-PCNA
clone 2 positive
clone 1 positive
ESC J1 WT
[kDa]
250
180
130
95
72
55
43
34
26
17
56 KDa
29 KDa
Ponceau staining of the membrane
anti-GFP antibody
ESCJ1 GFP-PCNA
clone 2 positive
clone 1 positive
ESC J1 WT
[kDa]
56 KDa
EdU
GFP-PCNA
DAPI
10 µm

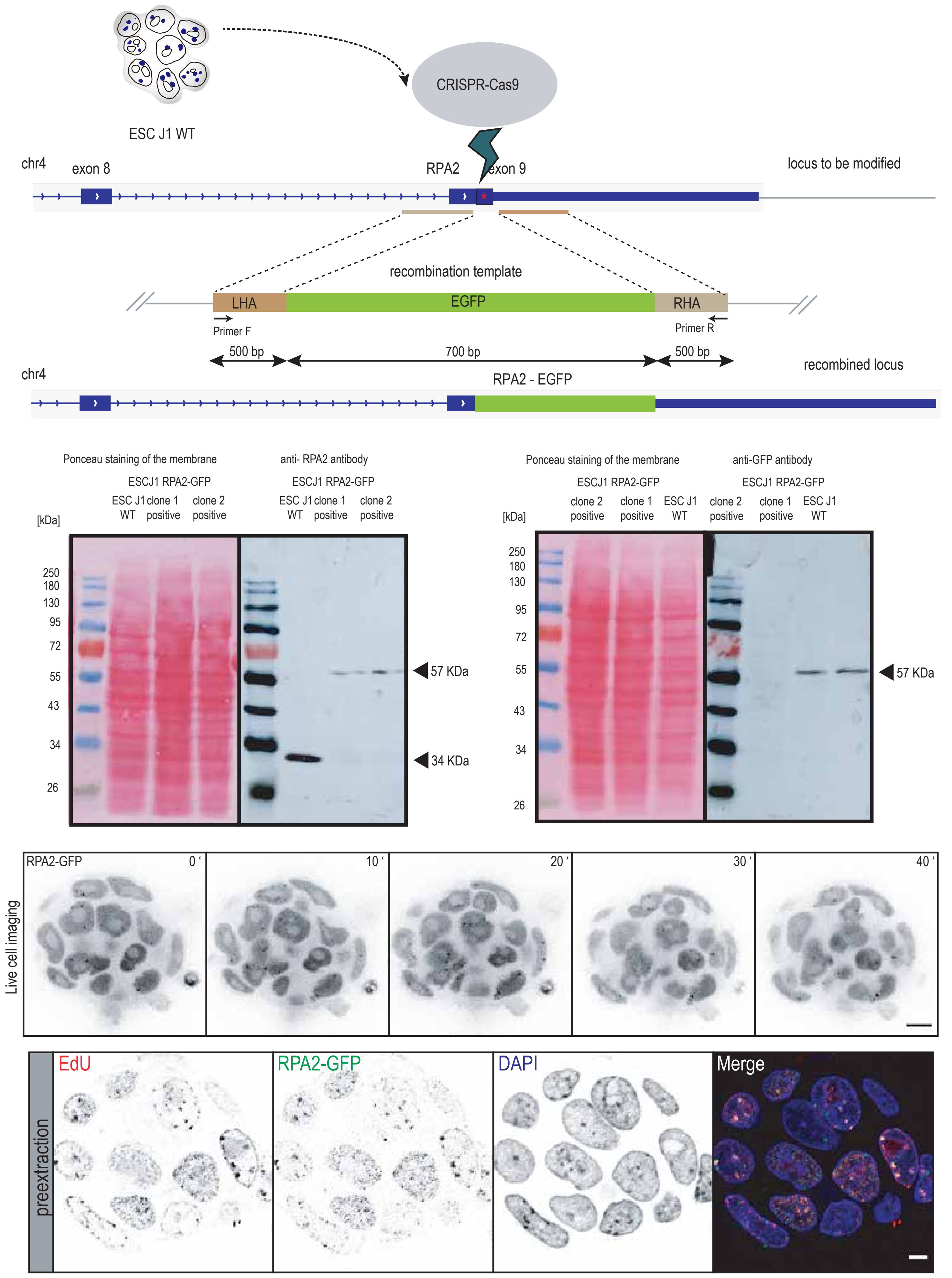

ESC J1 WT
CRISPR-Cas9
chr4
exon 8
RPA2
exon 9
locus to be modified
recombination template
LHA
EGFP
RHA
Primer F
Primer R
500 bp
700 bp
500 bp
recombined locus
chr4
RPA2 - EGFP
Ponceau staining of the membrane
anti- RPA2 antibody
ESCJ1 RPA2-GFP
ESC J1 WT
clone 1 positive
clone 2 positive
[kDa]
250
180
130
95
72
55
43
34
26
57 KDa
34 KDa
anti-GFP antibody
ESC J1 WT
57 KDa
Live cell imaging
RPA2-GFP
0 ‘
10 ‘
20 ‘
30 ‘
40 ‘
preextraction
EdU
RPA2-GFP
DAPI
Merge

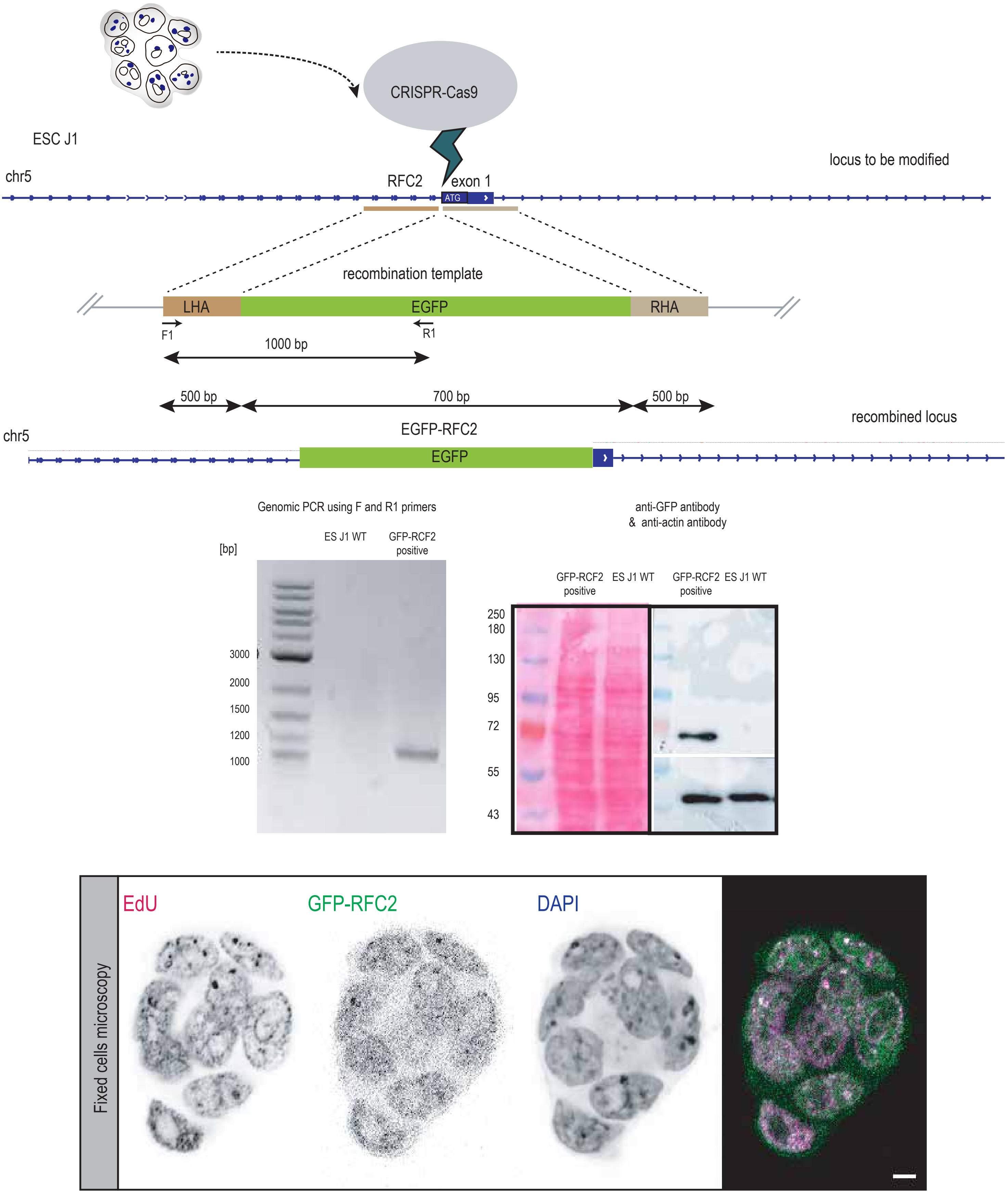
ESC J1
CRISPR-Cas9
locus to be modified
chr5
RFC2
exon 1
ATG
recombination template
LHA
EGFP
RHA
F1
R1
1000 bp
500 bp
700 bp
500 bp
recombined locus
EGFP-RFC2
chr5
EGFP
Genomic PCR using F and R1 primers
[bp]
ES J1 WT
GFP-RCF2 positive
3000
2000
1500
1200
1000
anti-GFP antibody
& anti-actin antibody
GFP-RCF2 positive
ES J1 WT
GFP-RCF2 positive
ES J1 WT
250
180
130
95
72
55
43
Fixed cells microscopy
EdU
GFP-RFC2
DAPI

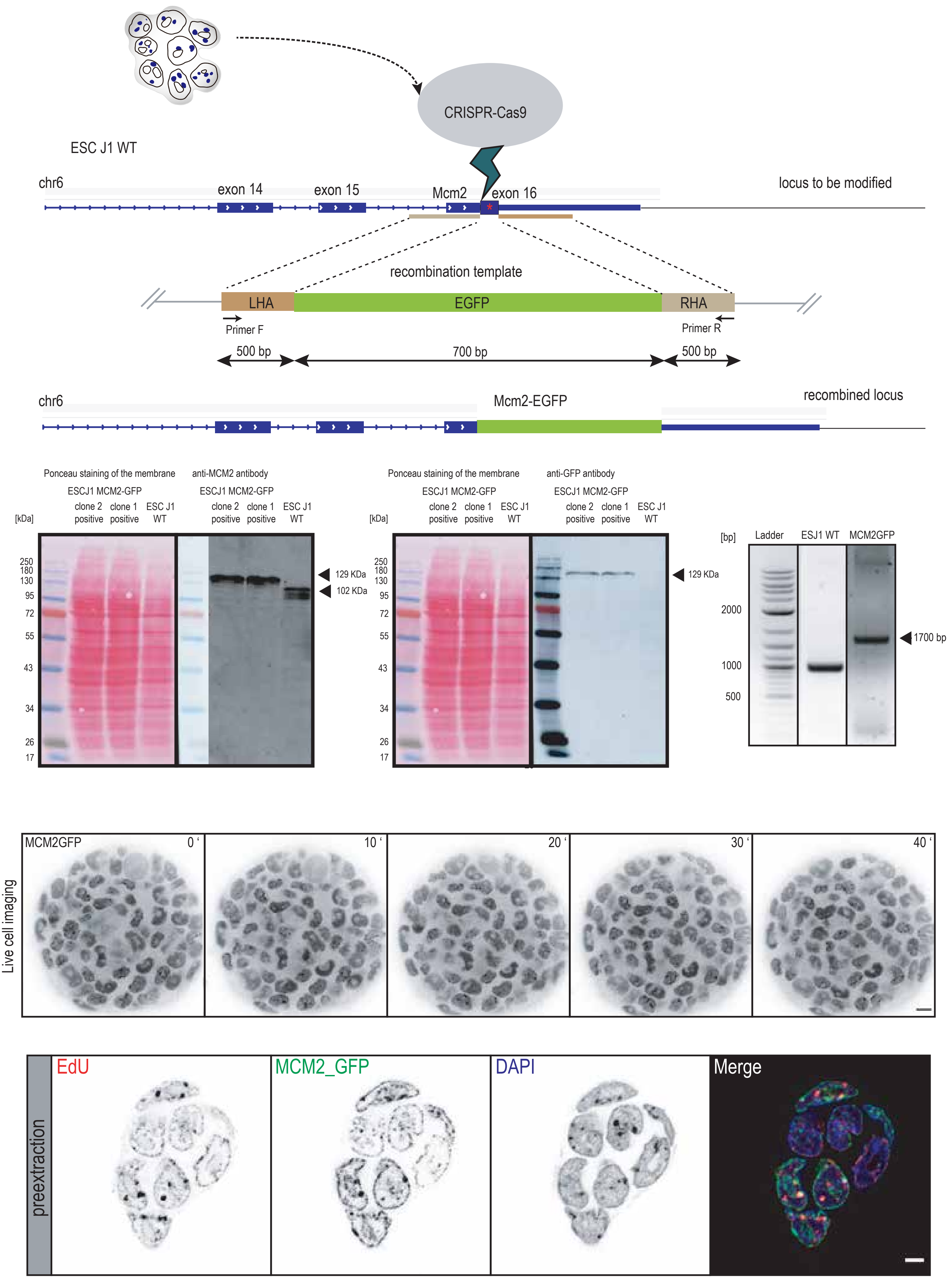

CRISPR-Cas9
ESC J1 WT
chr6
exon 14
exon 15
Mcm2
exon 16
locus to be modified
recombination template
LHA
EGFP
RHA
Primer F
Primer R
500 bp
700 bp
500 bp
chr6
Mcm2-EGFP
recombined locus
Ponceau staining of the membrane
anti-MCM2 antibody
ESCJ1 MCM2-GFP
clone 2 positive
clone 1 positive
ESC J1 WT
[kDa]
250
180
130
95
72
55
43
34
26
17
129 KDa
102 KDa
Ponceau staining of the membrane
anti-GFP antibody
ESCJ1 MCM2-GFP
clone 2 positive
clone 1 positive
ESC J1 WT
[kDa]
129 KDa
[bp]
Ladder
ESJ1 WT
MCM2GFP
2000
1000
500
1700 bp
Live cell imaging
MCM2GFP
0 ‘
10 ‘
20 ‘
30 ‘
40 ‘
preextraction
EdU
MCM2_GFP
DAPI
Merge

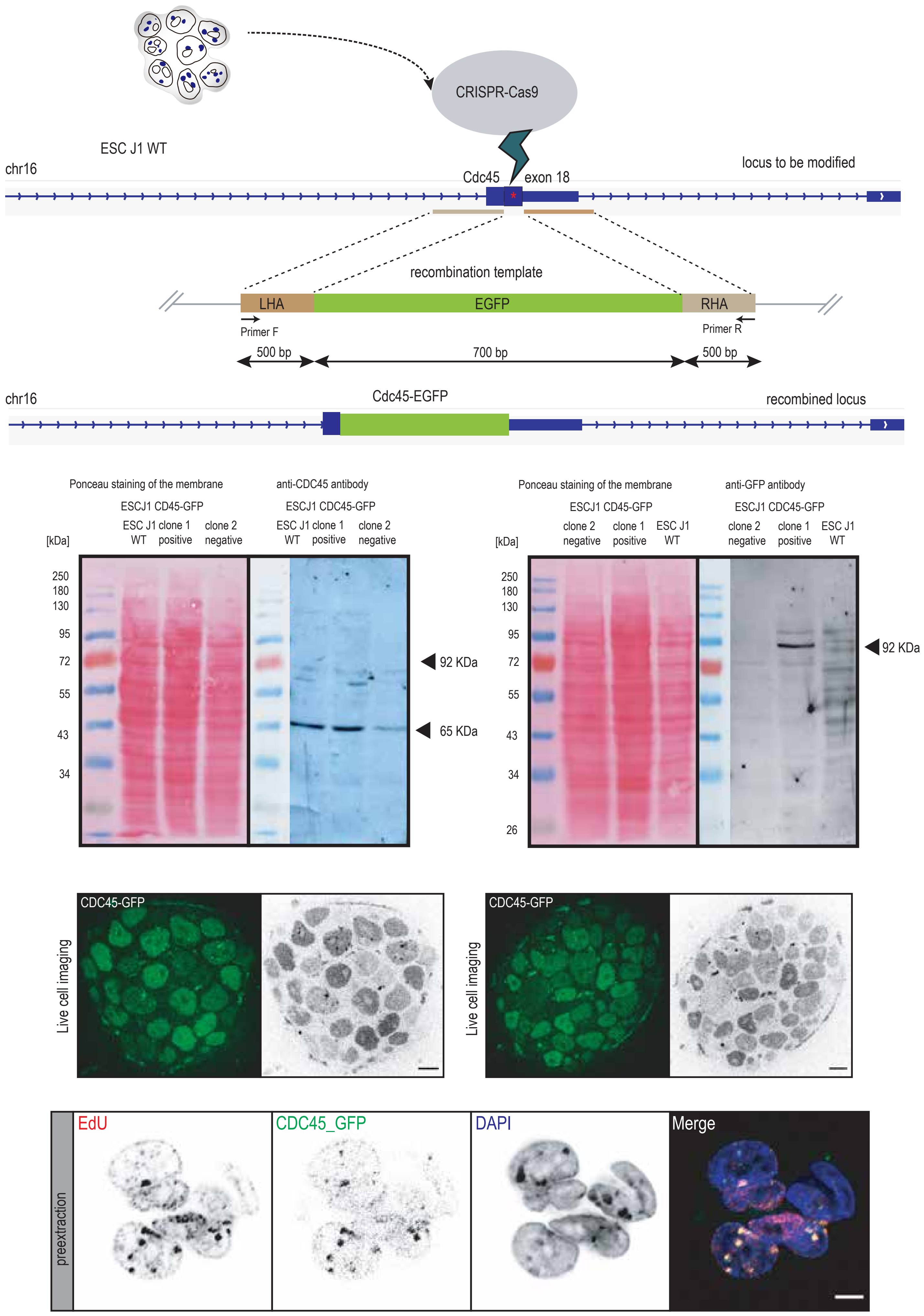

CRISPR-Cas9
ESC J1 WT
chr16
locus to be modified
Cdc45
exon 18
recombination template
LHA
EGFP
RHA
Primer F
Primer R
500 bp
700 bp
500 bp
chr16
Cdc45-EGFP
recombined locus
Ponceau staining of the membrane
anti-CDC45 antibody
ESCJ1 CD45-GFP
ESC J1 WT
clone 1 positive
clone 2 negative
[kDa]
250
180
130
95
72
55
43
34
92 KDa
65 KDa
Ponceau staining of the membrane
anti-GFP antibody
ESCJ1 CD45-GFP
clone 2 negative
clone 1 positive
ESC J1 WT
26
92 KDa
CDC45-GFP
Live cell imaging
CDC45-GFP
Live cell imaging
preextraction
EdU
CDC45_GFP
DAPI
Merge

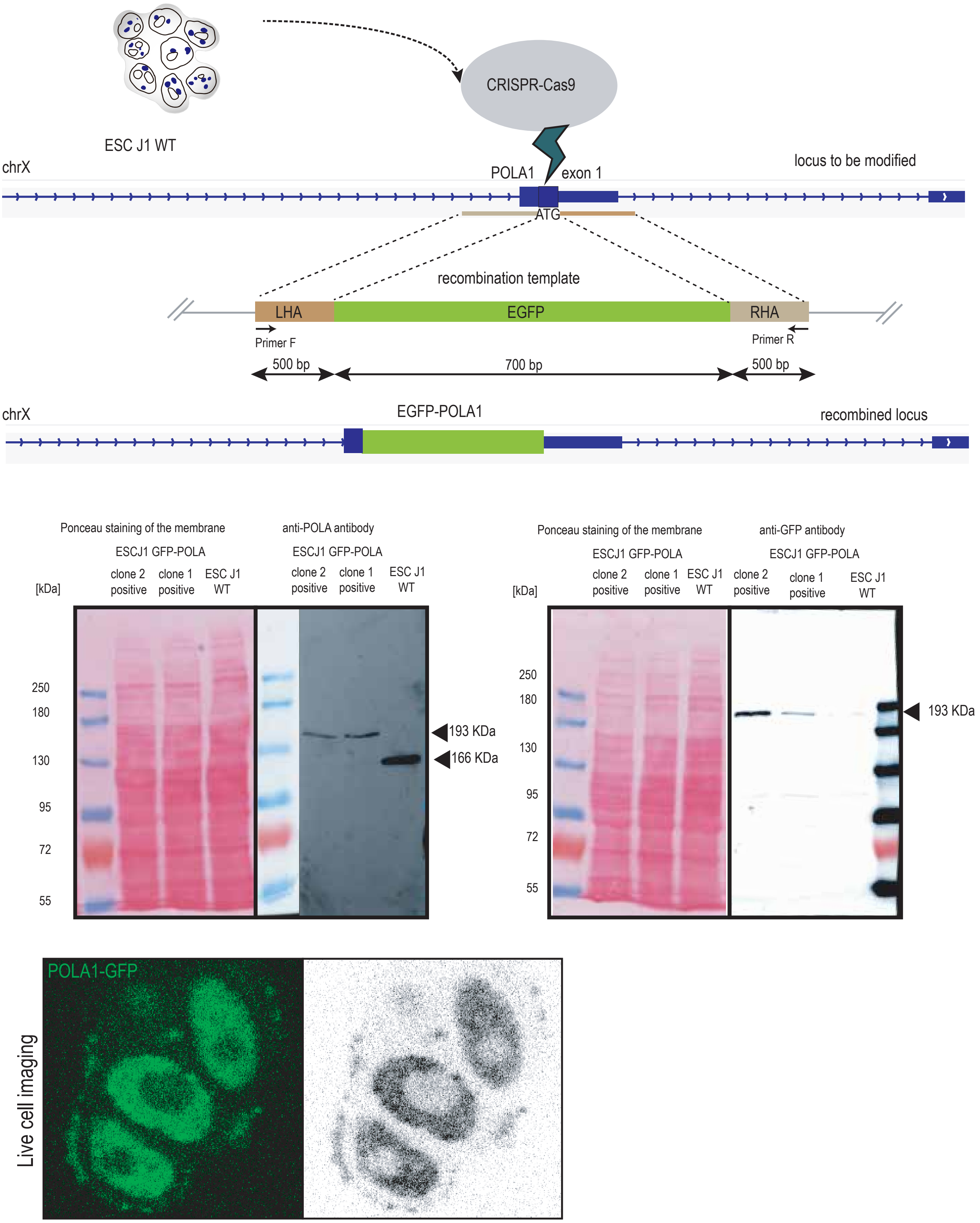

ESC J1 WT
CRISPR-Cas9
chrX
POLA1
exon 1
locus to be modified
ATG
recombination template
LHA
EGFP
RHA
Primer F
Primer R
500 bp
700 bp
500 bp
chrX
EGFP-POLA1
recombined locus
Ponceau staining of the membrane
anti-POLA antibody
ESCJ1 GFP-POLA
ESCJ1 GFP-POLA
[kDa]
clone 2 positive
clone 1 positive
ESC J1 WT
clone 2 positive
clone 1 positive
ESC J1 WT
250
180
130
95
72
55
193 KDa
166 KDa
Ponceau staining of the membrane
anti-GFP antibody
ESCJ1 GFP-POLA
ESCJ1 GFP-POLA
[kDa]
clone 2 positive
clone 1 positive
ESC J1 WT
clone 2 positive
clone 1 positive
ESC J1 WT
250
180
130
95
72
55
193 KDa
Live cell imaging
POLA1-GFP

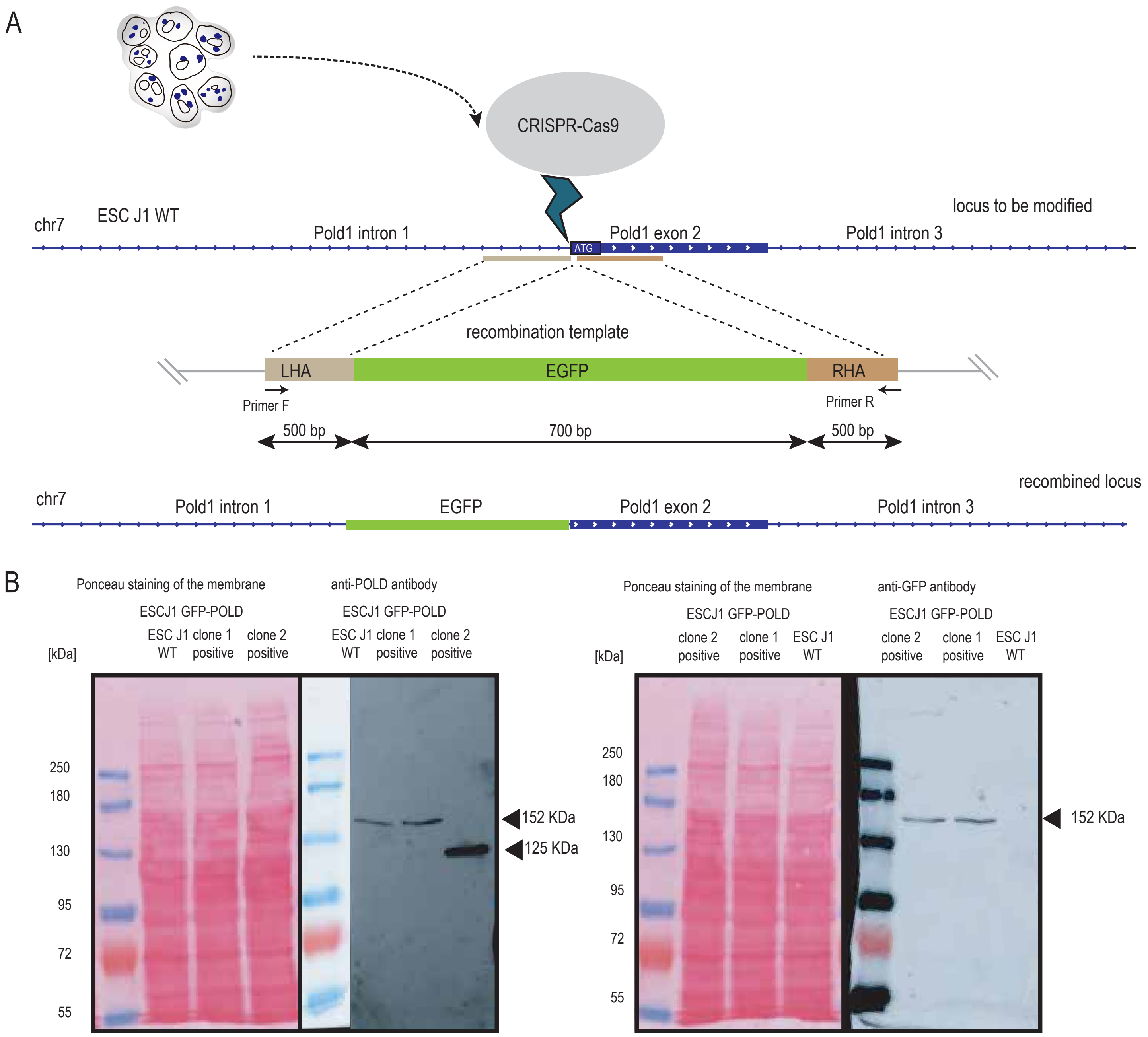
A
CRISPR-Cas9
chr7
ESC J1 WT
locus to be modified
Pold1 intron 1
ATG
Pold1 exon 2
Pold1 intron 3
recombination template
LHA
EGFP
RHA
Primer F
Primer R
500 bp
700 bp
500 bp
recombined locus
chr7
Pold1 intron 1
EGFP
Pold1 exon 2
Pold1 intron 3
B
Ponceau staining of the membrane
anti-POLD antibody
ESCJ1 GFP-POLD
ESCJ1 GFP-POLD
ESC J1 WT
clone 1 positive
clone 2 positive
ESC J1 WT
clone 1 positive
clone 2 positive
[kDa]
250
180
130
95
72
55
152 KDa
125 KDa
Ponceau staining of the membrane
anti-GFP antibody
ESCJ1 GFP-POLD
ESCJ1 GFP-POLD
clone 2 positive
clone 1 positive
ESC J1 WT
clone 2 positive
clone 1 positive
ESC J1 WT
[kDa]
250
180
130
95
72
55
152 KDa

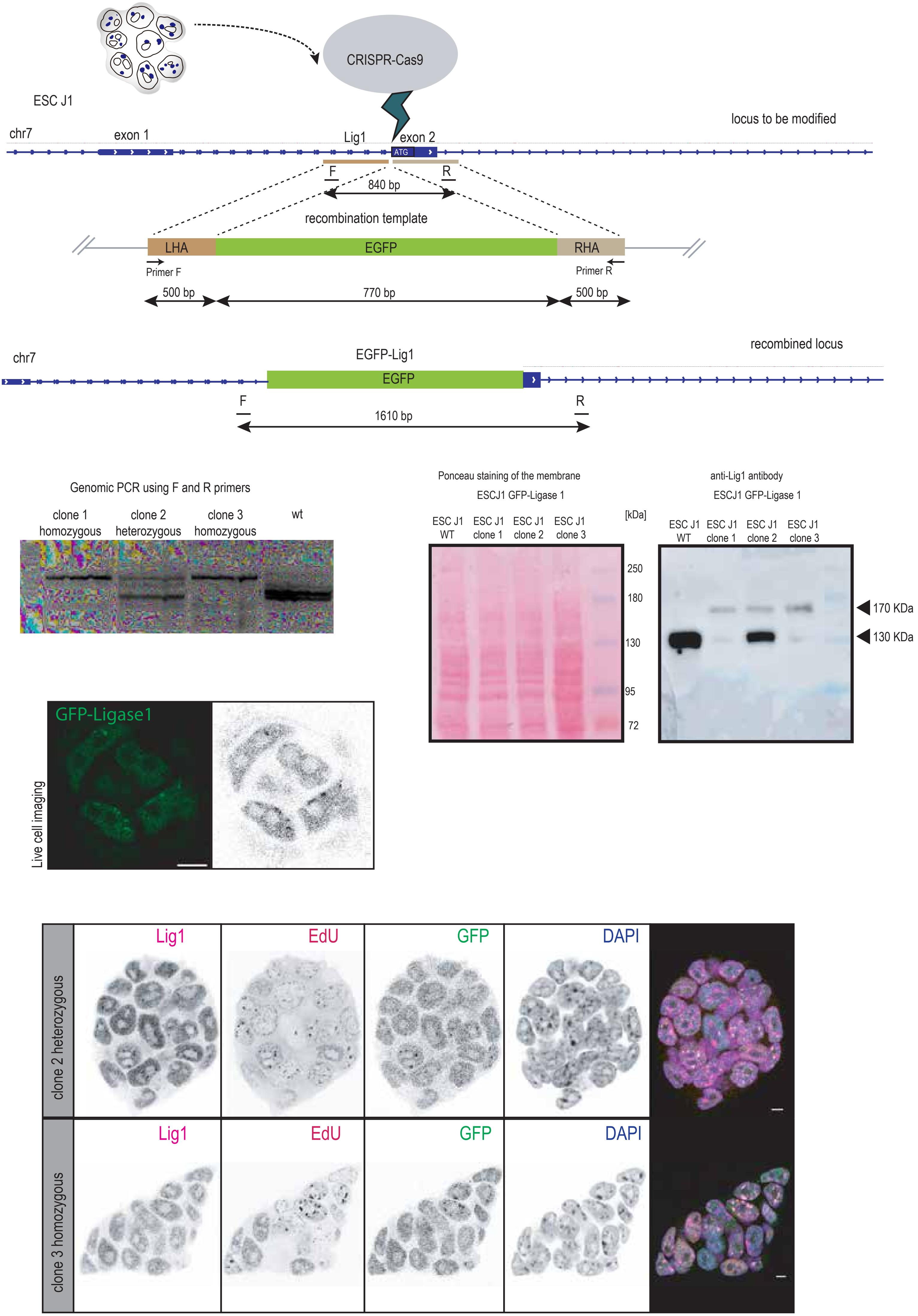
ESC J1
CRISPR-Cas9
locus to be modified
chr7
exon 1
Lig1
exon 2
ATG
F
R
840 bp
recombination template
LHA
EGFP
RHA
Primer F
Primer R
500 bp
770 bp
500 bp
recombined locus
chr7
EGFP-Lig1
EGFP
F
R
1610 bp
Genomic PCR using F and R primers
clone 1 homozygous
clone 2 heterozygous
clone 3 homozygous
wt
Ponceau staining of the membrane
ESCJ1 GFP-Ligase 1
ESC J1 WT
ESC J1 clone 1
ESC J1 clone 2
ESC J1 clone 3
[kDa]
250
180
130
95
72
anti-Lig1 antibody
ESCJ1 GFP-Ligase 1
ESC J1 WT
ESC J1 clone 1
ESC J1 clone 2
ESC J1 clone 3
170 KDa
130 KDa
GFP-Ligase1
Live cell imaging
clone 2 heterozygous
clone 3 homozygous
Lig1
EdU
GFP
DAPI

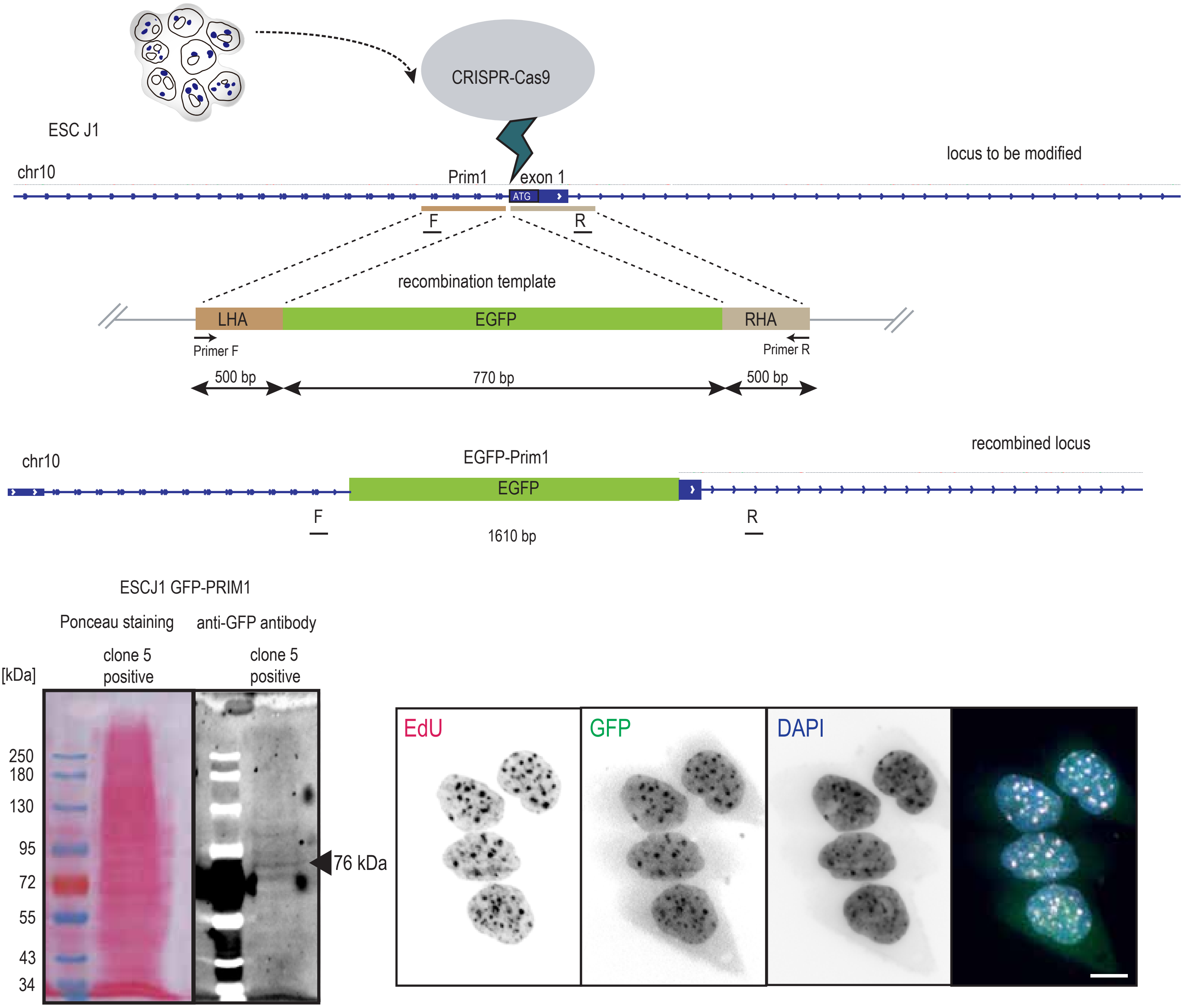
ESC J1
CRISPR-Cas9
locus to be modified
chr10
Prim1
exon 1
ATG
F
R
recombination template
LHA
EGFP
RHA
Primer F
Primer R
500 bp
770 bp
500 bp
recombined locus
chr10
EGFP-Prim1
EGFP
F
R
1610 bp
ESCJ1 GFP-PRIM1
Ponceau staining
anti-GFP antibody
clone 5 positive
clone 5 positive
[kDa]
250
180
130
95
72
55
43
34
76 kDa
EdU
GFP
DAPI
